# Lineage-resolved analysis of embryonic gene expression evolution in *C. elegans* and *C. briggsae*

**DOI:** 10.1101/2024.02.03.578695

**Authors:** Christopher R. L. Large, Rupa Khanal, LaDeana Hillier, Chau Huynh, Connor Kubo, Junhyong Kim, Robert H. Waterston, John I. Murray

## Abstract

What constraints govern the evolution of gene expression patterns across development remains a fundamental question of evolutionary biology. The advent of single-cell sequencing opens the possibility of learning these constraints by systematically profiling homologous cells across different organisms. The nematode *C. elegans* is a well-studied model for embryonic development, and its invariant lineage that is conserved with other *Caenorhabditis* species makes it an ideal model to directly compare gene expression between homologous progenitor and terminal cell types across evolution. We have measured the spatiotemporal divergence of gene expression across embryogenesis by collecting, annotating, and comparing the transcriptomes of homologous embryonic progenitors and terminal cell types, using a dataset comprising >200,000 *C. elegans* cells and >190,000 *C. briggsae* cells. We find a high level of similarity in gene expression programs between the species despite tens of millions of years of evolutionary divergence, consistent with their conserved developmental lineages. Even still, thousands of genes show divergence in their cell-type specific expression patterns, and these are enriched for categories involved in environmental response and behavior. Comparing the degree of expression conservation across cell types reveals that certain cell types such as neurons, have diverged more than others such as the intestine and body wall muscle. Taken together, this work identifies likely constraints on the evolution of developmental gene expression and provides a powerful resource for addressing diverse evolutionary questions.

## Introduction

Diverse animal species have a set of common cell types employed in a wide diversity of body plans that have persisted over hundreds of millions of years of evolution, as well as specialized cell types appropriate to their unique needs. We can easily recognize many ancient cell types, such as neurons, muscles, and epidermal cells, as homologous based on morphology and function. Whether these cells adopt similar or different functions ultimately results from maintenance or differences in gene expression, and determining how cell type expression programs evolve based on form, function, and evolutionary history is still an active area of exploration.

Evolution of development can involve many factors including changes in gene expression within homologous cells that may or may not alter cellular function, changes in developmental timing or changes in cell number or position. At sufficient evolutionary distances, conserved expression patterns are predictive of conserved functional roles [e.g. (van Noort *et al*. 2003)], and changes in gene expression are thought to be a major mechanism underlying organismal evolution [e.g. (King and Wilson 1975)]. However, comparisons between related species have found that cellular phenotype can be conserved despite significant changes in the underlying gene expression and regulatory networks (Barrière and Ruvinsky 2014). This “Developmental System Drift” can occur by multiple mechanisms including changes in the expression or function of existing genes, or gene duplication events followed by sub- or neo-functionalization (True and Haag 2001; Haag and True 2021). However, the full extent, shape, and evolutionary basis of the constraint on these developmental changes remain poorly resolved. Thus, defining the extent to which genome-wide expression is conserved in homologous cell types across evolution could both help to define the fundamental conserved features of each cell type and to understand how cells adaptively evolve for organismal function.

Single-cell genomics approaches have made it possible to measure gene expression at genome scale across cell states in developing embryos and differentiated tissues. Comparing single-cell gene expression data across species can identify common and diverse cell states, changes in cell type abundance, and gene expression differences between species (Shafer *et al*. 2022; Murat *et al*. 2023). However, comparing cellular transcriptomes across an entire animal’s set of cell types can be challenging for large or complex organisms. *Caenorhabditis* embryos provide an ideal system for the study of the evolution of homologous cell types and their specification at single-cell resolution. *C. elegans* embryos produce 558 terminal cells through an invariant and fully described sequence of cell divisions (Sulston *et al*. 1983). This lineage is conserved nearly perfectly across related species including *Caenorhabditis briggsae* (Zhao *et al*. 2008; Memar *et al*. 2019), despite the *C. elegans* and *C. briggsae* genomes differing by more than 1.6 substitution per neutral site (Stein *et al*. 2003). Embryonic gene expression in *C. elegans* is well characterized, with several single-cell resolution imaging and RNA-sequencing studies having revealed the transcriptomes and transcription factor (TF) protein expression profiles of nearly all progenitor lineage states and embryonic terminal cell types (Murray *et al*. 2012; Tintori *et al*. 2016; Packer *et al*. 2019; Ma *et al*. 2021; Cole *et al*. 2023). In most single-cell studies, cell types are defined using *ad hoc* markers, a practice that is still controversial and actively debated (Bard *et al*. 2005; Arendt *et al*. 2016; Morris 2019; Xia and Yanai 2019; Domcke and Shendure 2023)*. Caenorhabditis species* are unique among major models in providing a known enumerated list of cell states that can be used to validate the completeness and accuracy of annotations.

Here, we compare the gene expression of progenitor and terminal cell types in *C. elegans* and *C. briggsae* embryos by single-cell RNA sequencing of ∼200,000 cells per species (>150x coverage of the lineage) comprising ∼487 shared progenitor and terminal cell states. We find that a gene’s expression conservation is correlated with gene function, essentiality, overall expression level, and expression breadth across the animal. Comparing expression divergence between cell types identifies higher expression conservation in germline, intestinal, muscle, and hypodermal cell types with increased divergence of the neuronal and some specialized mesodermal cell types. Cell types also vary in their expression of species-specific genes or genes with complex (non 1:1) orthology relationships, with an especially high frequency of cell type-specific genes being from these rapidly evolving categories in early embryonic progenitors and the amphid sensory neurons. Overall, this dataset provides a powerful resource for studies into the evolution of developmental gene expression.

## Results

### Aligned single-cell transcriptome atlases of C. elegans and C. briggsae

We measured mRNA levels for embryonic cells from the *C. briggsae* AF16 strain by single-cell RNA-seq, using the 10x genomics platform. We prepared loosely synchronized embryo populations (detailed in methods) with the goal of maximizing representation of embryonic stages from gastrulation (∼28-cell stage) through terminal differentiation. After removing doublets and low-quality cells, this resulted in a *C. briggsae* dataset with 193,020 cells collected across seven independent cell preparations (∼157x average coverage of the 558 terminal cells and 670 internal lineage branches in the shared *C. elegans/C. briggsae* lineage). We compared these cells to the existing *C. elegans* single-cell embryonic atlas (Packer *et al*. 2019), supplemented with an additional 27,327 wild-type cells derived from late-stage embryos. We also included an additional 139,557 cells from three *C. elegans* TF mutants for which the TF was expressed only late or in specific cell types and for which we detected very limited expression changes (see methods). This gave a total of 255,027 *C. elegans* cells (∼207x coverage).

To compare between *C. elegans* and *C. briggsae*, we focused primarily on a set of 12,803 genes, which we refer to for simplicity as the 1:1 orthologous gene set. These were identified by combining previous orthology sets with gene synteny and reciprocal BLAST searches (Supp. Figure 1, see methods for details). This approach resulted in 1,683 additional orthologs not identified by previous methods. We estimated the embryonic stage of each *C. briggsae* cell by comparing the expression of 1:1 orthologous genes to a whole-organism *C. elegans* time course. This method agreed well with the expected stages based on the aging of embryos in each prep (Supp. Figure 2-4), and with the expected ages of cells with specific lineage annotations. Overall, both species’ datasets contained a similar range of cell stages, with slightly more early cells in the *C. briggsae* dataset due to the inclusion of more early-targeted samples in that dataset (Supp. Figure 2 and 5).

Despite the near perfect conservation of the lineage identity, we did not know if the cell fate identity of the two species would be similarly conserved. Therefore, we attempted to identify homologous cell types in a joint transcriptional space using the 1:1 orthologous gene set. We integrated the *C. elegans* and *C. briggsae* data sets by co-embedding with Canonical Correlation Analysis (CCA) (Butler *et al*. 2018; Stuart *et al*. 2019), yielding a single integrated dataset (Figure 1A-B). Clusters identified in this shared space generally were evenly composed of cells from both species (Figure 1C,D), were well-mixed in subsets selected by embryo time windows or annotated cell classes (such as mesoderm, epidermis, neurons), and cells from each species generally showed similar expression of cell type markers known from *C. elegans* (Supp. Figure 6-12). We manually annotated terminal cell types in both species, using the previous cell type annotations from (Packer *et al*. 2019), the cell class-specific co-embeddings of *C. elegans* and *C. briggsae* cells, and the expression of known marker genes from *C. elegans*. Comparisons between the cell type annotations for the terminal cell types in (Packer et al. 2019) with the jointly annotated cell types reveals high overlap, with minor deviations due to technical differences in annotation (Supp. Table 1). The high concordance of marker expression across species indicates that these annotations presumably represent homologous cells.

**Figure 1.**
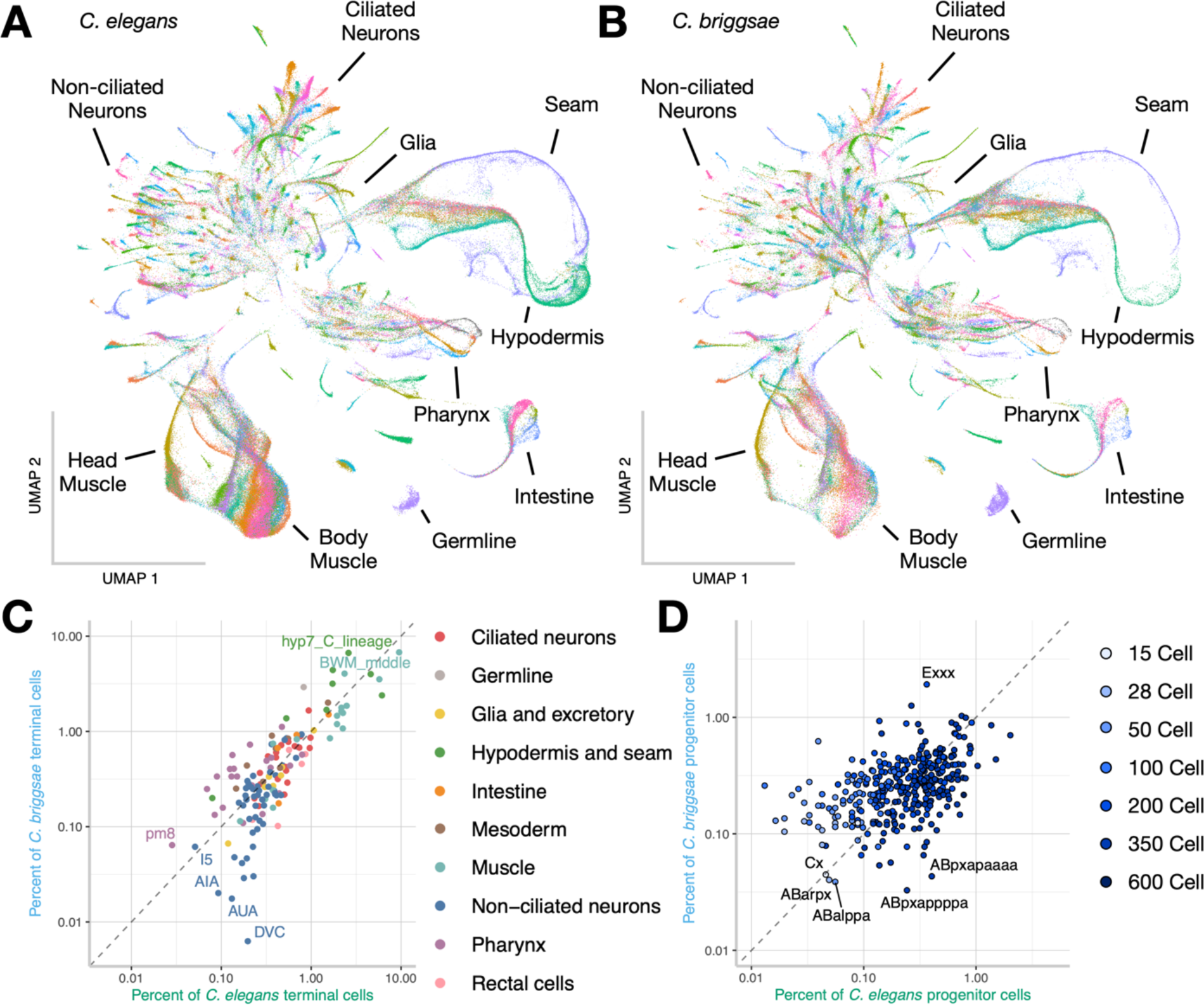
Homology in annotation of cell types between *C. elegans* and *C. briggsae*. A, B) Using the orthologous genes between the species, the datasets were co-embedded in a shared reduced dimension space. The terminal and progenitor cell type annotations for *C. elegans* (A) and *C. briggsae* (B) are shown with color labeling matching cell type identities between the species. C) The percentage of terminal cell types for each cell type was similar overall, but biased towards *C. elegans* for some non-ciliated neurons. D) The percentage of progenitor cells for each progenitor type was also similar, with a slight bias towards cells from younger embryos in *C. briggsae*.

Almost all terminal cell types (corresponding to 496 out of a total 558 terminal cells as well as the parents of 54 additional terminal cells) were found reliably in both species with similar frequencies (Figure 1C). Only nine cell types were largely not detected in *C. briggsae*, but detected in *C. elegans* (i.e. <10% of cells in the cluster were from *C. briggsae*); for example the DVC neuron cluster contained 8 *C. briggsae* cells and 313 *C. elegans* cells (Figure 1C). These relative depletions could reflect either technical or minor biological differences and were not considered in subsequent comparative analyses. Other known terminal cell types not found in *C. briggsae*, were also not found or poorly represented in the *C. elegans* embryonic cell atlas, likely due to biases in their amenability to dissociation (e.g. pharyngeal neurons) or a lack of clear cell type markers (see Supp. Table 2, Supp. Figure 13).

We similarly annotated the progenitor cell types from *C. briggsae* using markers with known lineage-specific expression in *C. elegans*. We used multiple markers per lineage branch, and confirmed that lineages often formed contiguous trajectories in reduced-dimensional space (Supp. Table 3; Supp. Figure 14-17). We observed similar proportions of cells with each lineage annotation in *C. elegans* and *C. briggsae*, with more early cells in *C. briggsae*, as expected due to this dataset being slightly more early-biased (Figure 1D; Supp. Figure 2). Comparing our observations to previous studies investigating the expression pattern of specific homologous genes between *C. elegans* and *C. briggsae*, we found agreement for 9 of 10 genes, supporting the accuracy of the cell type annotations (Maduro and Pilgrim 1996; Dufourcq *et al*. 1999; Aamodt *et al*. 2000; Marshall and McGhee 2001; Wang and Chamberlin 2002; Wang *et al*. 2004; Nayak *et al*. 2005; Konwerski *et al*. 2005; Barrière *et al*. 2011, 2012; Beadell *et al*. 2011; Chai *et al*. 2022); Supp. Table 4). Overall, we find that most *C. elegans* and *C. briggsae* cells have easily recognizable, homologous, molecular states. This provides a basis for comparative analysis of homologous cells and genes across development.

### Gene-level constraints vary by breadth of expression and functional class

We used the two species’ expression atlases to compare the expression patterns of each gene across cell types reliably detected in both species. For each gene, we calculated metrics of expression pattern conservation and expression specificity across species. To measure expression pattern conservation of each gene between *C. elegans* and *C. briggsae*, we used the Jensen-Shannon Distance (JSD_gene_), calculated on the expression levels (Transcript per Million (TPM)) in all homologous terminal cell types in pseudo-bulk. JSD_gene_ values range between 0 and 1, with lower JSD_gene_ values indicating conserved expression. To measure the degree of cell-type specificity of expression for each gene, we used the *Tau* metric (Yanai *et al*. 2005). A low *Tau* corresponds to broad expression, while a high *Tau* indicates patterned or cell type specific expression. We focus here on genes that were confidently detected (TPM > 100 in at least one cell type) in both species. This excludes lowly-expressed genes that may represent real differences but for which it is difficult to rule out technical artifacts. Supp. Figure 18 compares JSD_gene_ with other metrics and all gene statistics calculated on terminal cell types are available in Supp. Table 5.

Comparing expression across the terminal cell-types, we found several patterns of gene expression conservation:

- **Broadly expressed and conserved in expression pattern:** For example, the trithorax-related gene, *lin-59* has a JSD_gene_ of 0.23 and *Tau* of 0.16 (Supp. Figure 19).
- **Cell-type specific or patterned expression and conserved:** For example, *pha-4*/FoxA (JSD_gene_ = 0.29, *Tau* = 0.53), a well-studied TF required for pharynx specification, is consistently expressed in all pharyngeal cell types in both species (Figure 2A). Separately, *hlh-4* (JSD = 0.24, *Tau* = 0.85) is highly conserved with high expression exclusively in the ADL neuron in both species (Supp. Figure 20).
- **Conserved with quantitative changes:** Some genes, despite having conserved expression patterns, show substantial (>2.8 fold) changes in the absolute value of expression across cell types. For example, the Rab GTPase, *rab-7* (JSD_gene_ = 0.14) was detected in all cells of both species but at 3-fold higher levels in *C. briggsae* than in *C. elegans* (Supp. Figure 21). While such quantitative differences could arise for technical reasons such as errors in gene models or orthology assignments, manual inspection of their gene models, orthology relationships, and an independent embryonic bulk-RNA-seq, which shows similar species-specific enrichment (log2 difference > 1.5), supports the difference as biological.
- **Genes with poor or no expression conservation:** Genes vary continuously in JSD and other metrics (e.g. Figure 2F,H), emphasizing that conservation is not broken into discrete categories. For example, the AP-2 transcription factor, *aptf-2* (JSD_gene_ = 0.79) has broad and weak expression in both species, but in *C. briggsae* is also expressed at high-level in head body wall muscle and coelomocyte cells (Figure 2B), while the unstudied transmembrane gene, *T04A6.1* (JSD_gene_ = 0.98) is specifically expressed in the sensory neurons ADL, ADF and ASH in *C. briggsae* and in the inter/motor neuron RMG in *C. elegans* (Figure 2C).

**Figure 2.**
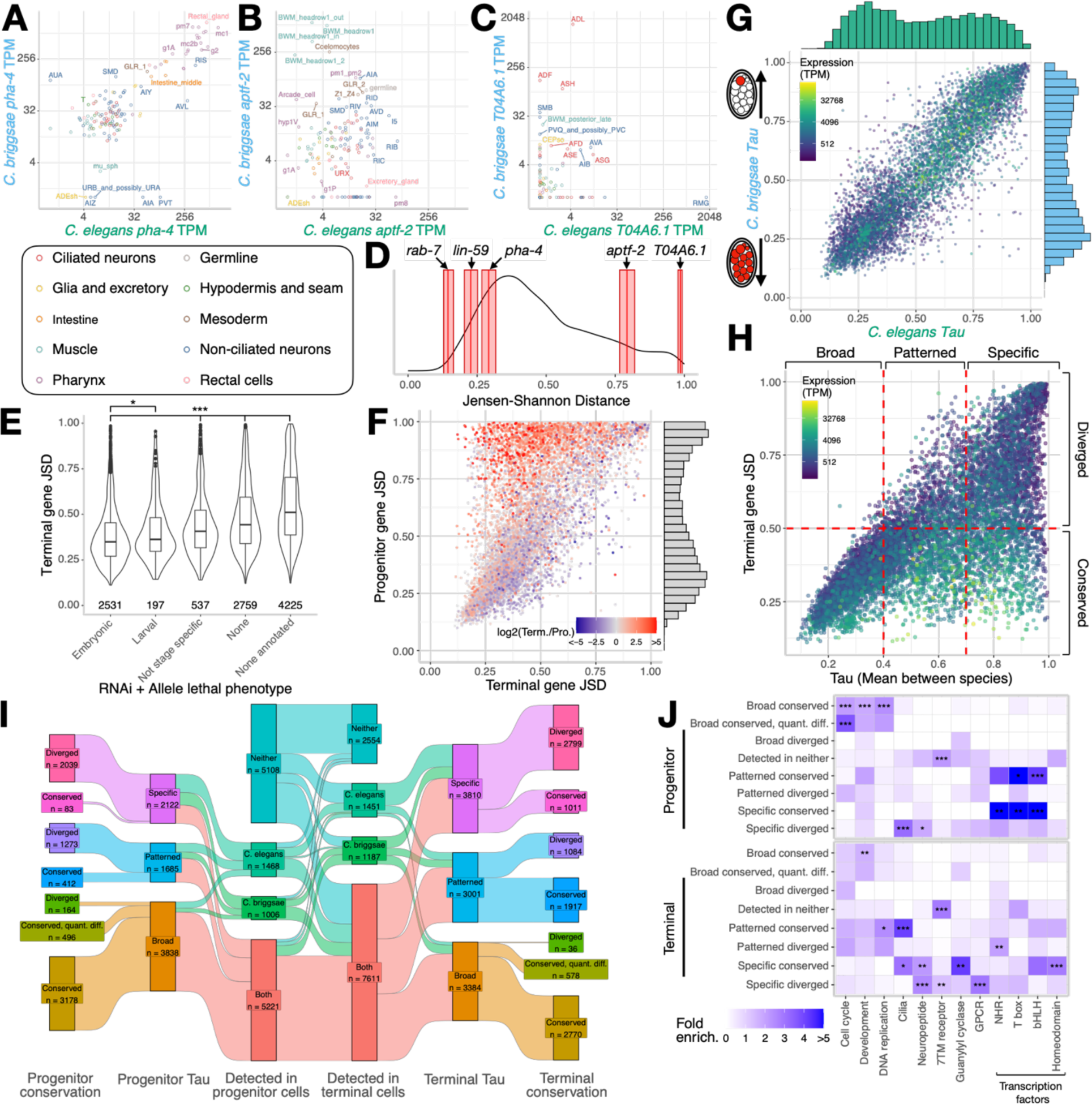
Orthologous genes between *C. elegans* and *C. briggsae* are mostly conserved in their gene expression patterns across embryonic development. A) The gene expression of *pha-4*, B) *aptf-2*, and C) *T04A6.1* for *C. elegans* and *C. briggsae* show a diversity of conservation patterns across the terminal cell types. D) The distribution of JSD_gene_ values for all well-detected genes (>100 TPM in at least one cell type) indicates that most genes have a conserved expression pattern. Each example gene is shown with a 95% confidence interval (red box) with the median value displayed as a dark red line. E) The JSD_gene_ in terminal cell types for genes with a *C. elegans* embryonic RNAi or allele lethal phenotype is lower by a Wilcoxon Rank Sum test (* is p <0.05, *** is p <0.0005). F) Comparison between the JSD_gene_ calculated on expression in the progenitors and terminal cell types for all genes that are well detected in both the progenitor or the terminal cell types. G) The *Tau* values from either species calculated on the terminal cell types for all well detected genes, with the maximum gene expression between either species. H) The mean of the *Tau* values from either species, calculated on the terminal cell types compared to the terminal JSD_gene_ with the maximum gene expression. I) A Sankey plot, showing the count and proportion of genes falling into the listed categories. J) Fold enrichment of the WormCat tier 1 categories within the outline conservation and gene expression breadth categories for the progenitor and terminal cell types. The significance of the enrichment is from a Bonferroni corrected Fisher’s exact test (* is p <0.05, ** is p <0.005, *** is p <0.0005).

Overall, most genes showed a conserved expression pattern (median JSD_gene_ = 0.44), while a smaller number showed higher JSD_gene_ values, corresponding to genes with expression changes between the species (Figure 2D). The relatively similar JSD_gene_ values of *rab-7* and *lin-59*, despite *rab-7* differing between species in overall level, highlights that this metric captures divergence in pattern, not level.

We would expect that genes essential for development should be more likely to have conserved expression patterns. Consistent with this, we found that genes that have an embryonic RNAi or mutant allele lethal phenotype in *C. elegans* have on average a lower gene expression pattern divergence than those that do not have an embryonic phenotype (Figure 2E). This suggests that the similarity measured by JSD_gene_ reflects biological constraints.

We asked how gene expression pattern conservation compares between terminal and progenitor cell types (Figure 2F). Since genes that are not well detected in either terminal or progenitor cell types will have a high JSD_gene_ due to measurement noise, we focused on genes that were well detected in both stages (>100 TPM in at least one cell type). A majority of genes showed good agreement in their JSD_gene_ between terminal and progenitor cell types. However, a subset of genes showed conserved expression patterns in terminal cell types, with more divergent patterns in the progenitor cell types. Most of these genes had greater expression in the terminal cell types. On average, genes with higher maximum expression in at least one cell type were more likely to have a conserved pattern (JSD_gene_ < 0.5) than genes with lower expression (Wilcoxon-rank sum test, p-value < 2.2e-16). By contrast, there were few examples of genes with similar expression in progenitors and divergent expression in terminal cells, partly due to the smaller number of genes detected at higher levels in progenitors (Figure 2F).

### Gene expression breadth remains consistent between species

We next assessed how well the breadth of expression (*Tau*) is conserved and how breadth relates to overall expression conservation. *Tau* shows a bimodal distribution, peaking in the lower range with many broadly-expressed genes, such as *rab-7* and *lin-59*, and in the upper range with specifically-expressed genes (*hlh-4* and T04A6.1; Figure 2G). Between these two peaks were genes corresponding to patterned expression across multiple cell types (*e.g. pha-4* and *aptf-2*). *Tau* was highly conserved between species (R^2^ = 0.7714), indicating that genes generally do not rapidly change in their expression breadth. Only 100 genes had a difference in *Tau* of greater than 0.3. Some of the divergent genes could be explained by incorrect orthology assignments or errors in gene models, but at least 62 appear to represent differences in either pattern, timing or level of expression (Supp. Table 6).

We found a modest linear relationship between the breadth of expression (*Tau*) and expression pattern conservation (JSD_gene_) (R^2^ of 0.53; Figure 2H). As expected, broadly detected (*Tau* < 0.4) genes generally show high conservation. For the genes that showed either patterned (0.4 < *Tau* < 0.7) or specific (Tau > 0.7) breadth of expression, we find a greater variance in the degree of conservation, with some genes showing either conserved or divergent expression.

### Gene classification reveals conservation of most transcription factors, and divergence of genes with neuronal function

To summarize general trends in which types of genes and patterns tend to have conserved or divergent expression, we classified genes based on their JSD_gene_ and *Tau* values (Figure 2H). Based on manual inspection of numerous genes (Supp. Figure 22), we defined genes as “conserved” if JSD_gene_ was less than 0.5 or “divergent” for higher JSD values. Similarly we used *Tau* to define genes as having “broad” (*Tau* < 0.4), “patterned” (0.4 <= *Tau* <= 0.7) or “specific” (*Tau* > 0.7) expression. While these thresholds are somewhat arbitrary, this classification makes it possible to visualize and understand major trends in gene expression conservation.

We found progenitors had more broadly expressed genes and fewer patterned or specifically expressed genes than terminal cells for each species (Figure 2I). For each *Tau* category, genes with higher maximum expression in at least one cell type were more likely to have conserved expression (as measured by JSD_gene_; Figure 2H; Supp. Figure 23). Progenitors and terminal cells differed in the conservation of patterned and specific gene expression, with substantially fewer conserved specific or patterned genes in the progenitors versus terminal cell types (Figure 2I). Between the species we found a similar proportion of broadly expressed genes that had similar patterns but quantitative differences (>2.8-fold difference in mean expression) in both the terminal (17%) and progenitor (13%) cell types (Supp. Figure 24-25).

We asked whether genes with conserved or divergent expression in each breadth (*Tau*) category are enriched for WormCat functional annotations (Figure 2J; (Yanai *et al*. 2005; Holdorf *et al*. 2020)). In progenitors, gene categories enriched in the broad-conserved category include cell cycle, development, and DNA replication, consistent with expectations for a rapidly dividing embryo. Genes annotated as transcription factors are enriched in both the specific-conserved and -divergent categories in progenitors. Specific transcription factor DNA binding domains especially bHLH, homeodomain and T-box families were enriched in the specific-conserved category, consistent with a conserved role of these families in development and fate specification. Conversely, the Nuclear Hormone Receptor (NHR) family is enriched in the specific-divergent category, consistent with the observation that the NHRs have rapidly expanded and diverged in nematodes (Antebi 2006).

Genes involved in the cilial program have differing conservation in progenitors vs terminal cells, being specific-diverged in the progenitors and patterned- or specific-conserved in the terminal cell types. Investigating expression of genes in the cilia program suggests this may be due to changes in the onset time of the cilial program, with *C. elegans* expressing cilial genes slightly earlier than *C. briggsae* (Supp. Figure 26). As a result, the same terminal cells express cilial genes in each species, but they are more reliably detected in ciliary neuron progenitors in *C. elegans* than in *C. briggsae*. This provides an interesting example of how genes can differ across evolution in expression timing in addition to pattern and level.

Within the terminal cell types, homeodomain transcription factors remain in the specific-conserved category, consistent with their importance in specification of terminal cell fates for neurons (Hobert 2021) and other cell types. Genes annotated with neuronal functions are enriched in both the specific-diverged and -conserved categories. Examination of more specific annotations revealed that cilial genes and guanylyl cyclases were enriched in the conserved categories, while neuropeptides were enriched in both conserved and divergent categories. Finally, the chemoreceptor-associated G-protein coupled receptor (GPCR) and seven transmembrane (7TM) receptor families were both enriched in the specific-divergent category; like NHRs, these gene families have rapidly expanded in *Caenorhabditis* (Thomas and Robertson 2008). Since neuropeptide, GPCR, and 7TM expression tends to begin fairly late in embryogenesis, it will be interesting to see if these divergently expressed genes become more similar at later developmental stages. Overall, we find that most genes show conservation in expression pattern and breadth, with the genes with changes in expression pattern largely limited to those expressed at low levels or in a small subset of cell types.

### Comparing expression in homologous cells identifies the complexity of transcriptome evolution

Single-cell sequencing of *Caenorhabditis* provides a powerful approach to ask not just how expression of individual genes has changed across evolution, but also which cell types are more or less constrained in expression profile. We found that cell types can differ in multiple ways, including in overall transcriptome, in the expression of cell-type specific “marker” genes, and in the level of expression of non-orthologous genes.

We first measured the overall difference in cellular gene expression by calculating the Jensen-Shannon Distance on the expression of 1:1 orthologous genes between the homologous cell type in the two species (JSD_cell_; terminal cell type statistics available in Supp. Table 7). JSD_cell_ calculated on the homologous cell types varied between 0.33 and 0.68. As an example, the body wall muscle from the middle of the animal (BWM middle), had relatively high similarity across species with a JSD_cell_ of 0.37 (Figure 3A). Most highly expressed genes were expressed at similar levels in both species including a large group of muscle specific genes, such as the myosin light chain gene, *mlc-3*. In contrast, the thermosensory AFD neuron had less similar expression between species with a JSD_cell_ of 0.50, with more of its highly expressed genes having greater expression differences than BWM middle. For example, the gene of unknown function *droe-4* was expressed >1000-fold higher in AFD from *C. briggsae* compared to that from *C. elegans*. This may reflect a temporal difference in expression, since *droe-4* is expressed in AFD in *C. elegans* larvae (Taylor *et al*. 2021; Harris *et al*. 2023).

**Figure 3.**
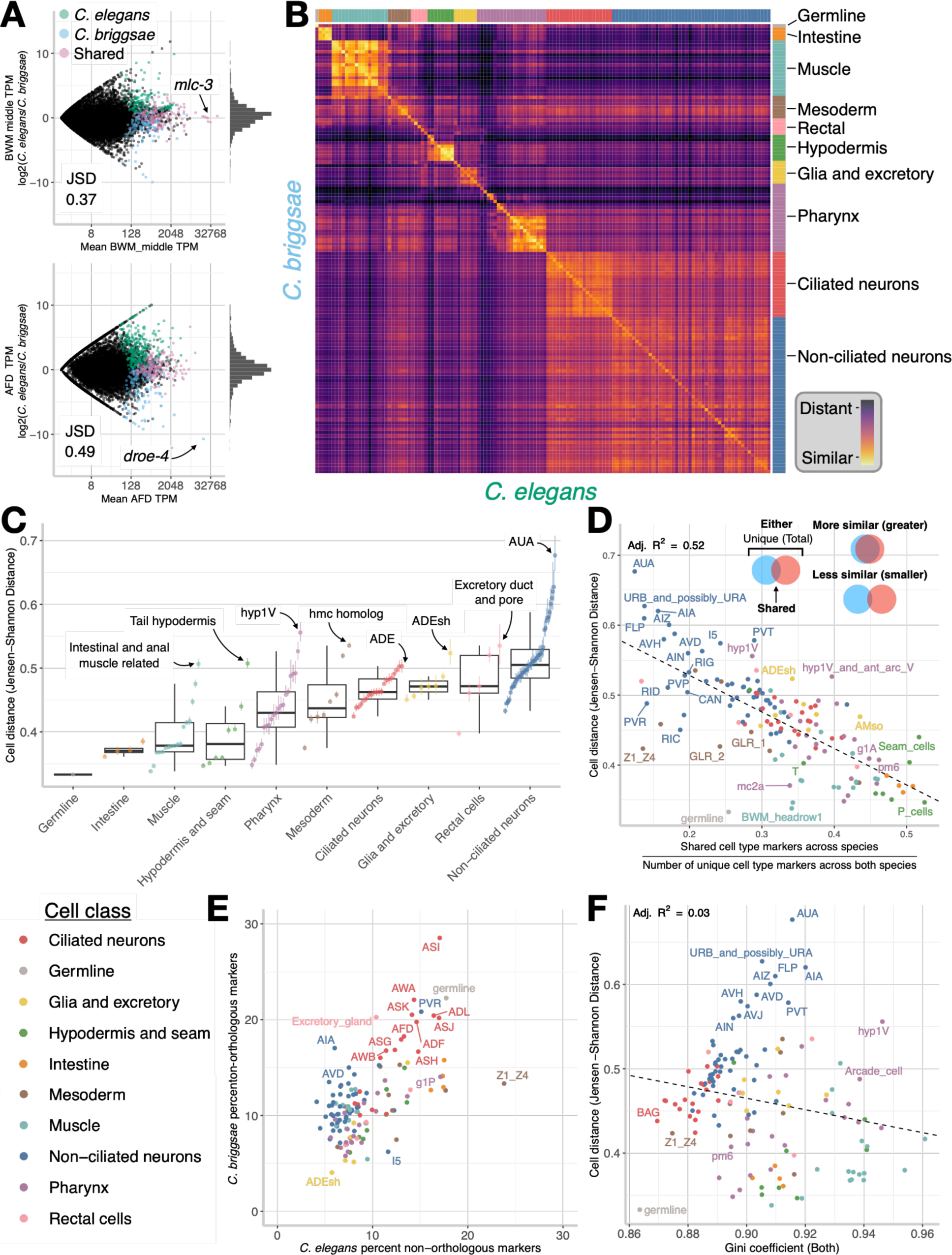
Cellular function defines divergence in transcriptomes between homologous cell types. (A) Pairwise comparison of the TPM values of all orthologous genes between *C. elegans* and *C. briggsae* for the AFD neuron and BWM middle shows overall similarity in transcriptome profile. Gene markers of the respective cell types are overlaid. (B) The cell distance, calculated as the Jensen-Shannon distance (JSD_cell_) between all terminal cell types. (C) The homologous cell type distances (median and 95% confidence interval of 1,000 bootstrapped values). (D) The quantity of cell type specific markers that are shared between *C. elegans* and *C. briggsae* divided by the total number of cell type markers in either species for the orthologous set of genes, compared with the cell distance. (E) The percent of the cell type markers that are not 1:1 orthologs between the species. (F) The mean Gini coefficient between both species of each terminal cell type compared to the cell distance.

Importantly, calculating JSD_cell_ between non-homologous cell types generally resulted in higher differences, for example 0.78 or 0.73 between AFD of one species and BWM middle of the other species. Comparing JSD_cell_ between all terminal cell types between the two species (Figure 3B) yielded several observations. First, most cell types are most similar by JSD_cell_ to their homolog in the other species. Homologous cells are the reciprocal best match 93.6% of the time, consistent with an accurate annotation of homologous cell types. In all cases where the most similar cell type is not the annotated homolog, the best match is a similar cell type of the same class (such as a muscle cell type matching a different, but closely related muscle cell type). Based on markers, these cases appear to be correctly annotated. Second, large blocks of cell types corresponding to cell classes (e.g. muscle, ciliated neuron, hypodermis) had higher similarity with other cell types of that class from the other species. This mirrors a similar structure seen for within-species comparisons (Supp. Figure 27-28) and likely results from conservation of broad features of the expression programs defining cell classes such as pan-neuronal, cilia or muscle genes. Some cell classes lacked these prominent similarity blocks, such as specialized mesodermal cell types and rectal cells, suggesting these classes may have more specialized expression profiles.

Grouping the JSD_cell_ by cell class revealed that there was variation both between and within major cell classes (Figure 3C). JSD_cell_ was lower for the germline, muscle, and intestinal and hypodermal cells, with pharyngeal and mesodermal cells having intermediate JSD_cell_ values, and neurons, rectal cells and glia having the most divergence (Figure 3C). However, JSD_cell_ varied substantially within cell classes, especially for the specialized mesodermal cell types, epidermis, pharynx and non-ciliated neurons, potentially reflecting the diversity of functions performed by cell types in these classes.

While JSD_cell_ measures overall transcriptome similarity, it does not account for the conservation of cell-type specificity of gene expression. To measure the level of conservation of genes that are cell-type specific, we first identified marker genes of every cell-type by testing whether genes had higher expression in that cell compared to all other cells from that species (Supp. Table 8-9). We then used the overall sharing of these cell type markers to measure the degree of conservation of cell type specific genes. For example, 40.6% of the 394 total BWM middle cell type markers were shared between the species versus 31.7% of the 586 total AFD markers, consistent with the higher similarity of BWM middle as measured by JSD_cell_. These shared markers were highly enriched for annotations expected for each cell type, such as Muscle Function, Cytoskeleton, Glycolysis for BWM middle, or Cilia, Neuronal Function, G-Protein for AFD (Supp. Figure 29). Cell-type markers that were not 1:1 orthologs between the species had less enrichment for these expected categories, suggesting conservation of marker status may be a useful tool to identify core cellular functions (Supp. Figure 30). Comparing the JSD_cell_ with the overlap of the orthologous markers between species revealed consistency in their rank order (Adj. R^2^ of 0.52; Figure 3D, Supp. Figure 31; See Supp. Figure 32 for a comparison of JSD_cell_ with Pearson correlation, Adj. R^2^ of 0.88). Some cell types including the germline and somatic germline precursor (Z1/Z4) showed lower fraction of overlap in their cell type markers than expected given their low JSD_cell_. Some of these non-shared markers show substantial differences in Z1/Z4 in *C. briggsae* compared to *C. elegans* while others were still robustly expressed in the other species, but below the fold-enrichment or significance threshold needed to be identified as a marker (Supp. Figure 33). Taken together, the degree of cellular gene expression conservation between species appears to be largely driven by a cell’s molecular function.

### Genes with complex orthology relationships show cell-type specific expression, but do not drive overall cell type conservation

While the analysis above focuses on the 1:1 orthologous gene set, cell types varied substantially in the fraction of their markers with complex orthology relationships in the two species (1:0, 1:many and many:many; Figure 3E). Certain cell types, including ciliated neurons and the somatic germline precursors, had a higher proportion than others of species-specific markers that had no identified ortholog in the other species (1:0 relationship). This is consistent with previous observations that certain families of genes involved in the sensory systems of *C. elegans* and *C. briggsae* are rapidly expanding and undergoing positive selection (Thomas and Robertson 2008). Other cell types, including muscle and epidermal cell types, were enriched for genes with 1:multiple or multiple:multiple orthology relationships, possibly due to an increased role of paralogous genes in these cell types, such as actin genes in muscle or collagen genes in the epidermis (Supp. Figure 34).

We tested whether cell types have similar divergence in their transcriptome when incorporating the 1:1 orthologous gene set alone vs if the non-1:1 orthologous genes are included as well. To do this, we organized the genes into orthogroups (sets of orthologous genes), then calculated orthogroup TPM values by summing the expression of all genes within that orthogroup (Emms and Kelly 2015, 2019). This allowed us to calculate the JSD_cell_ using these orthogroup expression values. This approach included an additional 4,290 *C. elegans* genes and 4,028 *C. briggsae* genes compared to the 1:1 ortholog set. The JSD_cell_ calculated on the orthologous genes or orthogroups led to a highly similar ranking of cell types (R^2^ = 0.96; Supp. Figure 35).

### What drives cellular transcriptome conservation?

To identify factors that could potentially explain why some cell classes have higher or lower expression similarity, we tested whether we could predict terminal cell type transcriptome conservation by using a multivariate linear model. We identified a model that incorporated 1) cell class, 2) Gini coefficient and 3) number of genes detected in that cell type as the simplest model that explained the most variance in the data (adjusted R^2^ ≈ 0.78, Supp. Table 10, model A). Gini is a metric of expression inequality and transcriptome complexity that can vary between zero and one, with low values indicating relatively equal expression of many genes and high values indicating a few highly expressed genes comprising much of the transcriptome. The large fraction of variance explained by our model suggests that cell type gene expression conservation, as measured by JSD_cell_, is positively related to cellular function and the number of genes expressed in a given cell type. We found that the number of cells recovered for that cell type had minimal additional explanatory power when added to the model (Supp. Table 10, model B). Consistent with this, downsampling the dataset to an equal number of cells per cell type leads to similar ordering of JSD_cell_ (Supp. Figure 36; R^2^ = 0.65). While the Gini coefficient had a significant effect on JSD_cell_ in the model, it had almost no measurable linear effect when tested for its relationship to JSD_cell_ in the absence of cell class, (Figure 3F; Adj. R^2^ = 0.03; P = 0.018). Together, this suggests that cell class is the dominant feature explaining the cellular expression divergence, with transcriptome complexity, as measured by Gini and number of genes expressed, explaining additional variation within each class.

### Progenitor cell transcriptome comparisons show diverse stage- and lineage-specific patterns of expression conservation

*C. elegans* and *C. briggsae* share nearly identical invariant lineages (Zhao *et al*. 2008; Memar *et al*. 2019), allowing a direct comparison of specific lineal progenitor cells between species. We used the same metrics as those described above to compare each progenitor cell state between species (Figure 4A; Supp. Table 11-12).

**Figure 4.**
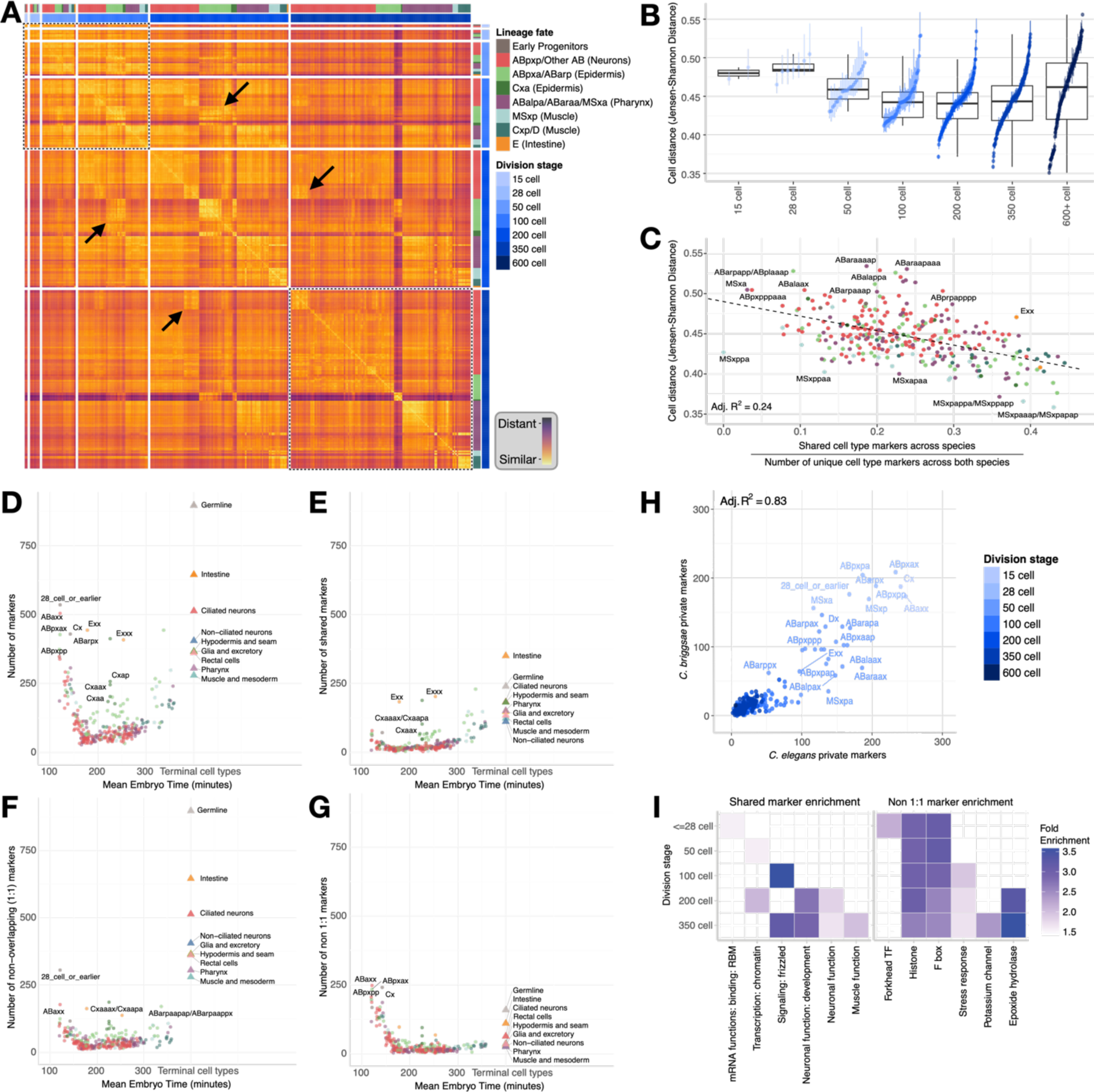
Progenitor transcriptome conservation and gene expression specificity depends on developmental stage. (A) Heatmap illustrating the transcriptome divergences between all progenitor cells across the two species (JSD_cell_). The dashed boxes highlight the early and late within-stage patterns. The arrows indicate off-diagonal similarities with cells from the same lineage. (B) JSD of homologous cell types, organized by their respective division stage. (C) The proportion of shared progenitor cell type-specific markers between *C. elegans* and *C. briggsae*, normalized by the total number of cell type markers in either species within the set of 1:1 orthologous genes, compared to the JSD_cell_. (D-G) Dynamics of cell type markers in *C. elegans* over time: the number of total cell type markers (D), markers overlapping between *C. elegans* and *C. briggsae* (E); non-overlapping *C. elegans* markers with a 1:1 ortholog in *C. briggsae* (F); and *C. elegans* markers with no 1:1 ortholog in *C. briggsae* (G). Circles represent progenitor cell types, while triangles indicate mean values for all terminal cells within each terminal cell class. (H) Comparison of the number of non 1:1 markers in *C. elegans* progenitor cells with that in homologous progenitor cell type in *C. briggsae*. (I) Fold enrichment of WormCat categories for genes that serve as shared markers across homologous progenitor cells (left) and those that are non 1:1 in *C. elegans* (right) - all terms shown are significantly enriched (P < 0.05) in at least one progenitor cell type.

Comparing the transcriptomes of each *C. elegans* progenitor with each *C. briggsae* progenitor state using JSD_cell_ identified several patterns (Figure 4A). First, most cells are more similar to their homologous cell state in the other species than to other progenitor states (apparent as a strong diagonal in Figure 4A). This provides transcriptome-wide support for the accuracy of our marker-based annotations. Second, comparisons between non-homologous cells show that early cells in one species are more similar to other differently-annotated early cells than to later cells in the other species (upper-left region of Figure 4A; Supp. Figure 37). This indicates the presence of a conserved expression signature of early-embryonic states. Third, progenitors from different lineage origins that will produce certain cell types, including muscle, epidermis and intestine, tend to be more similar to other similarly fated progenitors than to progenitors of other cell types (as indicated by the arrows pointing to the off-diagonal patterns in Figure 4A). This indicates that these progenitors have conserved cell class-specific expression profiles that arise prior to their exiting the cell cycle. Similar patterns are seen for within-species and between-species comparisons, emphasizing that relationships between progenitor transcriptomes are largely conserved (Supp. Figure 38-39).

We next asked whether the degree of similarity between homologous progenitor cells across species varied as a function of embryonic development time (Figure 4B). The overall distribution of JSD_cell_ in progenitors (∼0.35 - 0.59) was similar to that in terminal cells (∼0.35-0.55). Median JSD_cell_ values declined across stages from the 28-cell to a minimum distance at the 200-cell stage before rising slightly in later progenitors and terminal cells. The higher expression similarity between species at an intermediate time point is reminiscent of the “developmental hourglass” model (Kalinka *et al*. 2010; Levin *et al*. 2016). However, the temporal patterns of JSD_cell_ varied between individual lineage fates, suggesting that progenitor expression conservation across time is complex.

Similar to our analysis of terminal cells, we used cell type markers to assess the conservation of progenitor cell type-specific expression programs. We identified markers of each progenitor state in each species by testing for differential expression in that state compared to other progenitors. The fraction of progenitor cell markers shared between species was modestly correlated with JSD_cell_, but many cells had higher or lower marker overlap than predicted from their JSD_cell_ value (Figure 4C). Intriguingly, these included MSxppa and MSxppaa, the parent and grandparent of the terminal cell type Z1/Z4 (the somatic gonad precursors), which also had low marker overlap relative to JSD_cell_ (Figure 3D). Others, including intestinal progenitors and certain cells from the AB lineage, had relatively high marker overlap relative to their JSD_cell_. This could reflect certain lineages expressing genes associated with their terminal states earlier in development than others, as the intestine program in the E-lineage is known to be expressed especially early (Hashimshony *et al*. 2015).

We observed striking temporal patterns in the number and conservation of progenitor markers (Figure 4D-G). First, both the total number of markers (an estimate of that cell’s transcriptome distinctiveness) and proportion of shared markers (reflecting conservation of cell type specific expression) was generally lower in progenitors than in terminal cells, especially prior to 150’ (∼200-cell stage; median of 17.5 shared markers, 14% versus a median of 143 shared markers, 29% in terminal cells). A few progenitor states, including the E-derived intestinal progenitors and C- and AB-derived epidermal progenitors were exceptions to this and had high numbers of shared markers even at earlier stages, consistent with a conserved early onset of differentiation in these lineages (Figure 4E). Second, markers of early (<150’) progenitors are heavily enriched for markers that do not have 1:1 orthologs (Figure 4G); a pattern that is consistent across the two species (Figure 4H). Third, during the early stages, most progenitors had more markers with 1:1 orthologs that were not identified as markers in the other species (Figure 4F). Finally, at later stages, both the total number of markers and number of shared markers increased (Figure 4D, 4E). Overall this suggests broad stage-specific differences in constraints on gene expression and gene family evolution.

Given these patterns, we asked whether 1:1-orthologous and non-orthologous markers of progenitor states were associated with different functional annotations in *C. elegans* (Figure 4I). In general, shared markers were associated with fundamental developmental processes like mRNA-binding, signaling and chromatin regulation at earlier stages, and at later stages with appropriate cell type-specific annotations such as “muscle function” (Supp. Figure 40). In contrast, non-1:1 orthologous markers in early progenitors were associated with different annotations, such as ubiquitin-mediated proteolysis (F-box genes) and chromatin structure, which reflects the duplicated clusters of histone genes (Supp. Figure 41). These patterns suggest that different evolutionary constraints act on early progenitors versus late progenitors or terminal cells.

## Discussion

Here, we have compared the conservation of cell type transcriptome program and pattern of gene expression across development for two related species. Despite an estimated 1.78 substitutions per neutral site between the species (Stein *et al*. 2003), the developmental gene regulation of *C. elegans* and *C. briggsae* remain highly conserved. Nonetheless, thousands of individual genes have shifted in expression patterns in both terminal and progenitor cell types with the most notable changes being genes that are either lowly expressed or are highly specific in their expression pattern. We found specific gene families enriched for their divergence in gene expression patterns (neuropeptides, NHRs, and 7TM/GPCRs). These gene families showing higher divergence largely agrees with previous work that has found cell type specific variation in the expression of the *nlp-21* neuropeptide between strains of *C. elegans (Ben-David et al. 2021)* and the *frpr-14* neuropeptide between *C. elegans* and *C. briggsae (Chai et al. 2022)*. As these gene families, along with the F-boxes, constitute young and expanding gene families (Ma and Zheng 2023), it supports the growing body of evidence that gene duplication drives changes in the gene regulatory program (Ma *et al*. 2022).

A major goal of this work is to understand the evolution of gene expression, its constraints, and whether gene regulatory changes drive phenotypic innovation. Previous analyses have used RNA-seq on whole tissue or whole organs to compare the evolution of gene expression between species (Khaitovich *et al*. 2006; Brawand *et al*. 2011; Necsulea and Kaessmann 2014; Chen *et al*. 2019; Cardoso-Moreira *et al*. 2019; Fukushima and Pollock 2020; El Taher *et al*. 2021). These studies have found variation in the rate of transcriptome evolution between tissue types, with neuronal tissue being the most conserved and reproductive tissue being the most divergent. Yet, these analyses have the caveat that they are based on the average expression across the individual cells within each tissue and that the proportion of cell types inside of a tissue can differ dramatically between species, and can lead to misestimation of gene expression evolution (Price *et al*. 2022). For example, (Francis *et al*. 2014) suggests rapid evolution of neurite localized genes between mice and rats. Single-cell sequencing has the promise to resolve these issues by capturing transcriptomes from individual cells, allowing for direct comparisons of gene expression between homologous cells. However, issues associated with batch effects, capturing a large quantity of cells at scale across an animal, and estimating the homology between cell types can complicate proper estimation of gene expression evolution at the cell type level. Instead, most analyses comparing species at a single-cell scale have focused on resolving the evolutionary origins of extant cell types, determining changes in cell type proportions between species, and/or the evolution of gene expression across a single tissue (La Manno *et al*. 2016; Tosches *et al*. 2018; Sebé-Pedrós *et al*. 2018; Lau *et al*. 2020; Shami *et al*. 2020; Tarashansky *et al*. 2021; Shafer *et al*. 2022; Murat *et al*. 2023; Lamanna *et al*. 2023; Zemke *et al*. 2023; Sepp *et al*. 2024).

Here, due to the unambiguous homology of the cell types (Zhao *et al*. 2008; Memar *et al*. 2019), the conservation of cell fates between *C. elegans* and *C. briggsae*, and our ability to capture cells across an animal’s entire embryonic development we were able to measure conservation of gene expression between cells of different classes over developmental time. We found that the transcriptome of the germline precursors are highly conserved relative to other cell types such as the neurons and glia. These findings are in contrast to the observations within amniotes using bulk RNA-seq, where the testes and ovaries show the highest degree of divergence and the brain shows the most similarity (Khaitovich *et al*. 2006; Brawand *et al*. 2011; Necsulea and Kaessmann 2014; Cardoso-Moreira *et al*. 2019; Fukushima and Pollock 2020). More recent investigations using single-cell RNA-seq on whole testes from diverse mammals have found that spermatogonia and somatic support cells are more conserved than later differentiated cell types (Murat *et al*. 2023). A further comparative analysis of rabbit and mouse development additionally found higher similarities of neuronal tissue versus gut (Ton *et al*. 2023). Whether differences in transcriptome conservation of broad tissue and cell class categories between mammalian and *Caenorhabditis* systems reflect biological, technical, or embryo staging differences remains unclear. Further work applying single-cell RNA-seq to a broad set of tissues from a diverse set of animals to measure gene expression pattern conservation will help resolve these discrepancies.

Overall, these datasets allow for the investigation of specific differences in cell-type specific gene expression across embryonic development in *C. elegans* and *C. briggsae*. An explorable version of the datasets is available online (https://github.com/livinlrg/C.elegans_C.briggsae_Embryo_Single_Cell).

## Methods

### Strains, single-cell collection, and nematode culturing

Within this study, new single-cell RNA-seq data was collected for both *C. elegans* and *C. briggsae* embryos. The additional *C. elegans* (both wild-type and mutant) and the *C. briggsae* cells were reared, synchronized and collected as described previously in (Packer *et al*. 2019). The *C. briggsae* strain was an isolate of AF16. The *C. elegans* collections were of one wild-type N2 (VC2010) strain, and from several strains that were deletion mutants for a transcription factor: *ceh-9*(*tm2747*), *M03D4.4*(*gk5269*), and *mec-3*(*gk1126*) and presumptive null alleles. Several lines of evidence suggest that the mutants have wild type gene expression patterns in most of their cell types. The homozygous mutant lines were backcrossed with VC2010 three times. Both *ceh-9* and *M03D4.4* mutant homozygotes have no readily perceptible phenotype and the *mec-3* phenotype is limited to failure to respond to light touch. *ceh-9* is primarily expressed in the SMB neurons with much weaker expression in SMD, RME and RIA neurons (at least 10-fold lower expression). *mec-3* is expressed primarily in the touch neurons (ALM/PLM) and at lower levels in the FLP1 neurons. *M03D4.4* is more broadly, but transiently, expressed, including in body wall muscle, pharyngeal cells and mesodermal cells, but is not expressed in neurons, hypodermis/seam, germline or intestine. Co-embedding of the mutant cells with wild type cells shows the cell types from the different sources generally cluster together. Expression analysis shows they share the same markers and overall expression patterns. However, there were exceptions. Notably, in the *mec-3* mutants the touch neurons (ALM/PLM) were absent, in agreement with previous findings (Way and Chalfie 1988), with the cells expressing the *mec-3* deletion allele limited to ALM/PLM precursors (the deletion is internal and preserves the 3’ UTR and one of the transcription start sites). In the *M03D4.4* mutants, while the mutant cells generally co-clustered with wild type cells, some expression timing differences were observed in the body muscle cells, with some genes being expressed earlier in the *M03D4.4* mutant cells than in wild type. In the *ceh-9* mutants, cells co-clustered with SMB, SMD, RME and RIA neurons from wild type and the other mutants, and showed only minimal differences in expression patterns with wild type. In particular, inclusion of the *M03D4.4* cells for body wall muscle, pharynx and mesodermal cell type measurements may generate higher variability for some genes. But the M03D4.4 cells comprise less than 15% of the total *C. elegans* cells, meaning the impact on the analysis is unlikely to be large.

### Analysis of single-cell data

The single-cell RNA-seq reads were mapped using the 10X genomics CellRanger pipeline to a modified version of the WormBase WS260 *C. elegans* and *C. briggsae* reference genomes (Davis *et al*. 2022). As was previously done for the *C. elegans* reference genome in (Packer *et al*. 2019), the *C. briggsae* 3’ UTR annotations were similarly extended to improve gene expression quantification. After alignment, the *C. briggsae* and new/mutant *C. elegans* datasets were filtered for cells that were likely doublets between two different cell types. Within all cells, the percentages of reads coming from the background ambient RNA profile was determined similar to (Packer *et al*. 2019). Newly collected cells that were estimated to have more than 20% of reads coming from the ambient background were removed.

### Defining the list of C. elegans and C. briggsae 1:1 genes

First, all *C. briggsae* names assigned by WormBase to a *C. elegans* gene name were defined as common genes (e.g. the *C. briggsae* gene given the name by WormBase of *Cbr-aat-2* was defined as a 1:1 ortholog with *C. elegans aat-2*). Similarly, if there were other *C. briggsae* genes listed as orthologs to a *C. elegans* gene that already had a named *C. briggsae* ortholog, the other *C. briggsae* genes were not considered common genes.

Second, the *C. elegans/C. briggsae* WS260 orthologs were obtained from WormBase. For any 1:1 ortholog pair, regardless of synteny, the *C. elegans/C. briggsae* pair was considered to be “common.” For all other pairs, synteny was used to define other genes in common as follows: For each *C. elegans* chromosome, the *C. elegans/C. briggsae* ortholog pairs (1:1, 1:many, many:1, and many:many) were ordered along the chromosome by their position in *C. elegans*. As noted above, for those genes that WormBase had already assigned a *C. elegans* name in *C. briggsae*, all other *C. briggsae* genes that were not defined as being in common with *C. elegans*. If the *C. briggsae/C. elegans* ortholog pair were on the same chromosome in both species, and the order of the genes was retained with respect to any of the five neighboring genes on either side of the gene of interest, as long as there were not multiple copies of either the *C. elegans* or the *C. briggsae* gene within that region, by synteny, the pair was kept as “common” and any other *C. elegans* or *C. briggsae* gene in the orthologous group was not considered for the common list. If the *C. briggsae* gene was not on the same chromosome as the *C. elegans* ortholog pair, but there was a block of *C. briggsae* genes in order along the same *C. briggsae* chromosome without multiple copies of that gene in the same block of genes, again by synteny, the genes in the block were considered to be “common” genes and other pairs from the orthologous groups were discarded. If a single *C. briggsae* gene was a many:1 with respect to *C. elegans*, and one of the *C. briggsae* ortholog pairs was on an unlocalized *C. briggsae* contig, but all other *C. briggsae* genes were on a chromosome that did not agree with the placement in *C. elegans*, then the copy on the unlocalized *C. briggsae* contig was retained as a gene in common based on the assumption that unlocalized contigs could potentially be on the same chromosome as the *C. elegans* chromosome. Similarly, all 1:many, many:1 and many:many genes that could not be defined as common genes by synteny were removed from consideration. In this way, genes that were not 1:1 *C. elegans/C. briggsae* orthologs were included in the list of “common” genes. In total, there are 12,803 common genes of which 11,120 are 1:1 orthologs.

### Embryo time estimation

As computed previously in (Packer *et al*. 2019), the developmental time of the embryo from which each cell was collected was estimated. Briefly, the log normalized expression data of individual cells were correlated with log normalized bulk-RNA-seq data from (Hashimshony *et al*. 2015). For the *C. briggsae* cells, the estimated embryo time was calculated using the 1:1 orthologous genes between *C. elegans* and *C. briggsae*. For both species, the estimated embryo time matched with expected age distributions given the collection scheme (Supp. Figure 2).

### Single-cell integration and UMAP generation

To jointly project the two species data into the same reduced dimensional space, we used canonical correlation analysis (CCA), implemented in Seurat V4. Both species gene by cell matrix were first subsetted to the 1:1 orthologs. The cells of both species were then jointly split by collection batch and each batch was run through a standard pipeline of normalization and finding the 2,000 most variable features. Subsequently, the datasets were integrated by batch. We found that integrating by batch instead of by species resulted in both a faster run-time and better integration.

After integration, the data was scaled and projected into 300 principal components, and further reduced to 2 dimensions using UMAP implemented in SeuratV4 (4.3.0). Clusters were identified using louvain clustering and matched to tissue classes using known marker genes. Cells belonging to these tissue classes were further subsetted, reintegrated using RPCA, and re-projected for subsequent cell type annotation (Supp. Figures 6-12). Additional cell subsets based on the inferred embryo time were similarly reintegrated using CCA and re-projected for subsequent progenitor annotation (Supp. Figure 14-17).

### Cell-type annotation

*C. elegans* and *C. briggsae* terminal cell types were annotated jointly in an integrated space to have direct homology between like-cells. The terminal cell types were annotated using markers, trajectory relationships, and embryo time as described previously (Packer *et al*. 2019). *C. briggsae* progenitor cell types were annotated similarly to as described previously for *C. elegans* (Packer *et al*. 2019). Cells were labeled within both the tissue and embryo time based sub-UMAPs using known markers that were homologous between *C. elegans* and *C. briggsae* (Supp. Table 3).

### TPM calculation, bootstrapping, and number of genes detected

To generate an aggregated quantification of each cell type’s gene expression, we calculated a relative expression metric, the Transcripts Per Million (TPM) for each cell type. As 10x single-cell RNA-seq data is 3’ biased, there is no requirement for gene size correction of the TPM values. First, the background corrected gene by cell matrix was normalized for library sequencing depth and complexity by dividing by a per-cell size factor (calculated in Monocle3). Second, the matrix was subsetted to the cells corresponding to a single cell type in one species. Next, the mean expression of every gene was calculated. Finally, each cell type’s expression vector was scaled to 1,000,000 using each species’ full gene set. The process was repeated for every cell type from either species. The proportion of relative expression coming from each species non-1:1 orthologous set differed between *C. elegans* and *C. briggsae* by ∼8%, with *C. briggsae* having more relative expression originating from non-1:1 genes.

As the quantification of an aggregate gene expression value can be skewed by few aberrant cells, we determined the variance in a gene’s quantification through a confidence interval using bootstrapping. Using the same TPM calculation procedure as described above but before calculating the mean expression, the cells of a given cell type were randomly resampled across 1,000 iterations. Then, a median value and 95% confidence interval was calculated on the iterated values. We then determined whether a gene was confidently detected in a cell type based on whether its 95% confidence interval was greater than zero.

### Jensen-Shannon distance, Spearman/Pearson correlation, and Gini coefficient calculation

To determine the relative conservation of cell type transcriptome and gene expression patterns between species, we calculated a variety of metrics including the Pearson correlation, Spearman correlation, Jaccard distance, Gini coefficient and Jensen-Shannon distance (JSD). The Pearson and Spearman correlation were calculated on log transformed TPM values with a small pseudocount added. The Jaccard distance was calculated on binarized expression data (whether a gene was confidently detected in that cell type). The Gini coefficient was calculated on all genes in each species separately using the reldist R package (1.7-1) on the TPM normalized data. The calculation of the JSD_gene_ and JSD_cell_ started with the addition of a small pseudocount equivalent to the inverse of the number of categories being calculated (number of cell types or number of genes respectively). Next, the expression in TPM across either all of the cell types (JSD_gene_) or across every gene (JSD_cell_) was converted into a probability distribution. Then the following calculation for the JSD was applied:

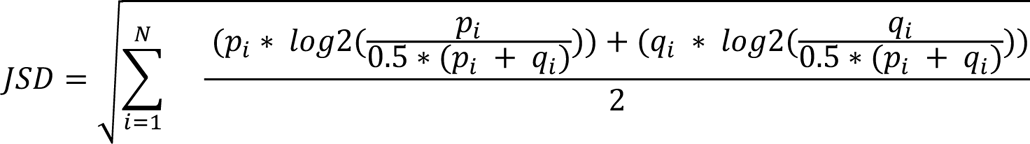

where p and q are the probability distributions for either species. To incorporate measurement noise into our calculation of cell distance, 95% confidence intervals surrounding the JSC_cell_ and the JSD_gene_ values were also estimated using the bootstrapped TPM estimates described above.

In comparison of these metrics, we found that the JSD_gene_ and JSD_cell_ best represented the conservation of expression profile and were least vulnerable to data quality. These observations came through both inspection of hundreds of individual cases and determining sensitivity of the different metrics to downsampling the number of cells and UMI’s (full set of gene and cell examples available at http://github.com/livinlrg/C.elegans_C.briggsae_Embryo_Single_Cell) (Supp. Figure 18, 36, and 42).

### Identification of cell-type specific markers

Using SeuratV4 (4.3.0), cell-type specific markers were identified using default settings of the FindAllMarkers function (Wilcoxon Rank-Sum test). For the terminal cell types, a given cell type’s gene expression specificity was compared against the rest of the dataset from that species. For the progenitors, the gene expression specificity was instead compared within just the progenitors of that species. The terminal and progenitor cell type markers were then filtered using a log2 fold-change cutoff of 1, and a Bonferroni adjusted p-val value cutoff of 0.05.

### WormCat enrichment and fold-change

The WormCat genes annotations (Holdorf *et al*. 2020) were downloaded from http://www.wormcat.com/ (version from November 11, 2021). The *C. briggsae* genes were annotated here by transferring the WormCat labels for the genes where a 1:1 orthologous gene had been identified. Using a background gene set consisting of either the 1:1 genes (Figure 2J), or the *C. elegans* genes (Figure 4H,I), a one-sided Fisher’s Exact test and Bonferonni correction was used to find category enrichments for a query gene set using methods and code derived from (Holdorf *et al*. 2020). A fold-enrichment versus the background gene set was then calculated on the query gene set using the following:

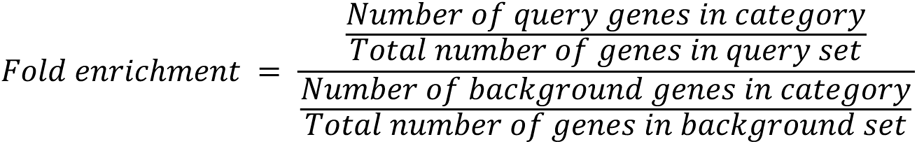

### JSD_Cell_ downsampling calculation

To ensure that the JSD_Cell_ calculations were not unduly influenced by biases in the cell collection rate of terminal cell types between *C. elegans* and *C. briggsae*, the JSD_Cell_ value was recalculated 10,000 times on randomly sampled ten cell subsets. First, each species cell by gene matrix was randomly downsampled to ten cells of each cell type. Second, the TPM was recalculated for each cell type on this downsampled set of cells. Finally, the JSD_Cell_ and Pearson correlation was calculated on the TPM matrix as described above.

### Cell-type specificity (Tau) calculation

The cell-type specificity index, *Tau*, was calculated as previously described in (Yanai *et al*. 2005). Briefly, the TPM normalized expression data was used to calculate the *Tau* metric for the progenitor and terminal-cell types independently using the following formula:

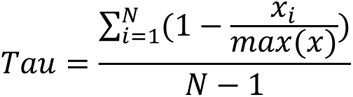

where N is the total number of cell types in either the progenitors or the terminal cell types, x_i_ is the TPM value of a gene in a cell type divided by that gene’s maximum expression across all progenitor or terminal cell types.

## Supplemental Figures

**Supp. Figure 1.**
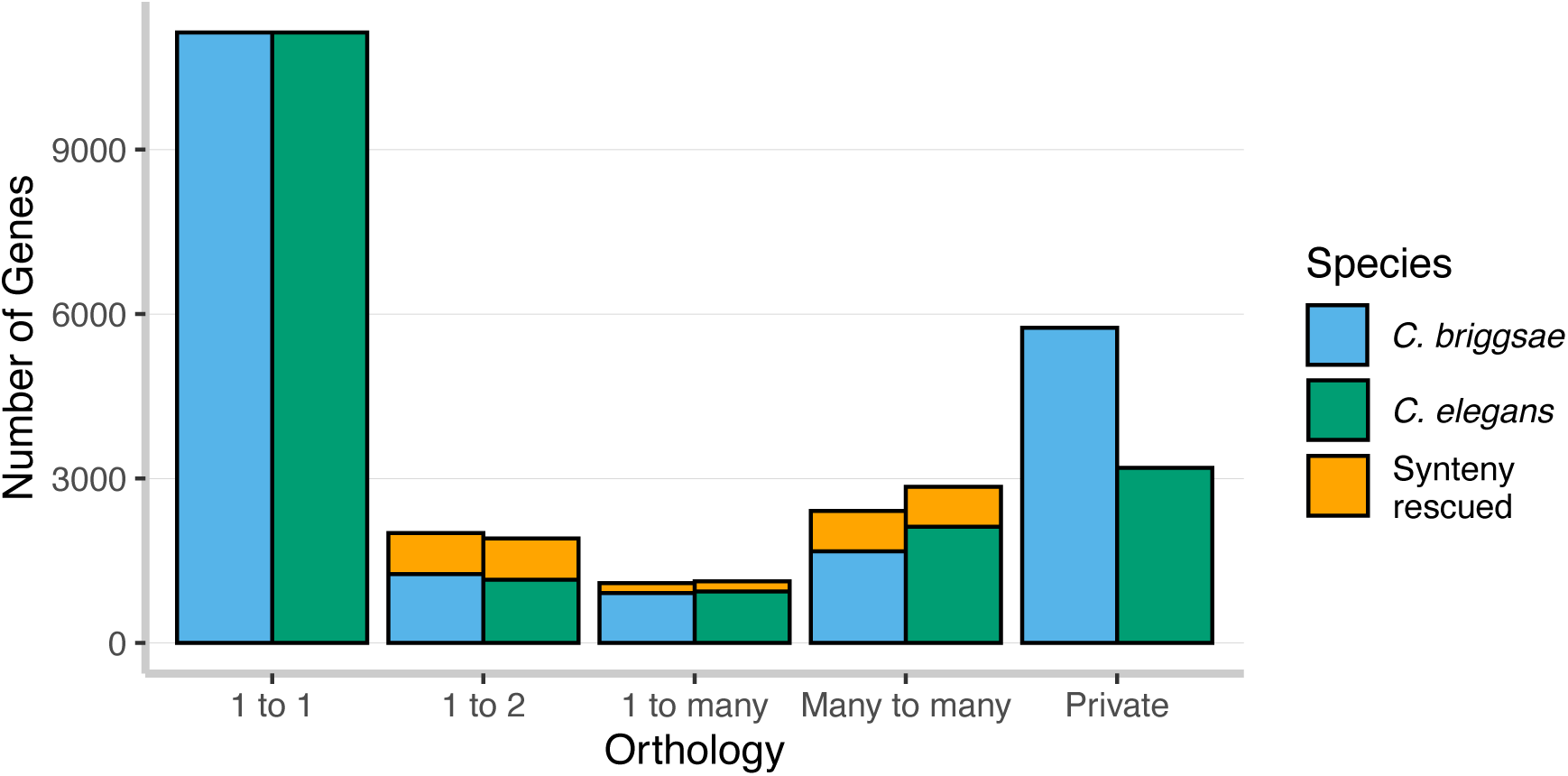
The orthology relationship of the genes in *C. elegans* and *C. briggsae* were downloaded from WormBase (WS260) and used to define the initial 11,120 1:1 orthology gene set. Then, an additional 1,683 genes were identified as being syntenic between the species and added to the 1:1 orthology gene set, totaling 12,803 total 1:1 orthologs. Genes where no ortholog was identified in the other species were considered private.

**Supp. Figure 2.**
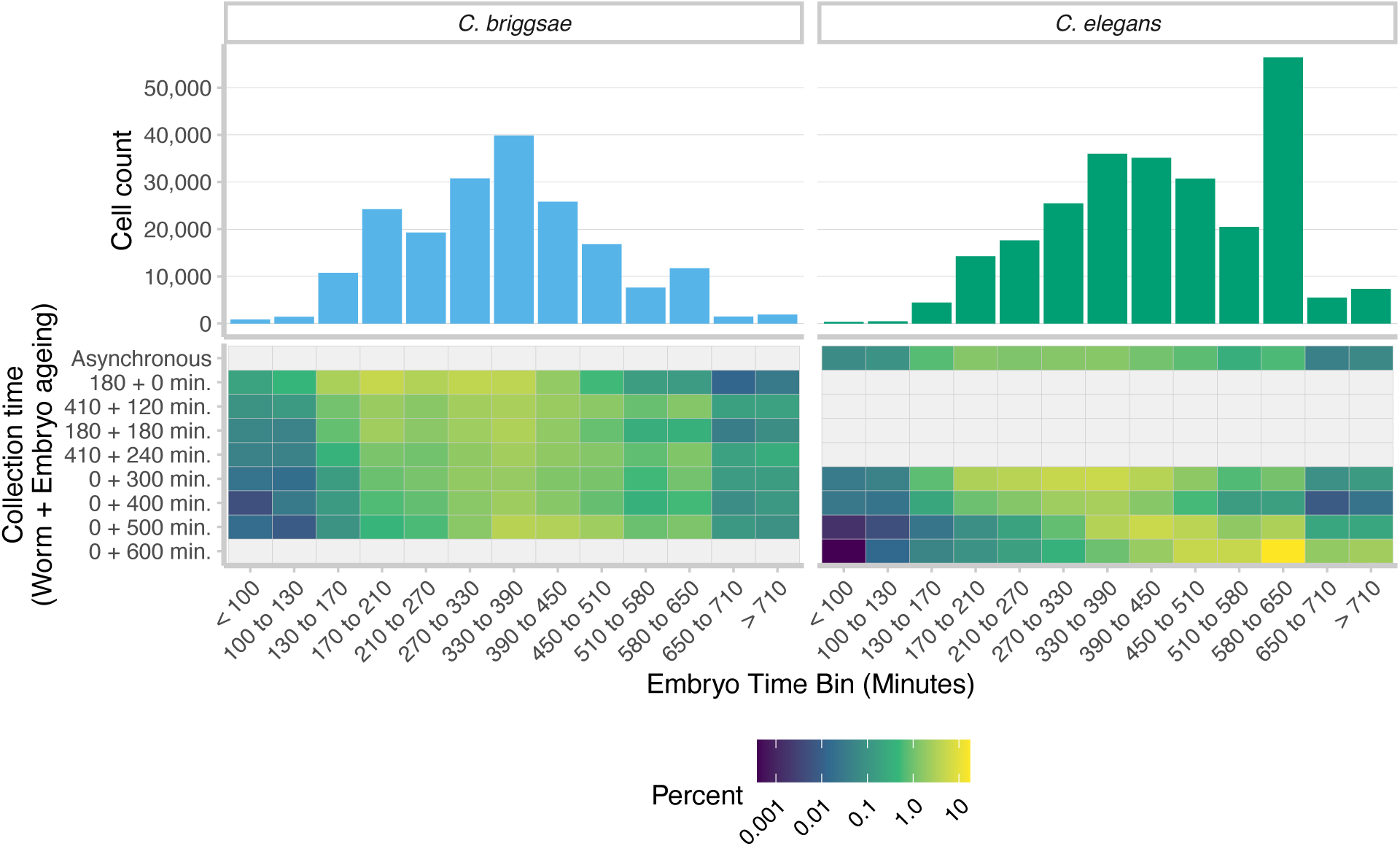
The distribution of cells within the estimated embryo time bins (top) for *C. elegans* and *C. briggsae* show a slight early bias in embryo collection for *C. briggsae* and slight late bias for *C. elegans*. The distribution of estimated embryo times matched with expectations given the embryo collection strategy (bottom). Embryos were aged both inside of the hermaphrodite (worm ageing) or after the embryos were released from the adult hermaphrodite with hypochlorite treatment (embryo ageing).

**Supp. Figure 3.**
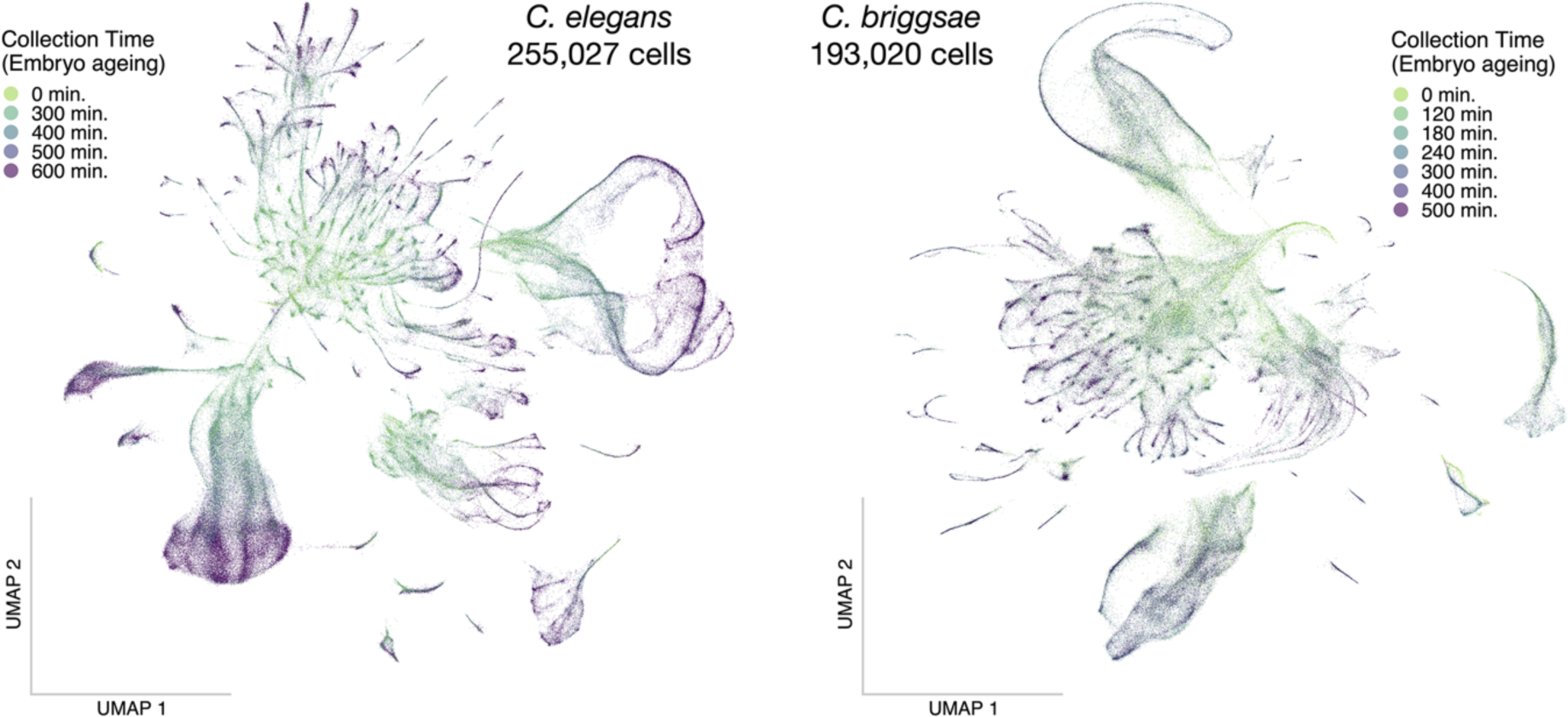
Batch integrated UMAPs of either the *C. elegans* or *C. briggsae* cells with the collection time of the embryos labeled for each cell. The cells from the youngest embryos are generally within the center of the projection, with later differentiated cell types spanning out.

**Supp. Figure 4.**
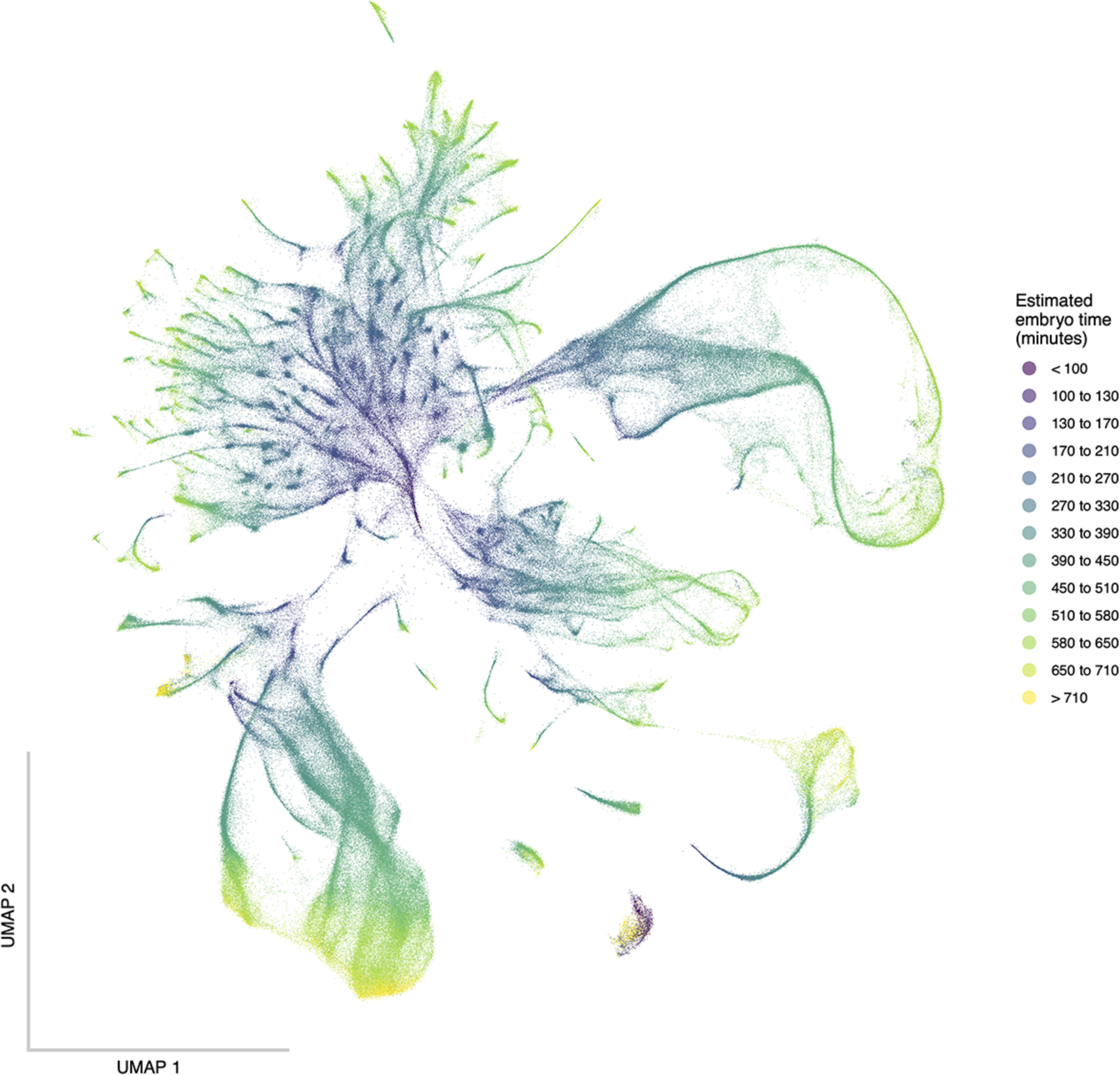
An integrated UMAP of *C. elegans* and *C. briggsae* cells together showing that the cells from the youngest embryos are near the center of the projection, with cells from older embryos projecting outwards.

**Supp. Figure 5.**
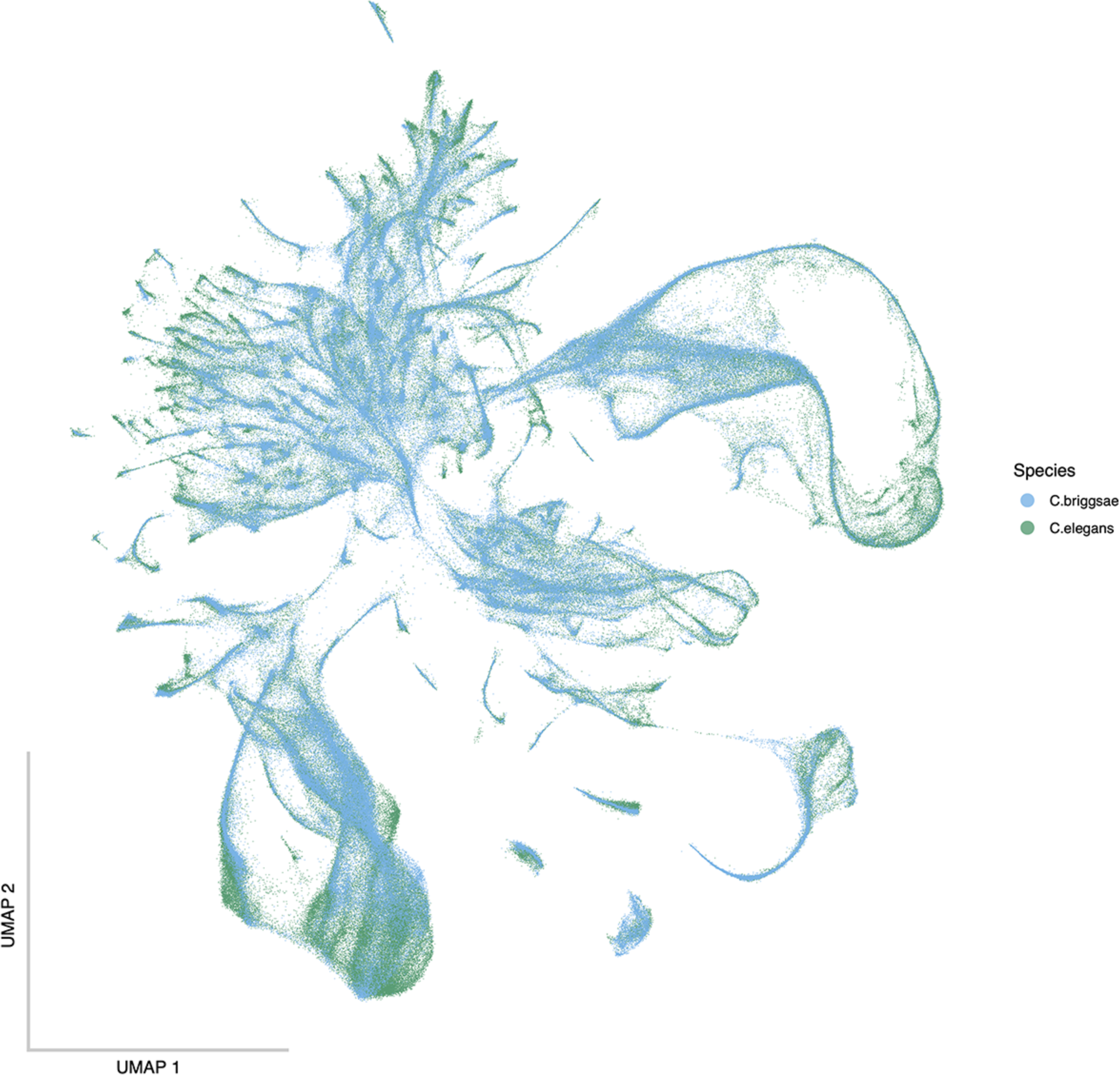
The same integrated UMAP as Figure 1A,B showing the species composition of the single-cells. The late bias of the *C. elegans* dataset is visible at the tips of the neuronal projections. Reciprocally, the early bias of the *C. briggsae* dataset is evident at the center of the projection. Overall, the datasets are well-integrated.

**Supp. Figure 6.**
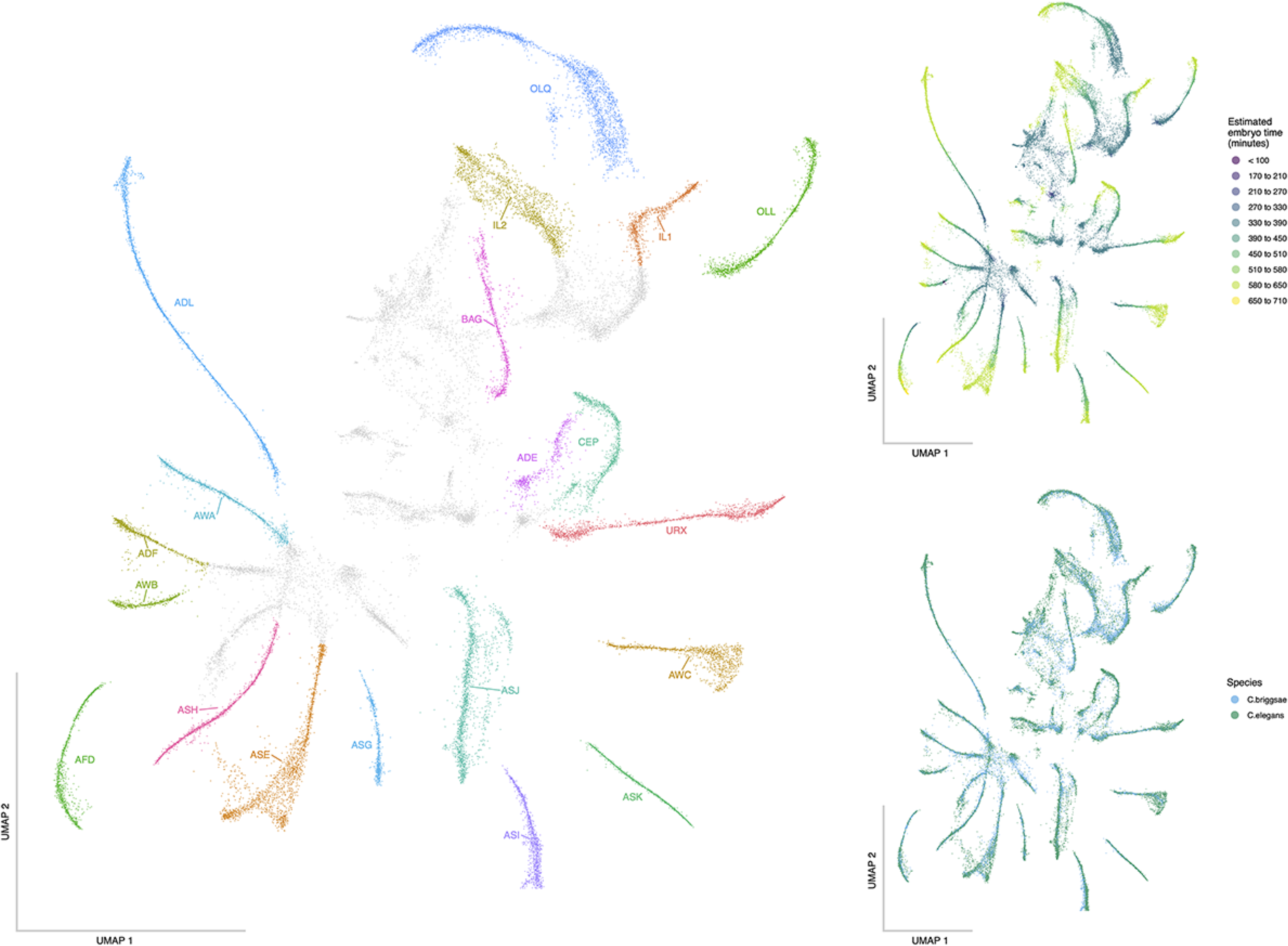
These plots show the ciliated neurons cell class subset integrated UMAPs. The cell labels (left), estimated embryo time (top-right), and the species composition (bottom-right) are shown.

**Supp. Figure 7.**
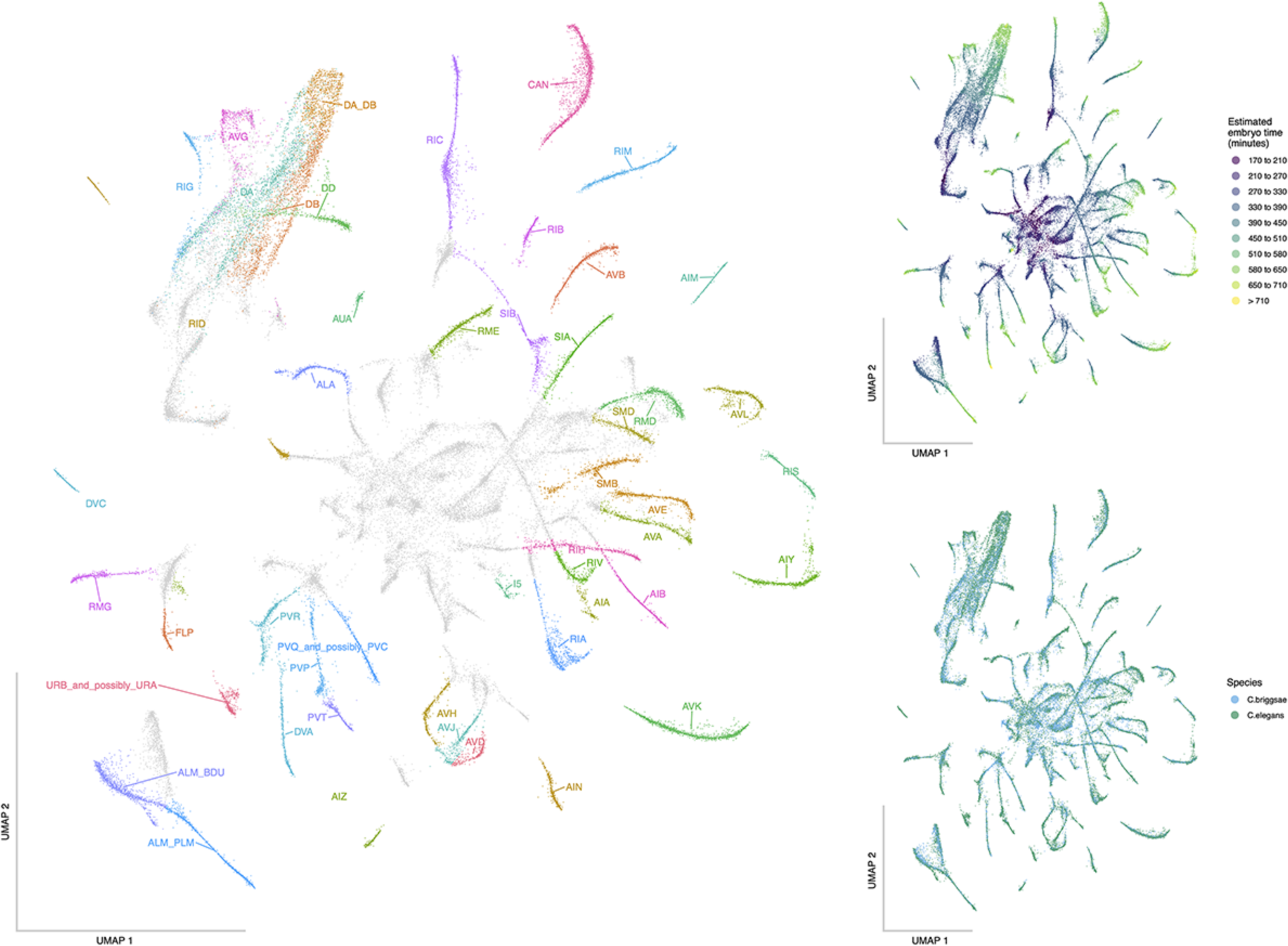
These plots show the non-ciliated neurons cell class subset integrated UMAPs. The cell labels (left), estimated embryo time (top-right), and the species composition (bottom-right) are shown.

**Supp. Figure 8.**
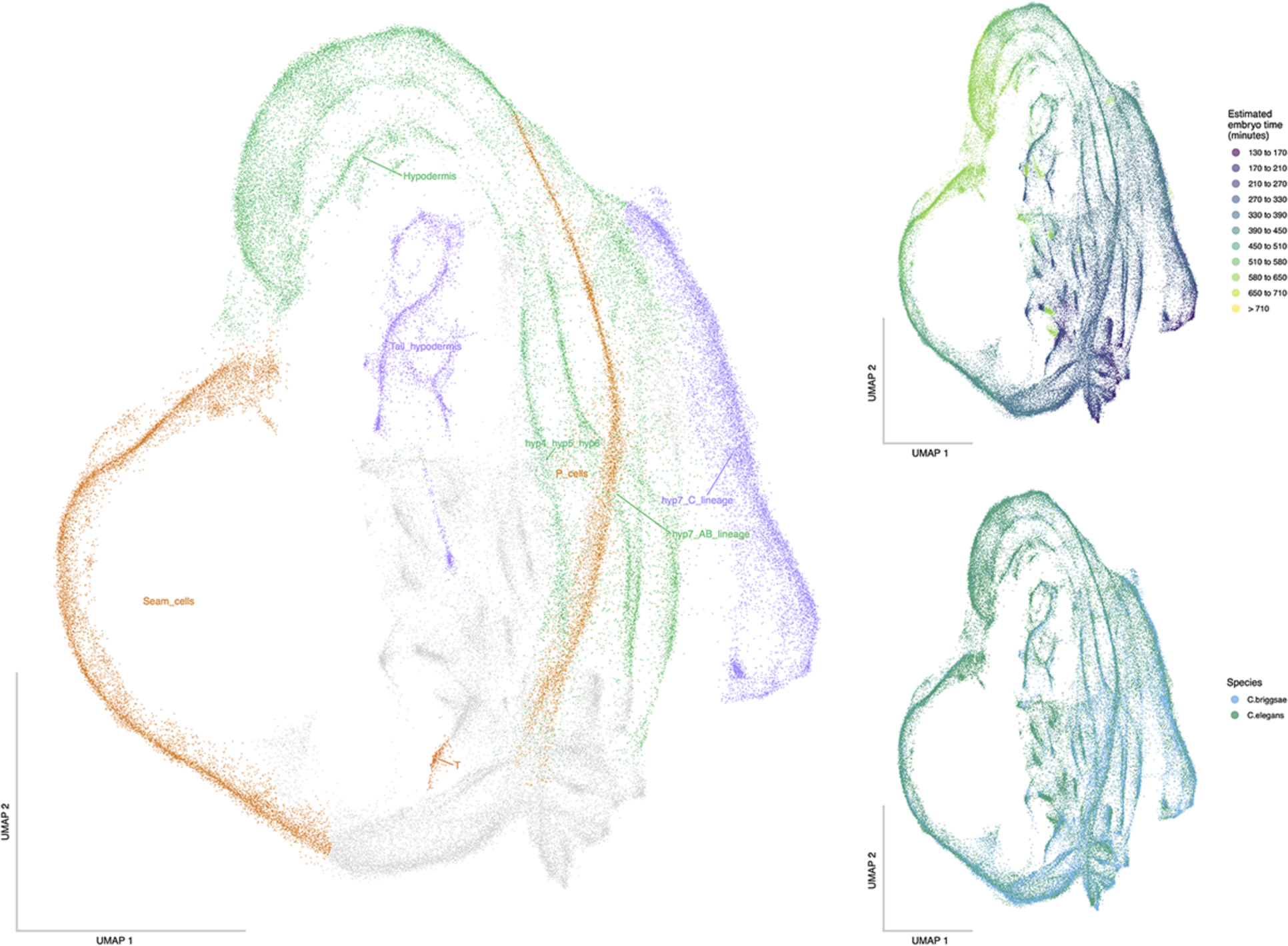
These plots show the hypodermis and seam cell class subset integrated UMAPs. The cell labels (left), estimated embryo time (top-right), and the species composition (bottom-right) are shown.

**Supp. Figure 9.**
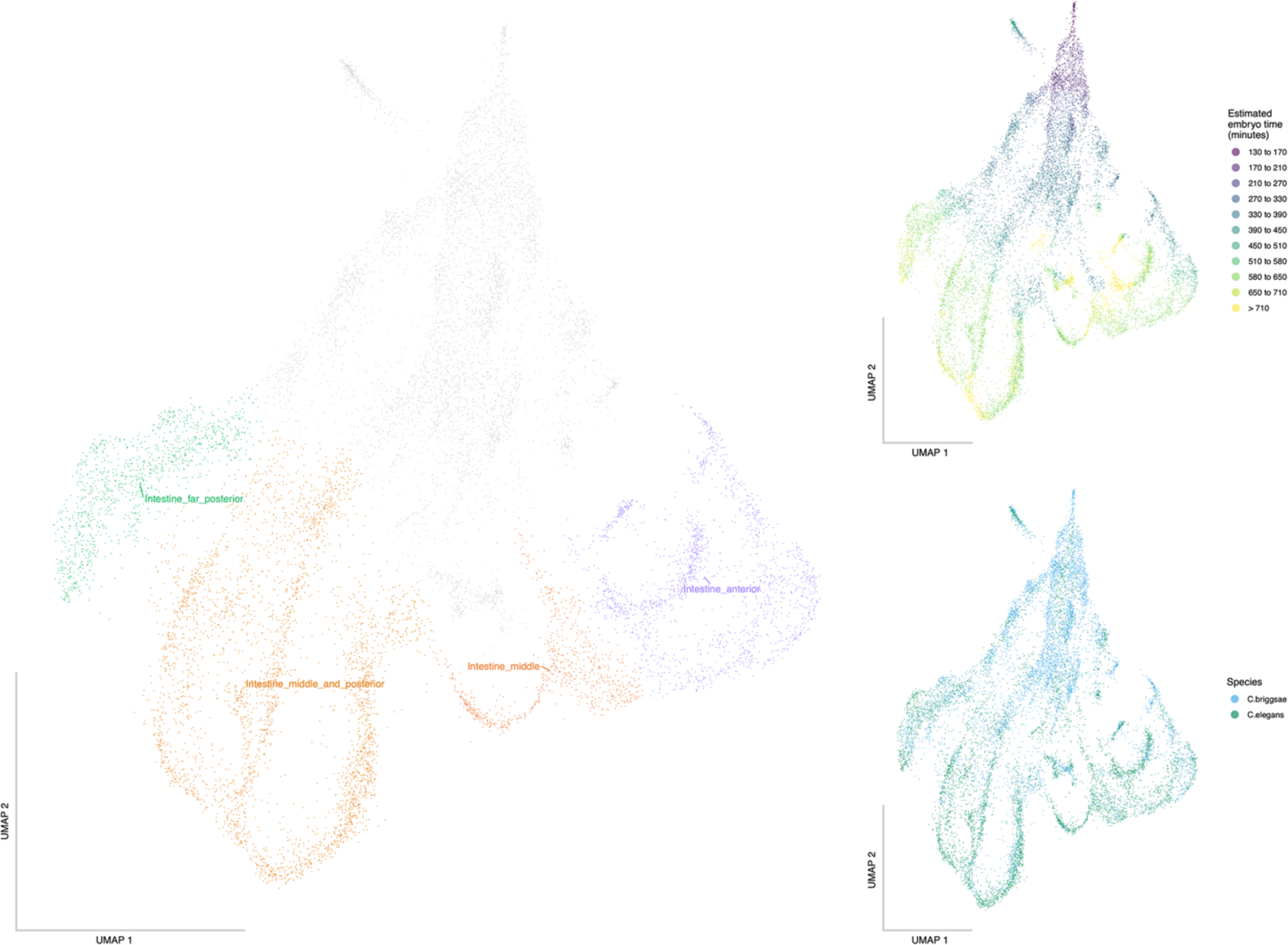
These plots show the intestine cell class subset integrated UMAPs. The cell labels (left), estimated embryo time (top-right), and the species composition (bottom-right) are shown.

**Supp. Figure 10.**
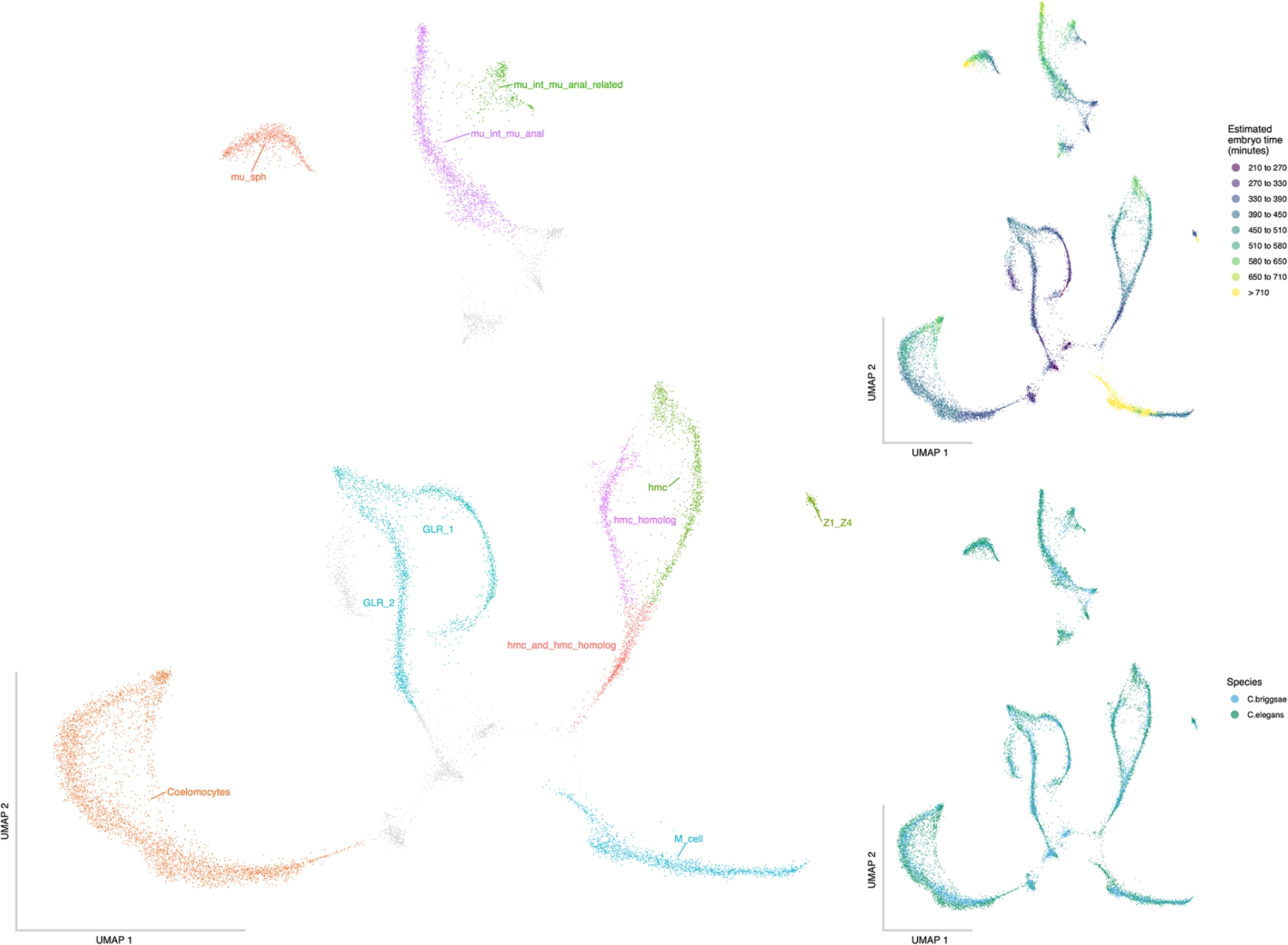
These plots show the mesoderm cell class subset integrated UMAPs. The cell labels (left), estimated embryo time (top-right), and the species composition (bottom-right) are shown.

**Supp. Figure 11.**
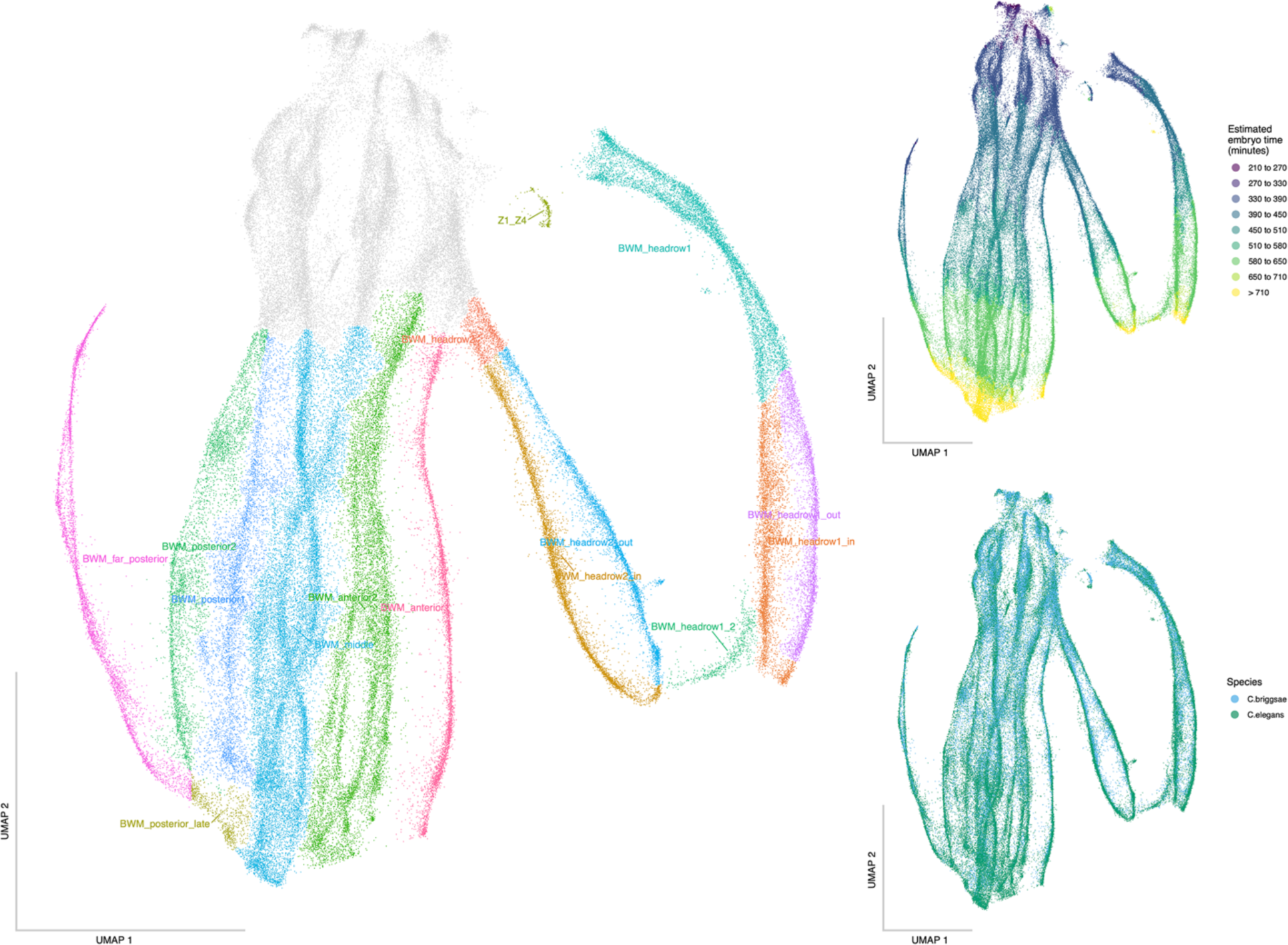
These plots show the muscle cell class subset integrated UMAPs. The cell labels (left), estimated embryo time (top-right), and the species composition (bottom-right) are shown.

**Supp. Figure 12.**
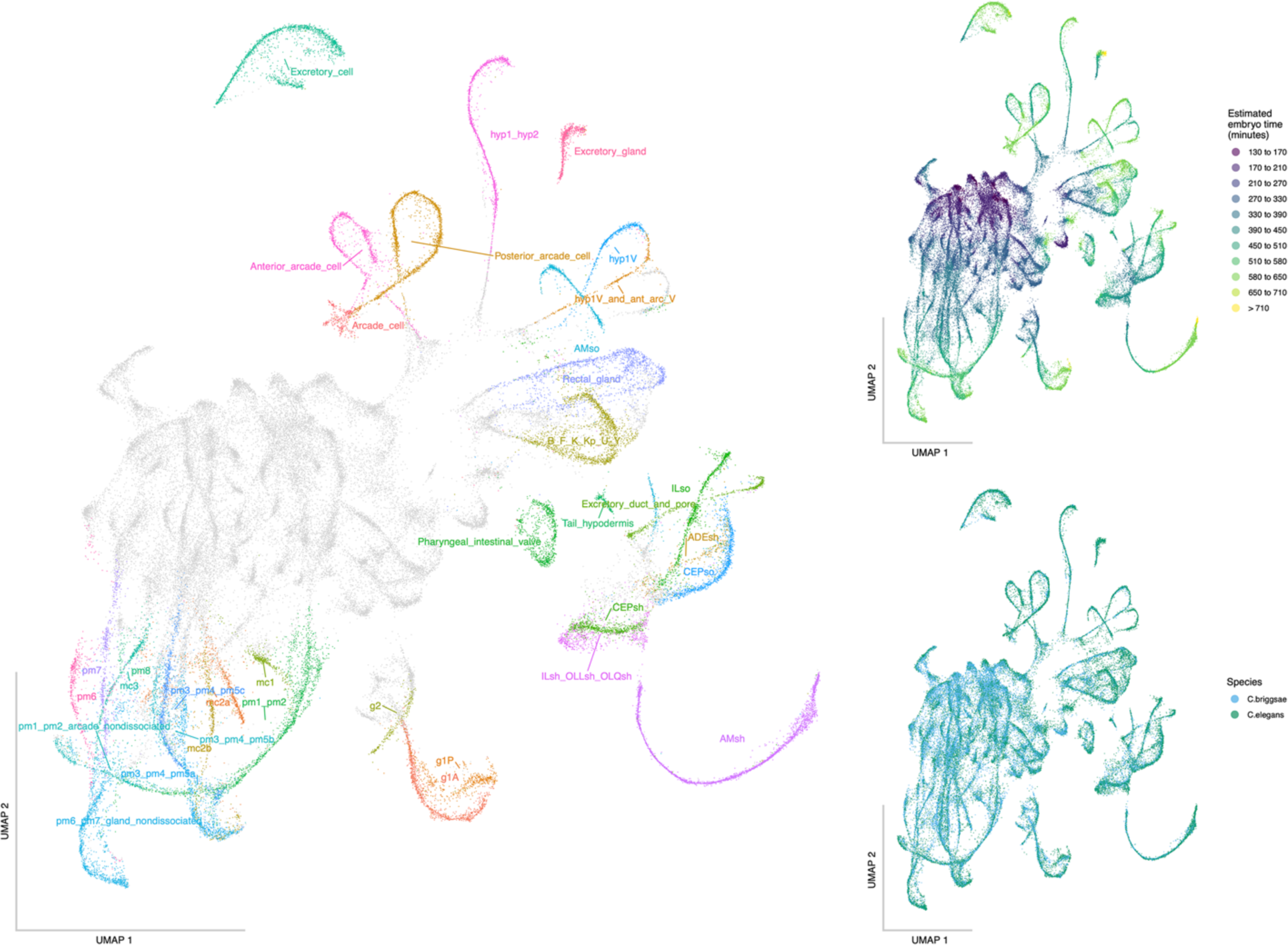
These plots show the pharynx, glia and excretory cell class subset integrated UMAPs. The cell labels (left), estimated embryo time (top-right), and the species composition (bottom-right) are shown.

**Supp. Figure 13.**
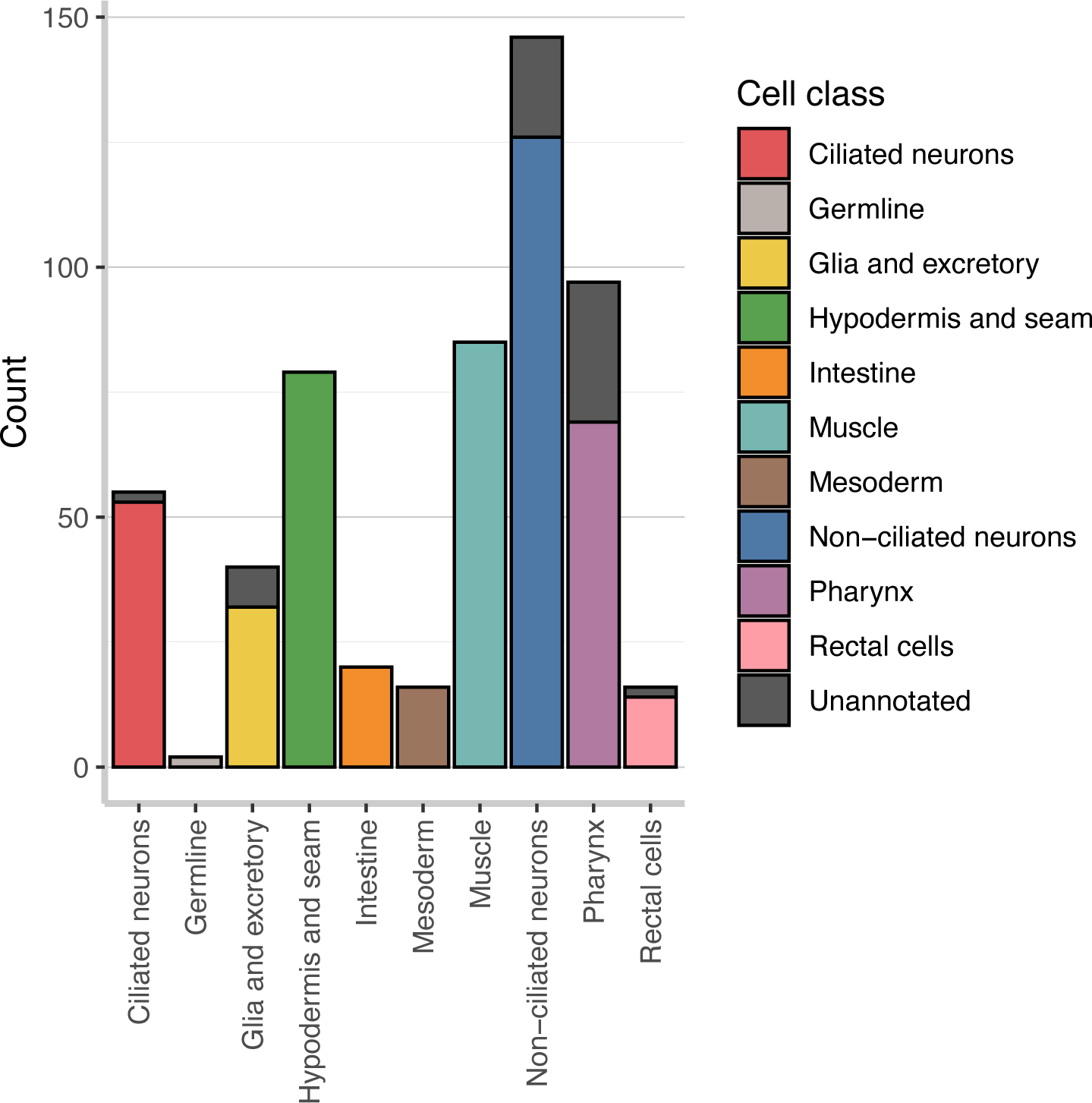
The number of terminal cell types that were annotated is shown by their respective cell class color. The number of unannotated terminal cell types is shown with a dark gray color. Overall, most terminal cell types were annotated with the exception of the late dividing and pharyngeal neurons.

**Supp. Figure 14.**
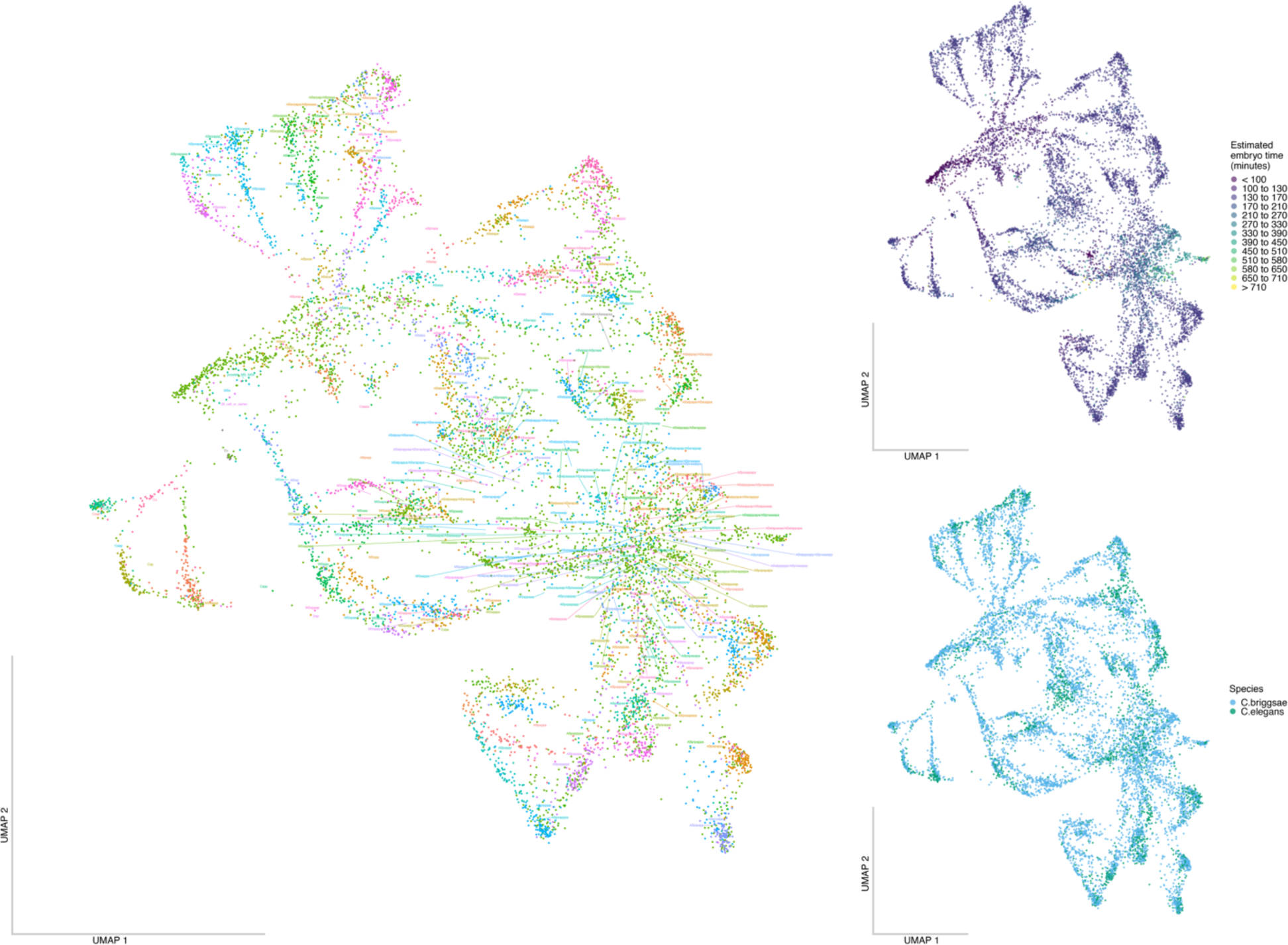
These plots show the cells that are within a 0 to 150 estimated embryo time subset in integrated UMAPs. The cell labels (left), estimated embryo time (top-right), and the species composition (bottom-right) are shown. The cell label position is based on the centroid of the labeled cells and can occasionally be shifted relative to the labeled cells position.

**Supp. Figure 15.**
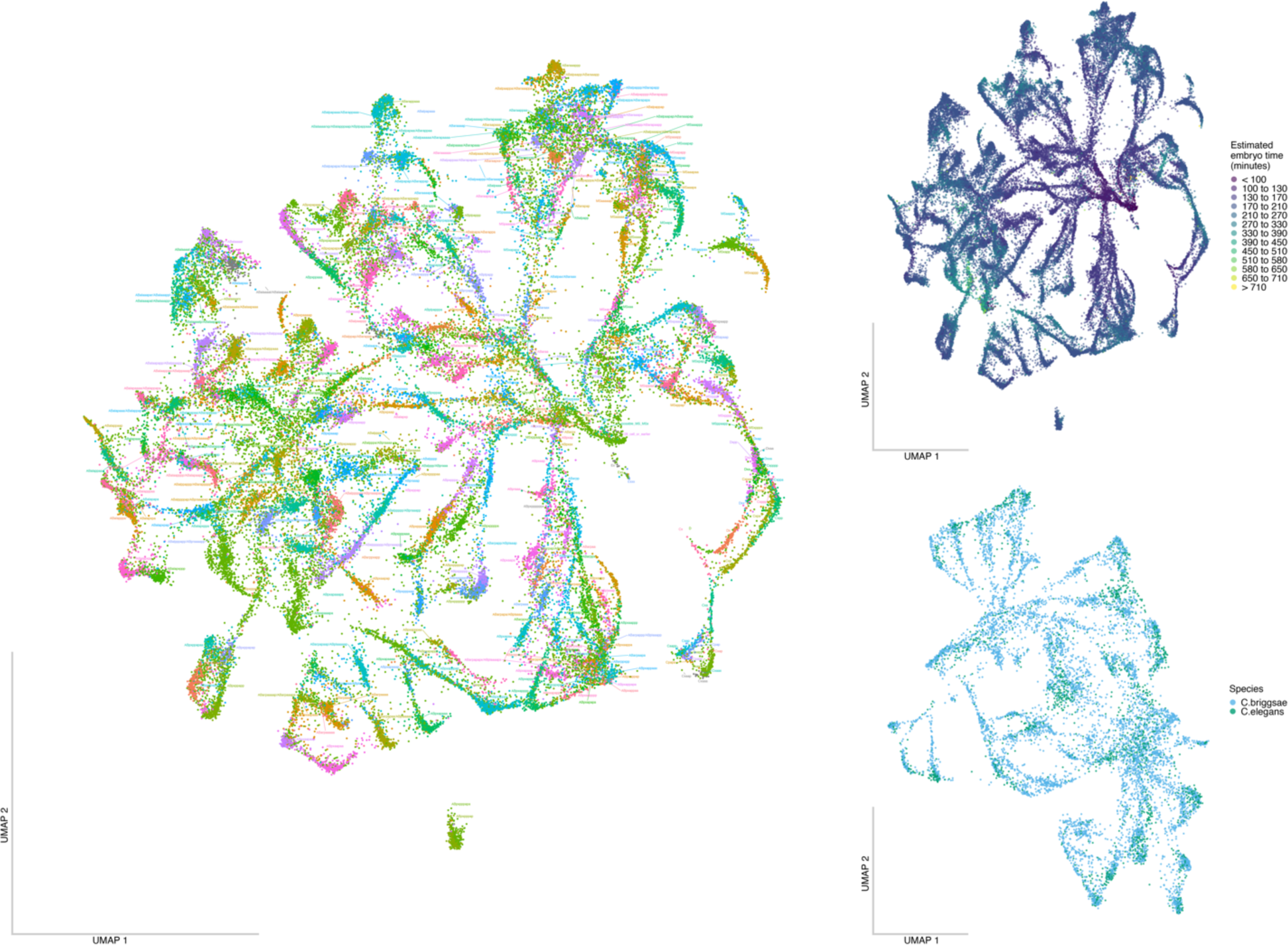
These plots show the cells that are within a 0 to 200 estimated embryo time subset in integrated UMAPs. The cell labels (left), estimated embryo time (top-right), and the species composition (bottom-right) are shown. The cell label position is based on the centroid of the labeled cells and can occasionally be shifted relative to the labeled cells position.

**Supp. Figure 16.**
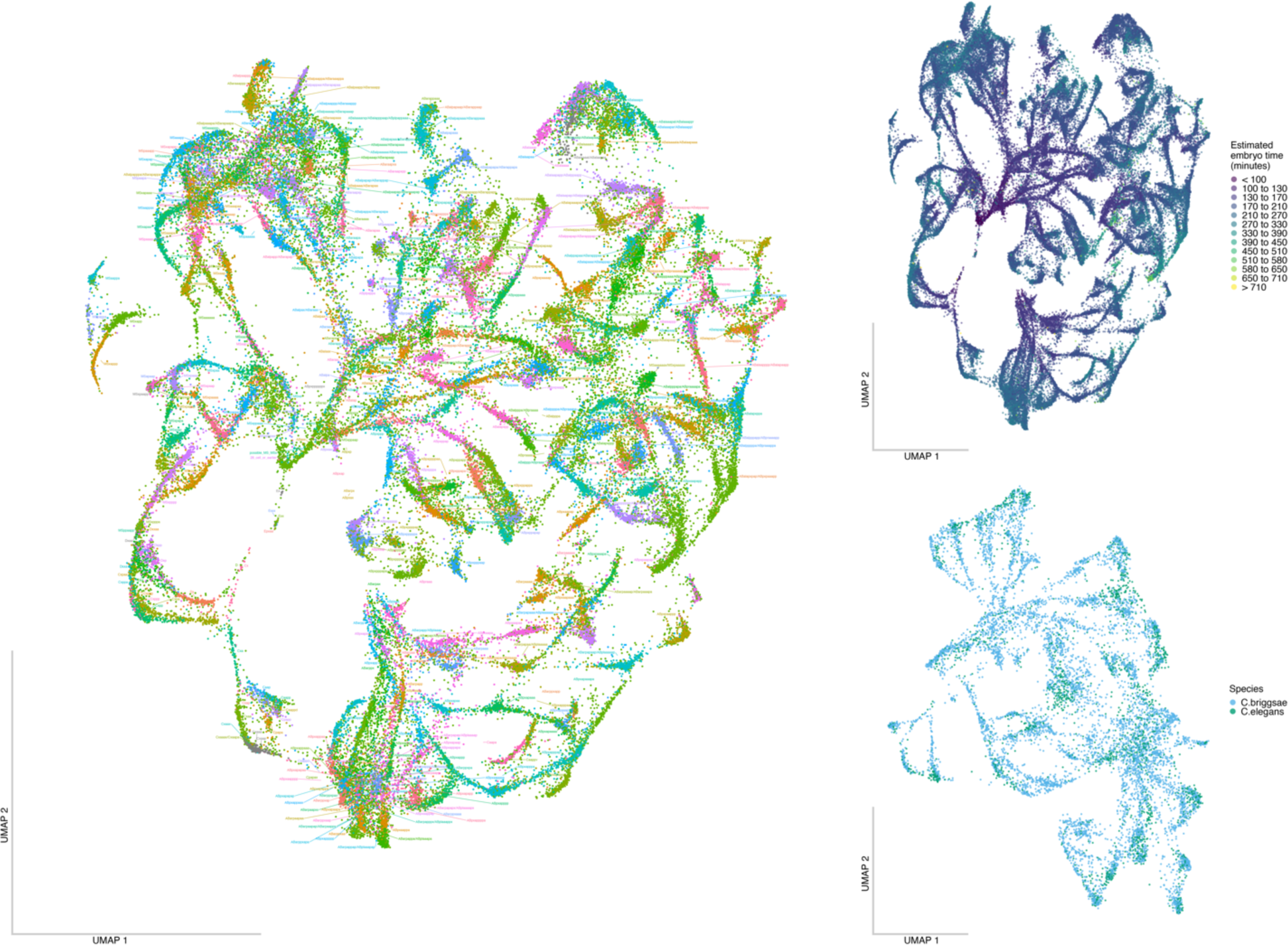
These plots show the cells that are within a 0 to 250 estimated embryo time subset in integrated UMAPs. The cell labels (left), estimated embryo time (top-right), and the species composition (bottom-right) are shown. The cell label position is based on the centroid of the labeled cells and can occasionally be shifted relative to the labeled cells position.

**Supp. Figure 17.**
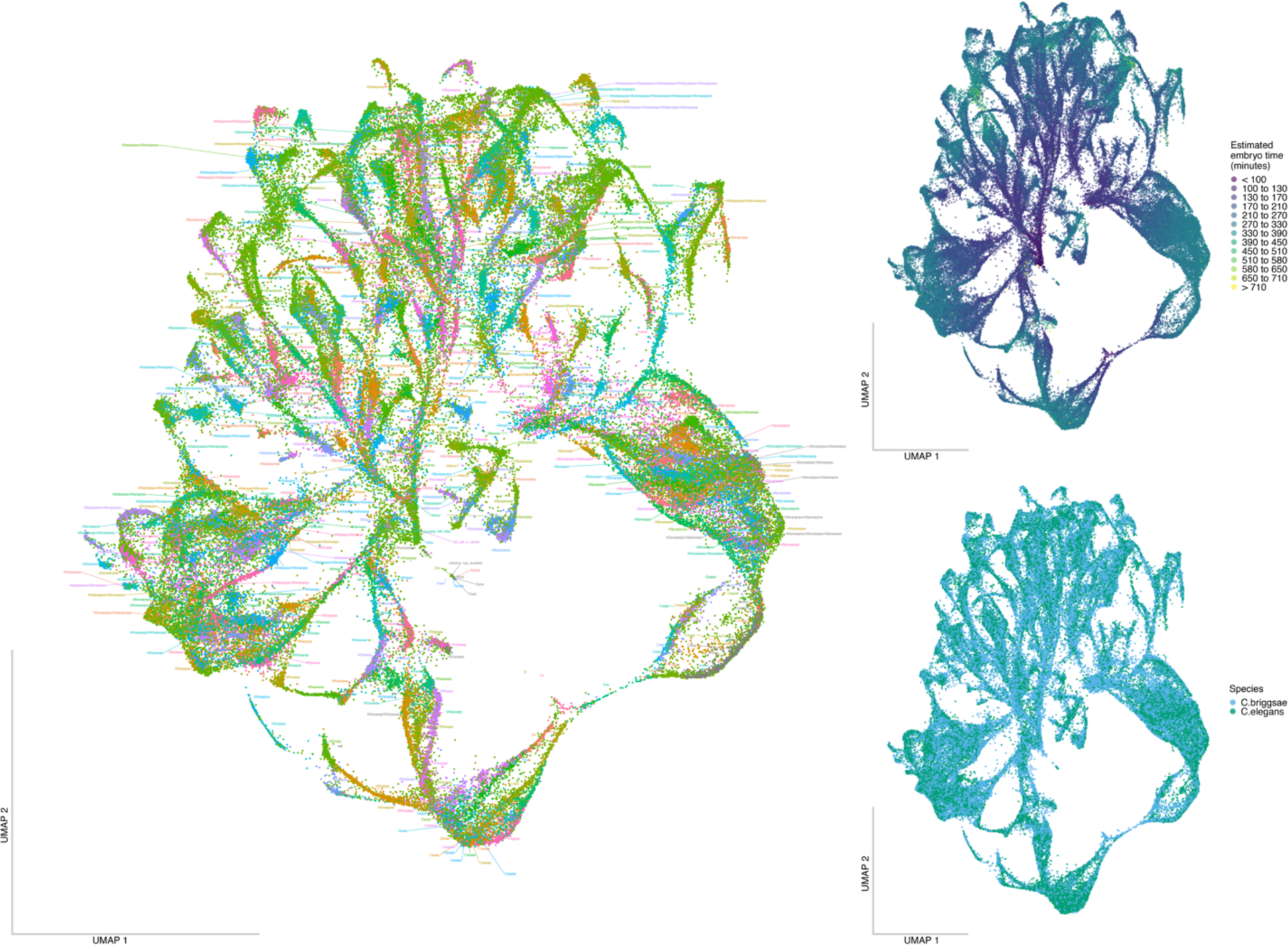
These plots show the cells that are within a 0 to 300 estimated embryo time subset in integrated UMAPs. The cell labels (left), estimated embryo time (top-right), and the species composition (bottom-right) are shown. The cell label position is based on the centroid of the labeled cells and can occasionally be shifted relative to the labeled cells position.

**Supp. Figure 18.**
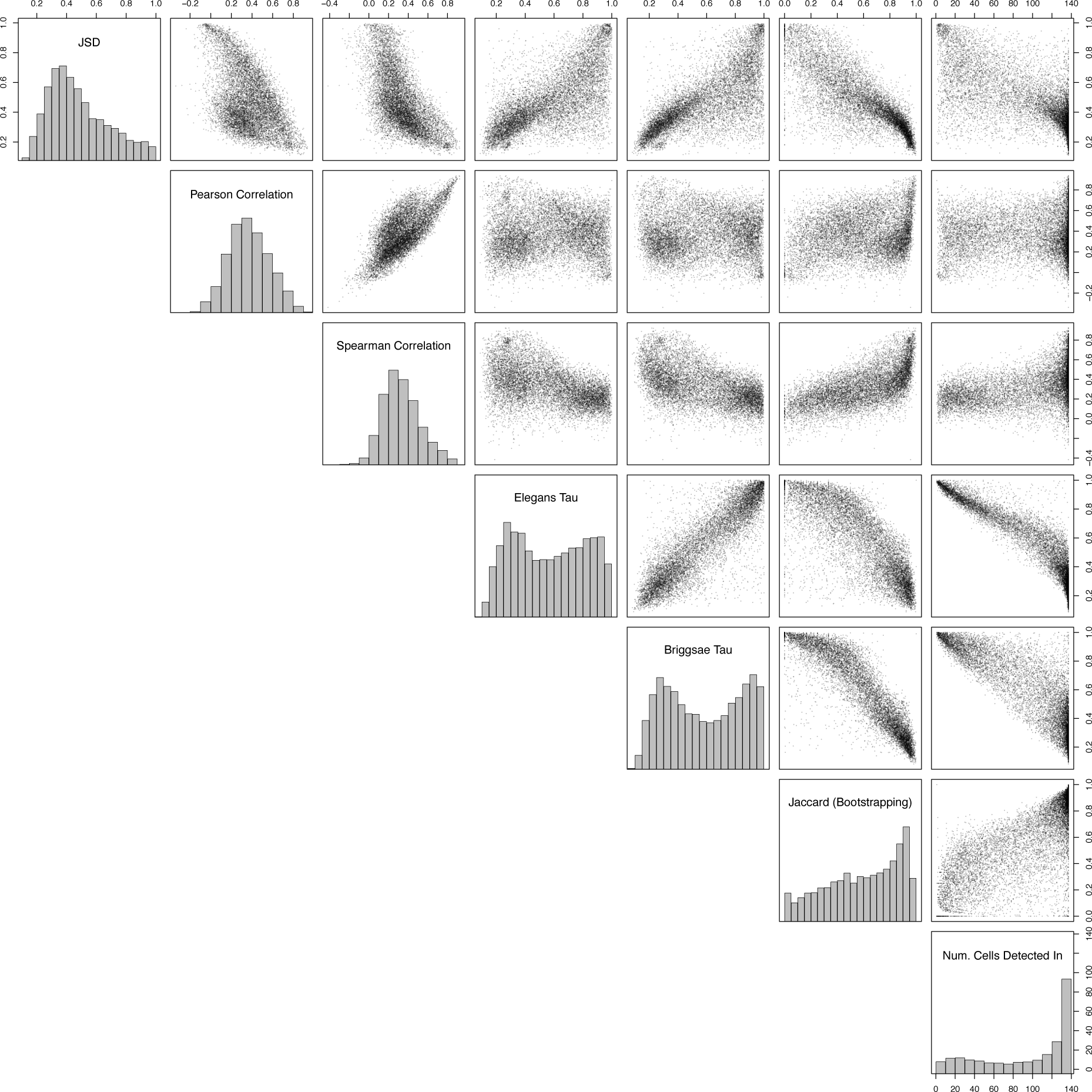
The relationship between various gene metrics calculated on the pseudo bulked terminal cell-types is shown with a histogram of its distribution shown on the diagonal and the relationship between metrics shown on the off-diagonal. The gene expression pattern distance (JSD_gene_), gene Pearson correlation, gene Spearman correlation, *C. elegans Tau*, *C. briggsae Tau*, Jaccard distance (calculated on how many cell types was that gene expressed in common between the species using the 95% TPM confidence interval cutoff to binarize the expression data), and the number of cell types a gene was detected in are shown.

**Supp. Figure 19.**
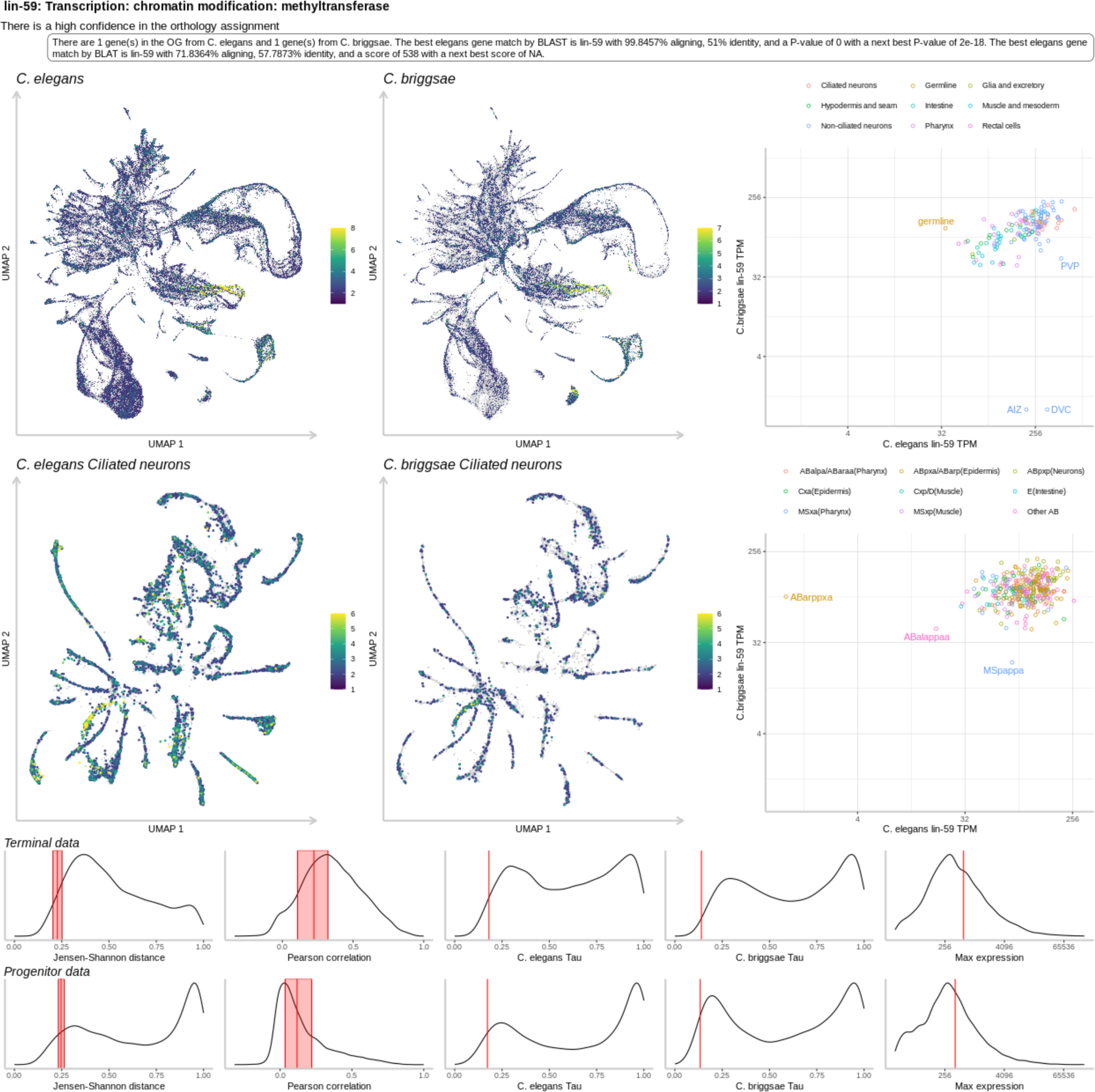
An example gene page for the *lin-59* gene. Shown in the top of the gene page is a global UMAP showing the expression of the gene of interest, a cell subset UMAP showing the expression of the gene of interest, the terminal cell-type comparative TPM values shown in log2 space, and the progenitor cell-type comparative TPM values shown in log2 space. The progenitors are summarized by their general lineage type. Shown in the bottom of the gene page is a bunch of gene metrics, where the values for this gene is shown in red as a confidence interval (CI) range on top of the dataset wide distribution. These metrics are shown for the terminal (top) and progenitor (bottom) cell-types. The metrics shown are the gene expression pattern distance shown as the JSD_gene_ calculated on the bootstrapped TPM values, the gene expression pattern distance shown as the Pearson correlation coefficient calculated on the bootstrapped TPM values, the broadness of gene expression pattern shown as the *Tau* value for *C. elegans*, the broadness of gene expression pattern shown as the *Tau* value for *C. briggsae* and the maximum TPM value across any cell-type in both species. More gene examples can be found on the associated github page.

**Supp. Figure 20.**
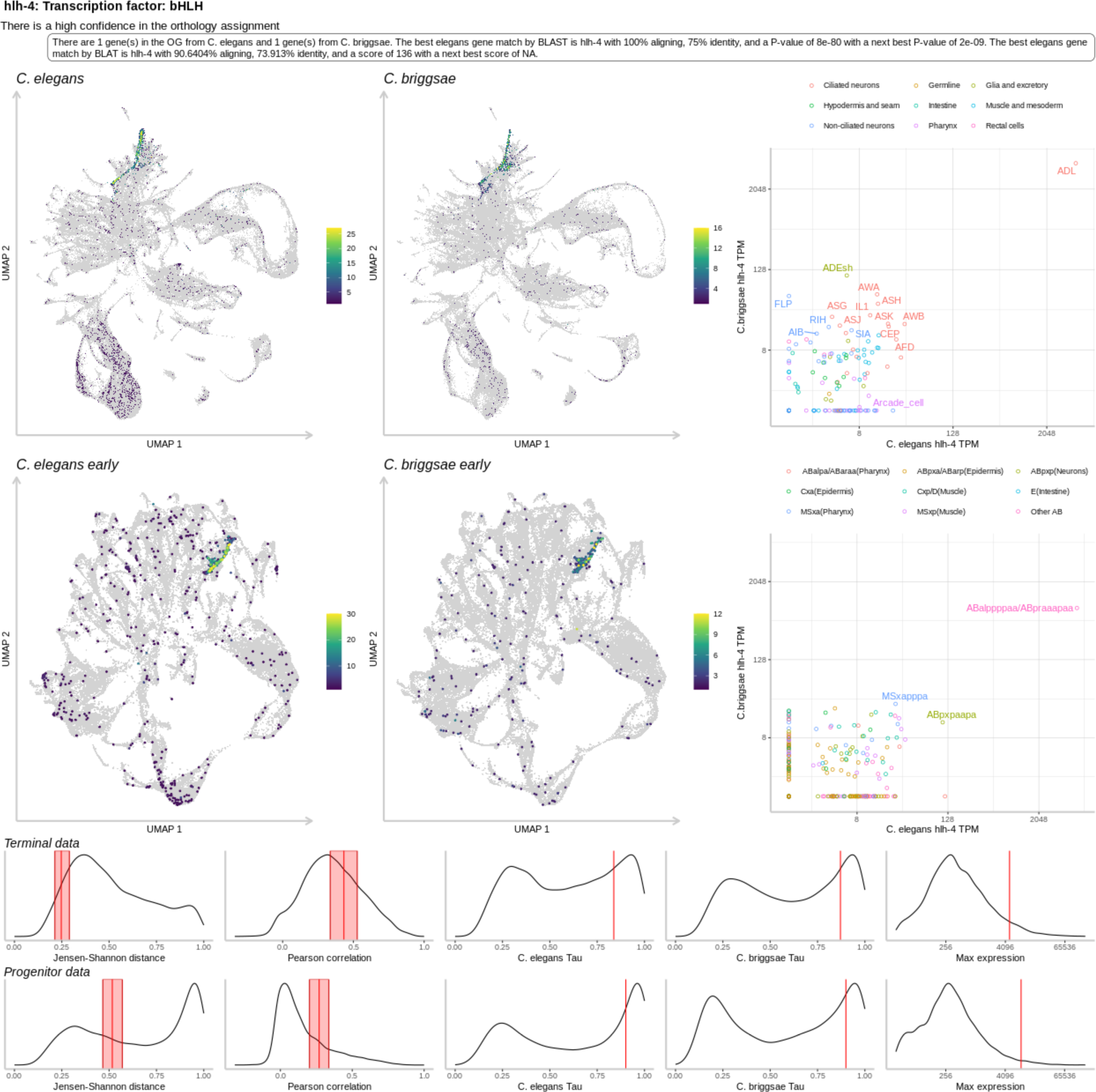
An example gene page for the *hlh-4* gene. Shown in the top of the gene page is a global UMAP showing the expression of the gene of interest, a cell subset UMAP showing the expression of the gene of interest, the terminal cell-type comparative TPM values shown in log2 space, and the progenitor cell-type comparative TPM values shown in log2 space. The progenitors are summarized by their general lineage type. Shown in the bottom of the gene page is a bunch of gene metrics, where the values for this gene is shown in red as a confidence interval (CI) range on top of the dataset wide distribution. These metrics are shown for the terminal (top) and progenitor (bottom) cell-types. The metrics shown are the gene expression pattern distance shown as the JSD_gene_ calculated on the bootstrapped TPM values, the gene expression pattern distance shown as the Pearson correlation coefficient calculated on the bootstrapped TPM values, the broadness of gene expression pattern shown as the *Tau* value for *C. elegans*, the broadness of gene expression pattern shown as the *Tau* value for *C. briggsae* and the maximum TPM value across any cell-type in both species. More gene examples can be found on the associated github page.

**Supp. Figure 21.**
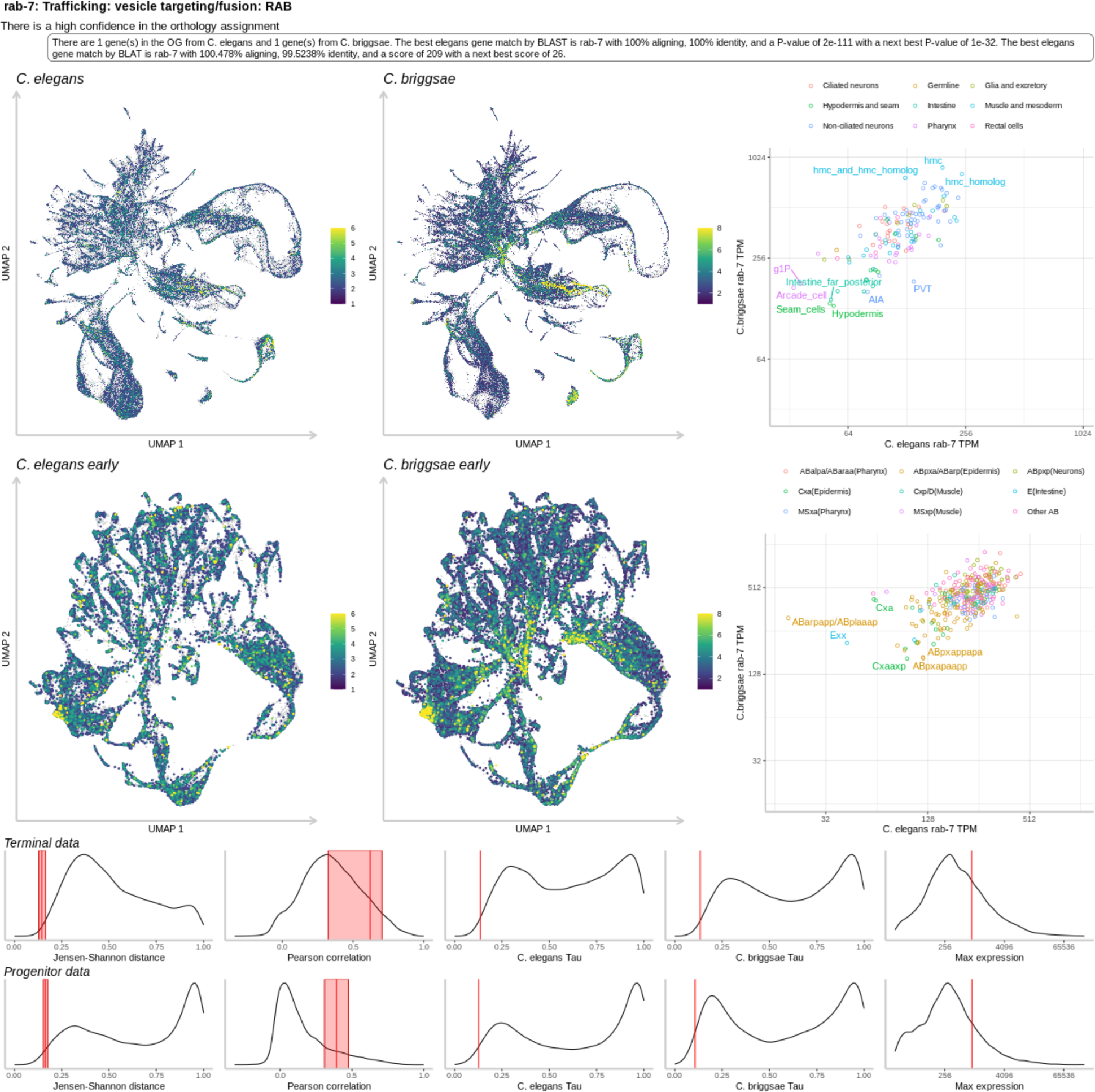
An example gene page for the *rab-7* gene. Shown in the top of the gene page is a global UMAP showing the expression of the gene of interest, a cell subset UMAP showing the expression of the gene of interest, the terminal cell-type comparative TPM values shown in log2 space, and the progenitor cell-type comparative TPM values shown in log2 space. The progenitors are summarized by their general lineage type. Shown in the bottom of the gene page is a bunch of gene metrics, where the values for this gene is shown in red as a confidence interval (CI) range on top of the dataset wide distribution. These metrics are shown for the terminal (top) and progenitor (bottom) cell-types. The metrics shown are the gene expression pattern distance shown as the JSD_gene_ calculated on the bootstrapped TPM values, the gene expression pattern distance shown as the Pearson correlation coefficient calculated on the bootstrapped TPM values, the broadness of gene expression pattern shown as the *Tau* value for *C. elegans*, the broadness of gene expression pattern shown as the *Tau* value for *C. briggsae* and the maximum TPM value across any cell-type in both species. More gene examples can be found on the associated github page.

**Supp. Figure 22.**
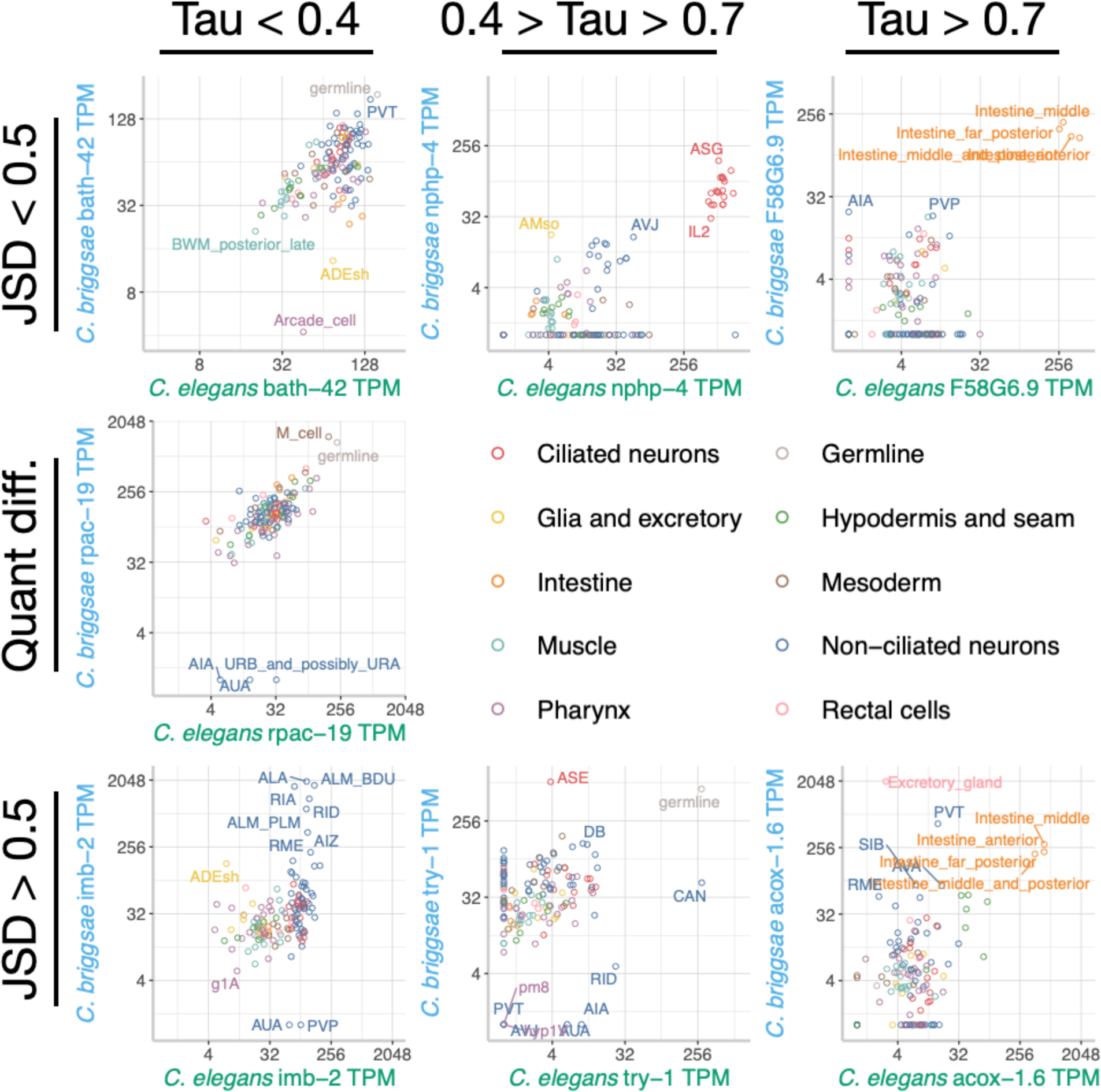
Example gene expression TPM values across the terminal cells between *C. elegans* and *C. briggsae* for the conserved, diverged, and broadness categories. These gene examples are representative of the overall categories.

**Supp. Figure 23.**
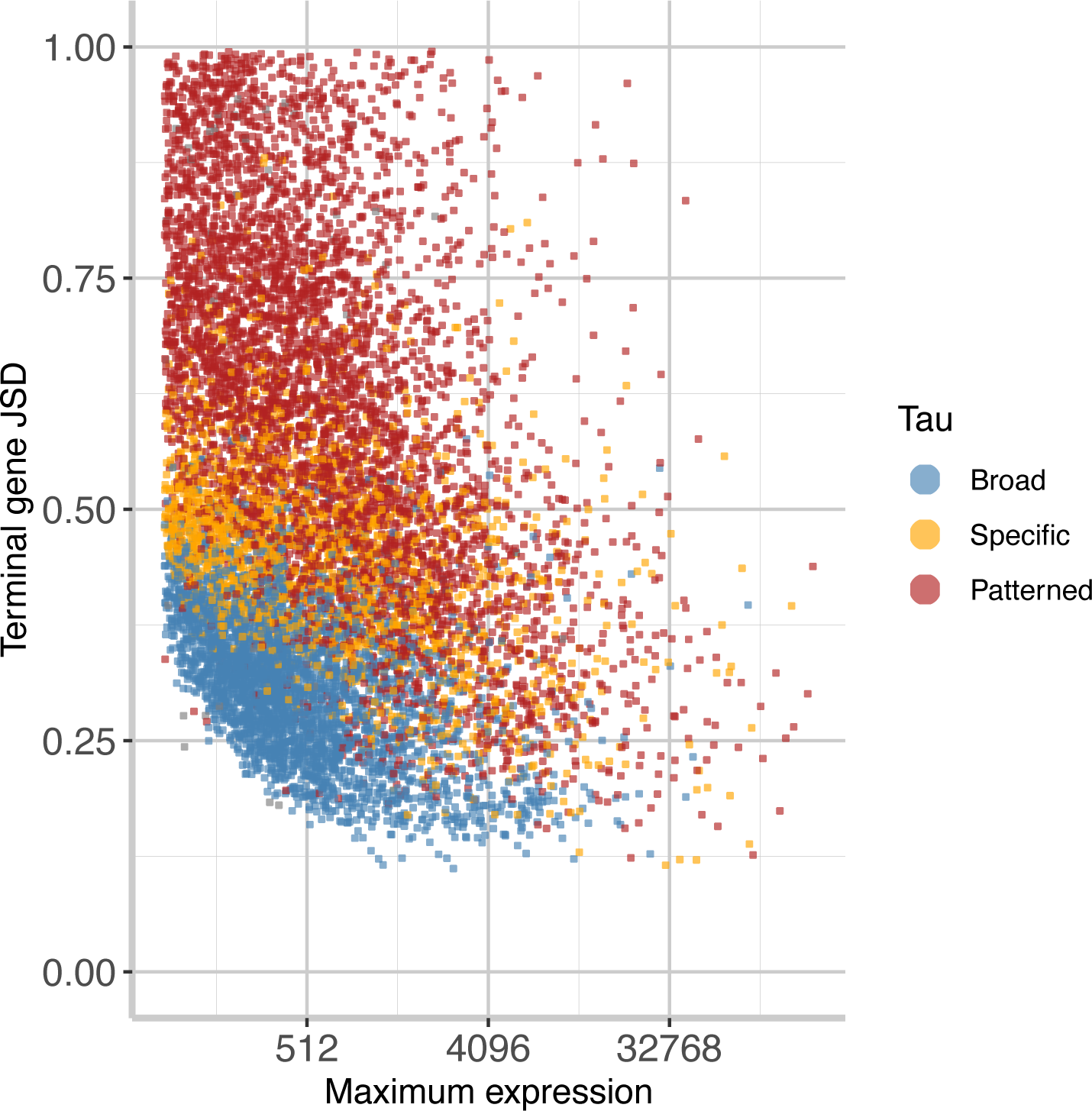
A plot showing the maximum expression value of every gene across the terminal cell types versus the terminal JSD_gene_ with the gene broadness category shown as a color. This plot generally shows that more broadly expressed genes are generally more conserved.

**Supp. Figure 24.**
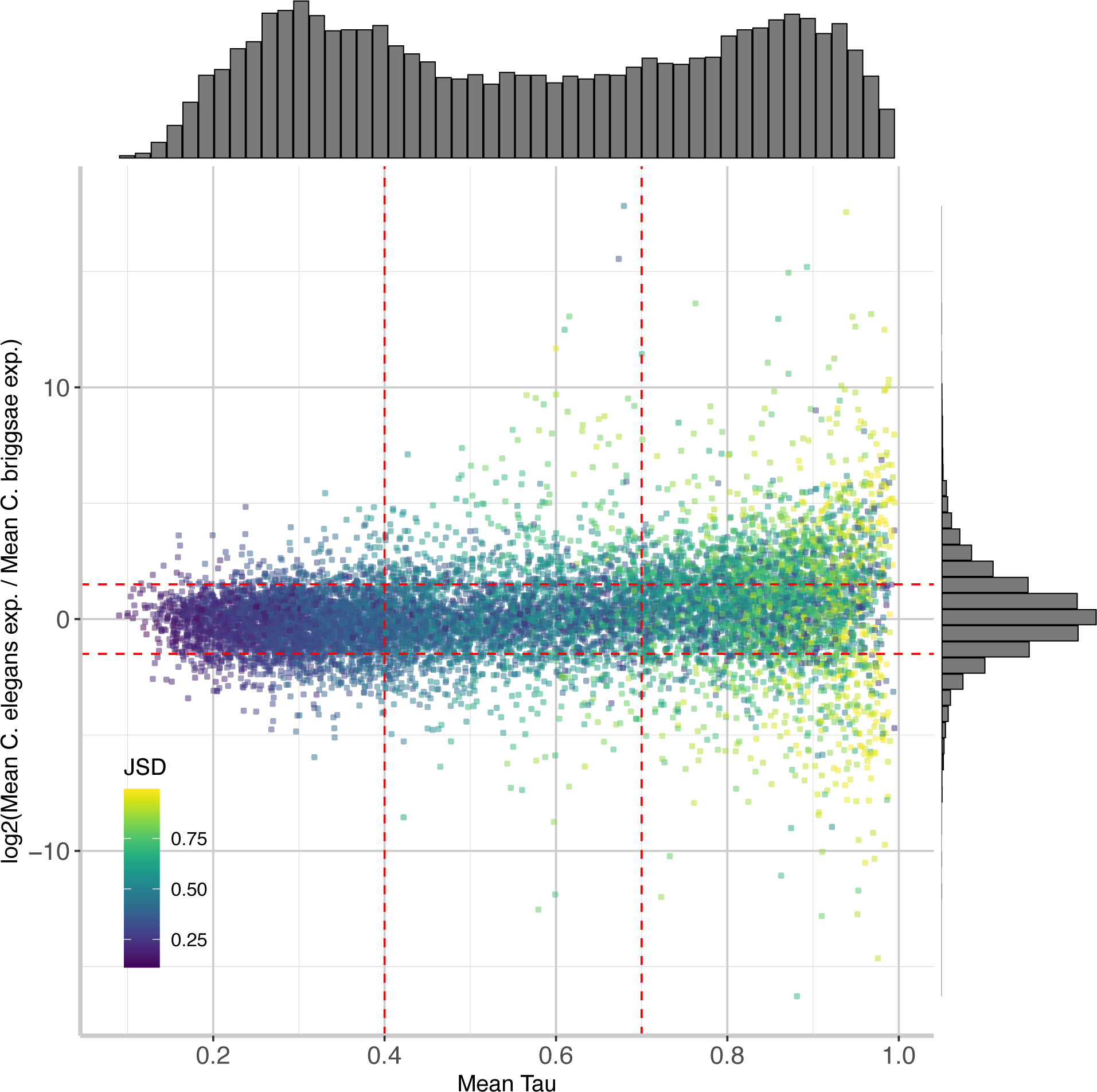
The mean gene expression broadness (Tau) versus the log2 ratio of the mean gene expression value across all terminal cell types shows the relative even distribution of differential gene expression. The horizontal red lines indicate a value of 1.5 and the vertical lines indicate whether the gene has broad, patterned, or specific expression. The color indicates the gene expression distance across the terminal cell types.

**Supp. Figure 25.**
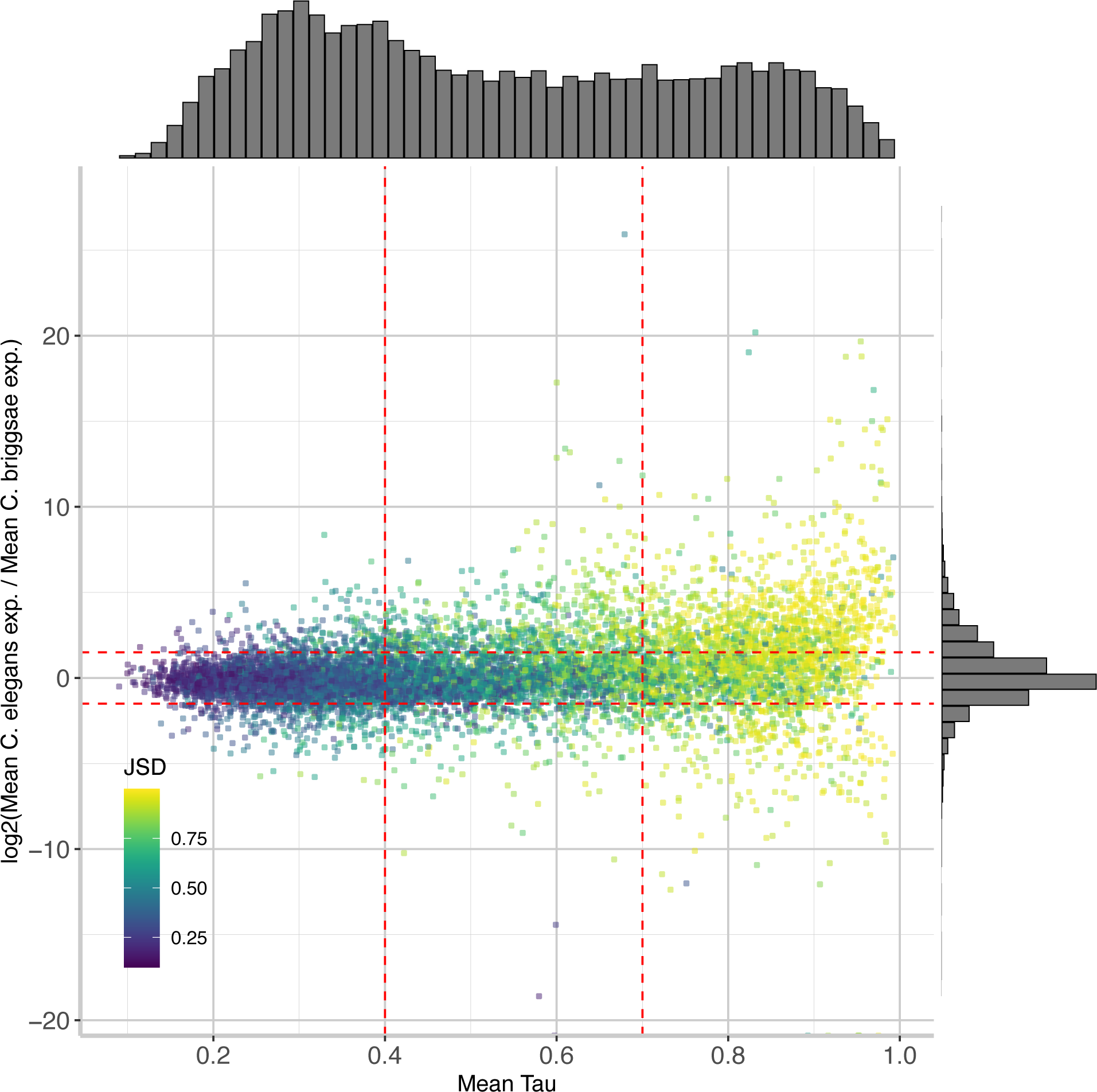
The mean gene expression broadness (Tau) versus the log2 ratio of the mean gene expression value across all progenitors shows the relative even distribution of differential gene expression. The horizontal red lines indicate a value of 1.5 and the vertical lines indicate whether the gene has broad, patterned, or specific expression. The color indicates the gene expression distance across the progenitors.

**Supp. Figure 26.**
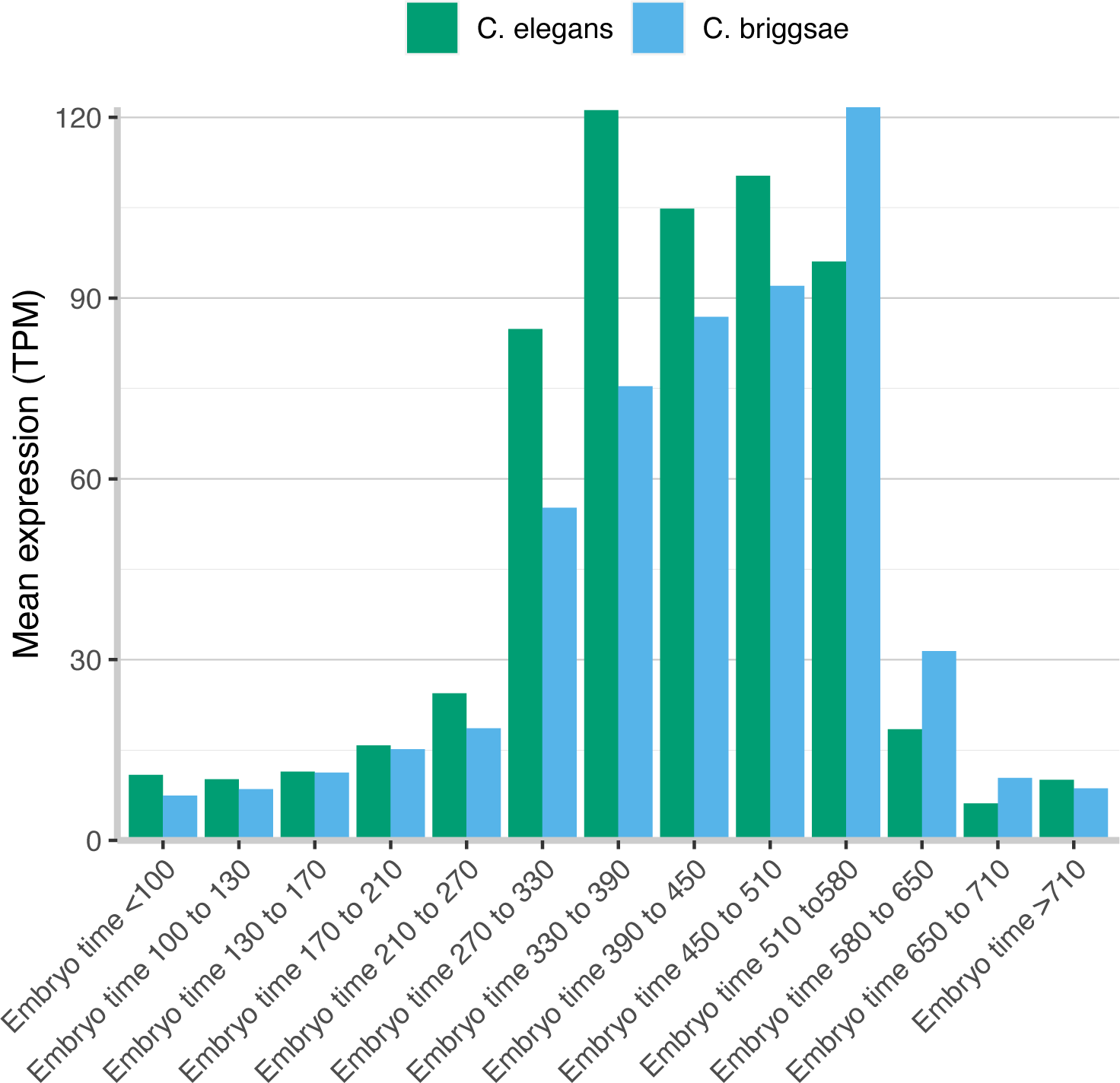
The mean TPM expression value of genes inside of the cilial program (genes in the cilia WormCategory) for *C. elegans* and *C. briggsae* across the embryo time bins shows a later time bias in the onset of the cilia program for *C. briggsae*.

**Supp. Figure 27.**
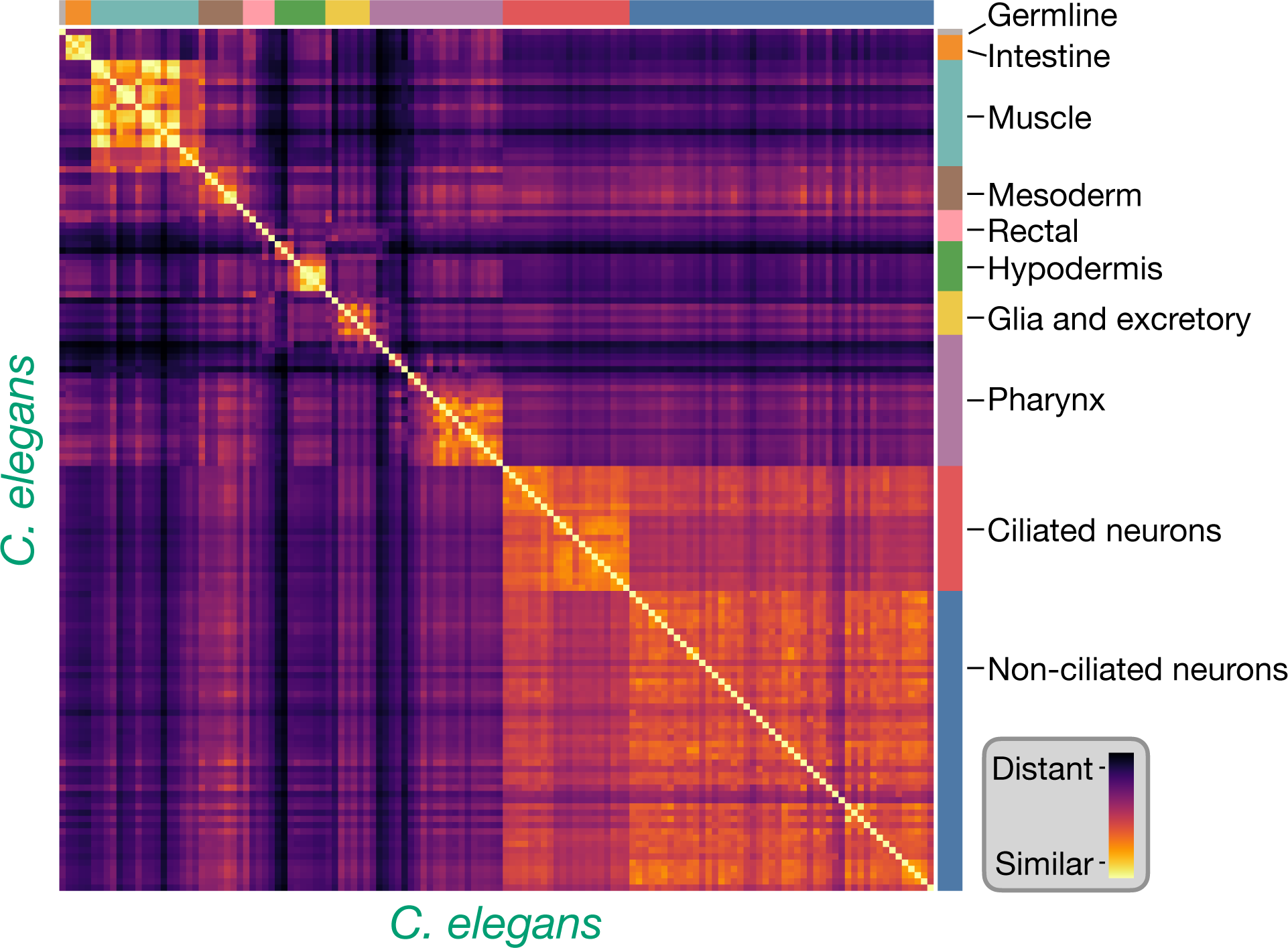
A JSD_cell_ matrix between cell types from *C. elegans*. Cell types of the same cell class tend to have similarity versus unrelated cell types similar to the within species comparisons.

**Supp. Figure 28.**
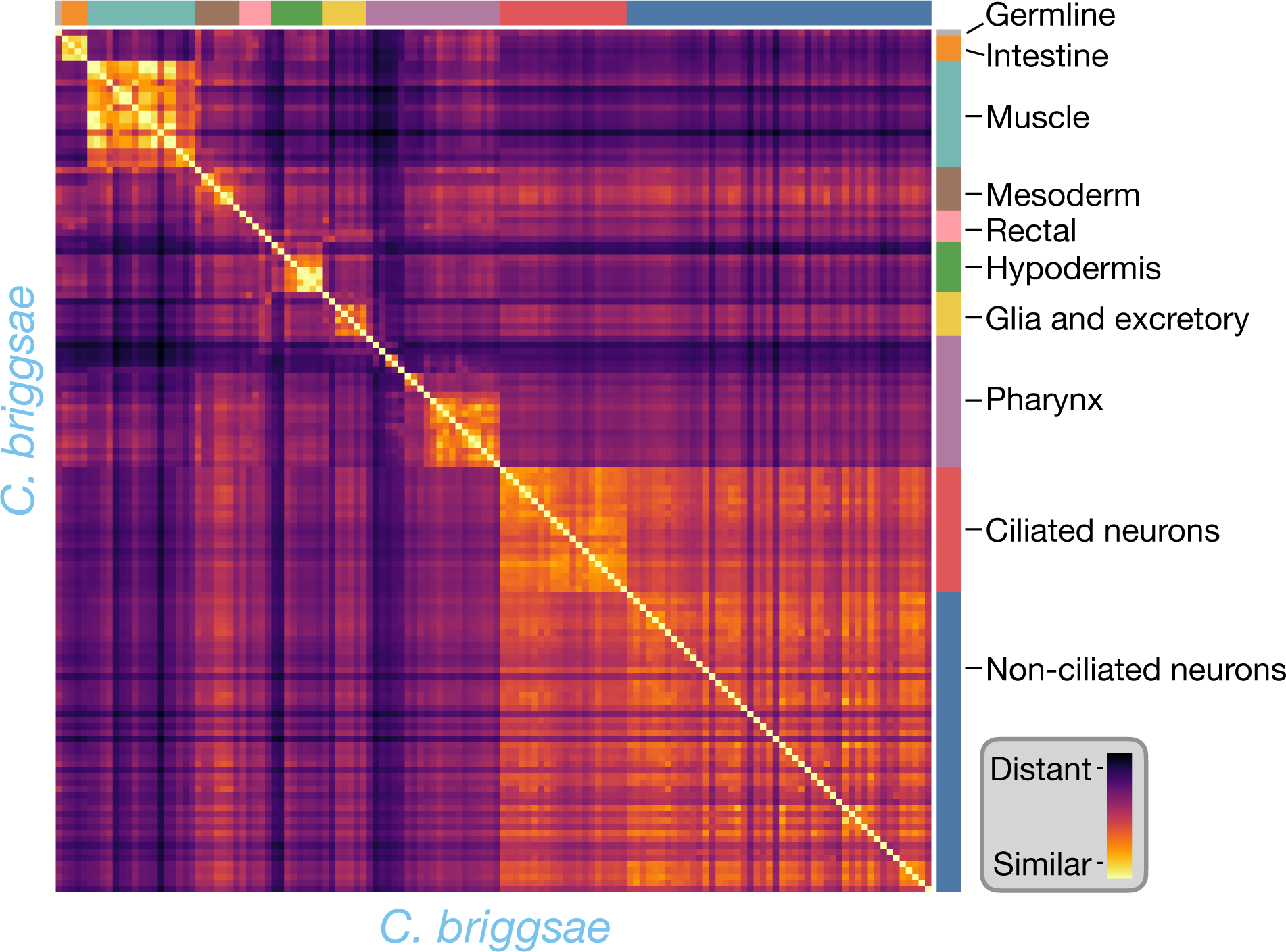
A JSD_cell_ matrix between cell types from *C. briggsae*. Cell types of the same cell class tend to have similarity versus unrelated cell types similar to the within species comparisons.

**Supp. Figure 29.**
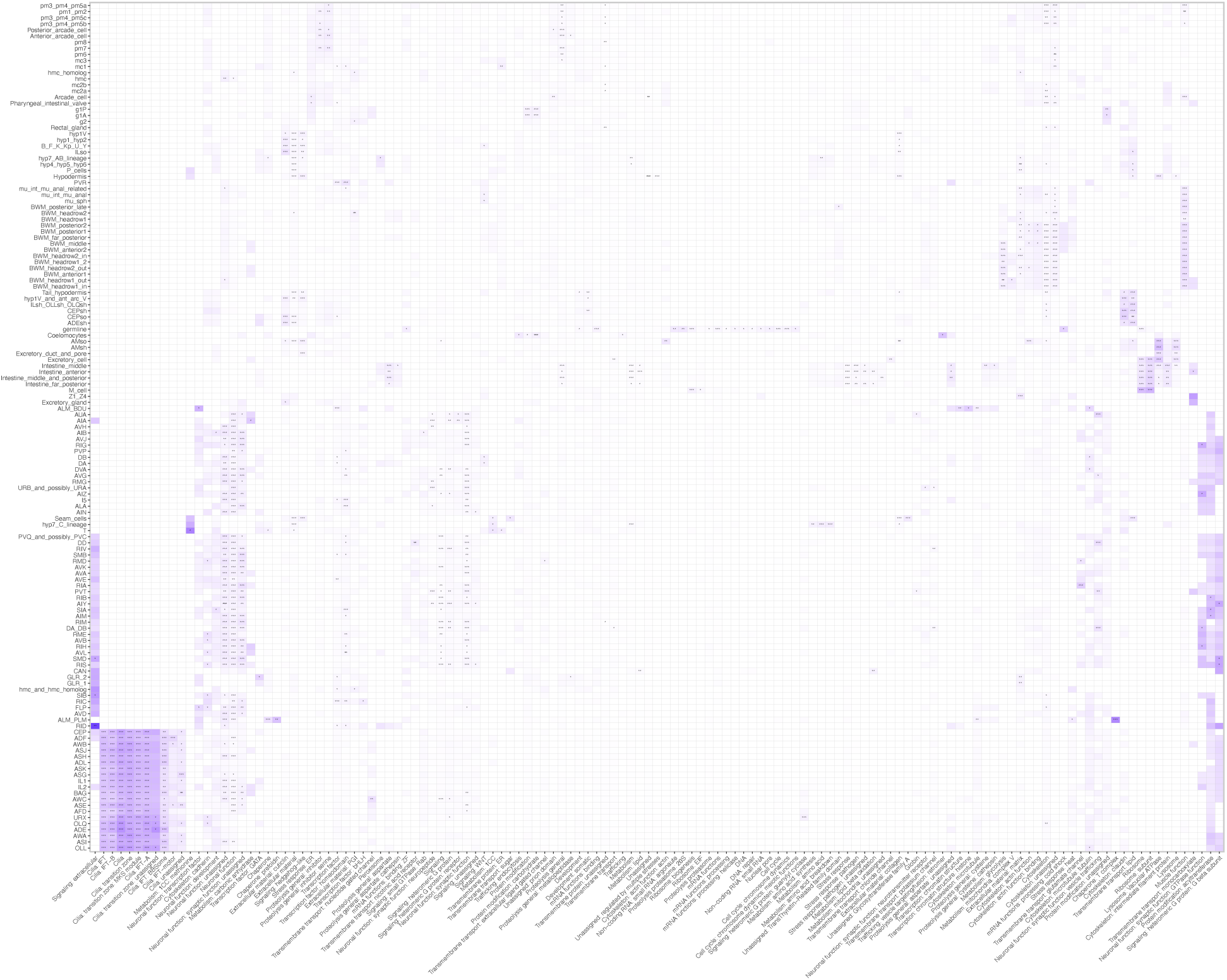
The enrichment of the WormCat category annotations for the shared gene markers of the terminal cell types between *C. elegans* and *C. briggsae*. The stars represent Bonferroni corrected p-values of a Fisher’s Exact Test where * is p <0.05, ** is p <0.005, and *** is p <0.0005.

**Supp. Figure 30.**
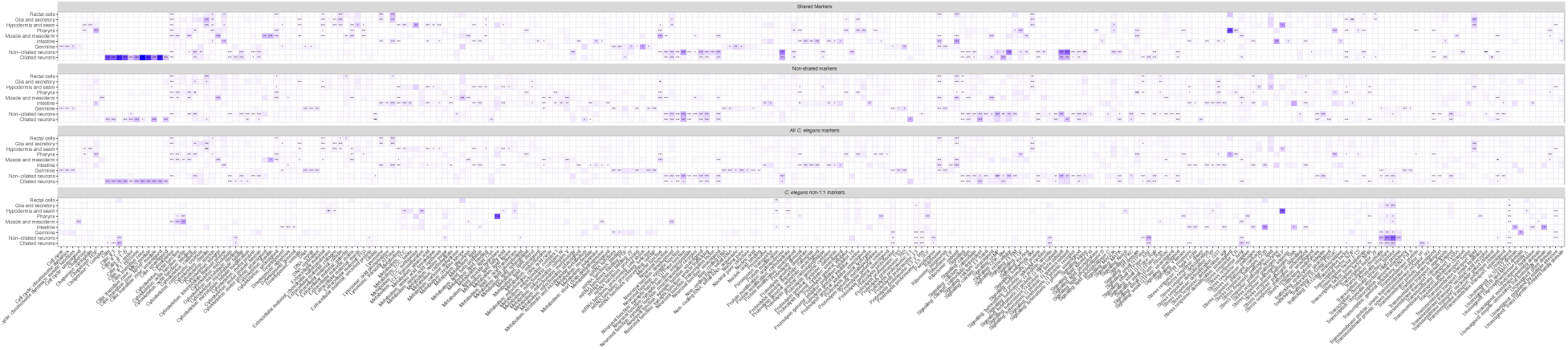
The enrichment of the WormCat category annotations for the shared, non-shared, private, and all ^gene^ markers of the terminal cell types for *C. elegans*, abstracted into cell classes. The stars represent Bonferroni corrected p-values of a Fisher’s Exact Test where * is p <0.05, ** is p <0.005, and *** is p <0.0005.

**Supp. Figure 31.**
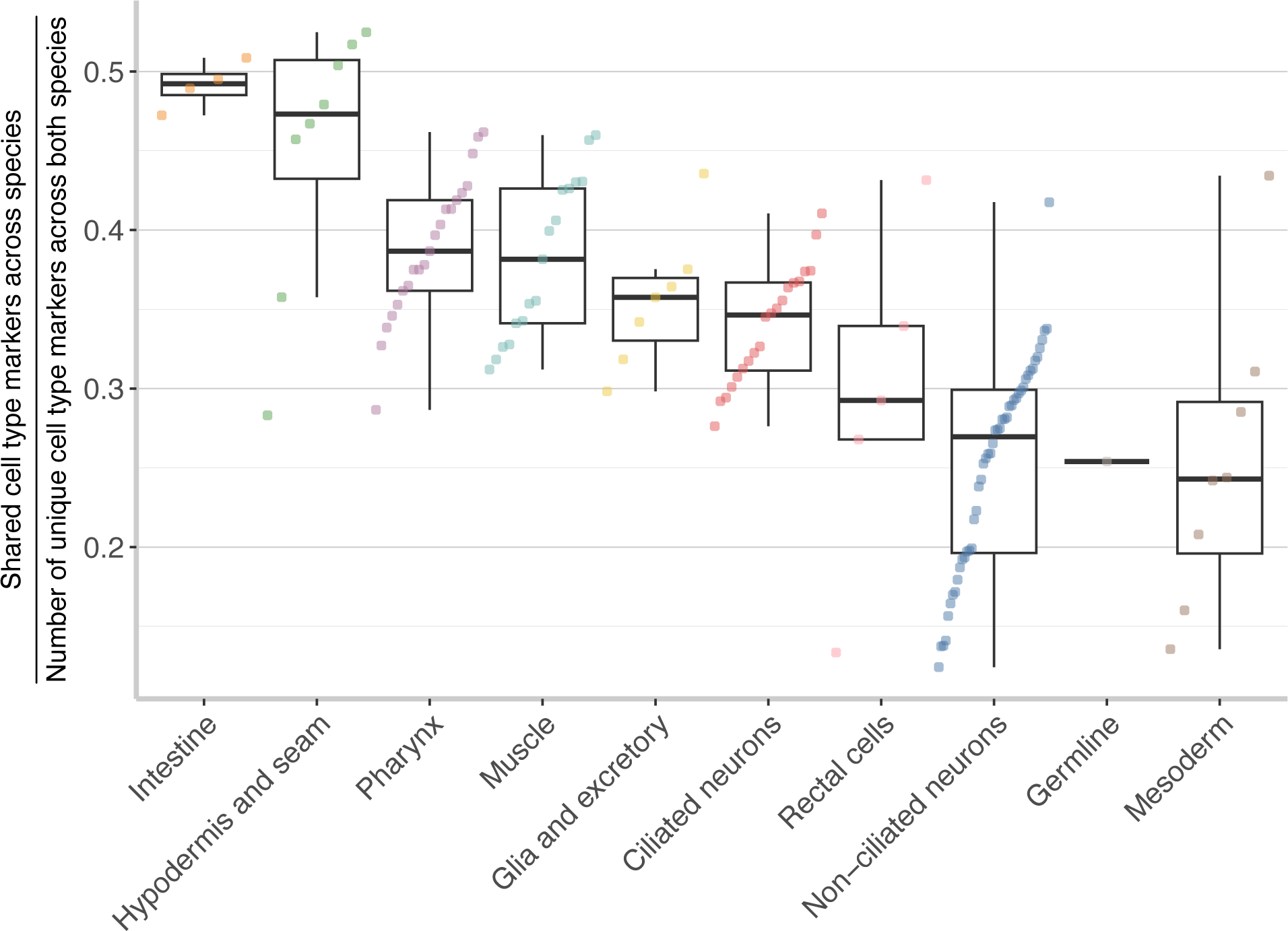
The degree of marker sharing across terminal cell types shows a similar rank ordering with the exception of the mesoderm and germline cell classes.

**Supp. Figure 32.**
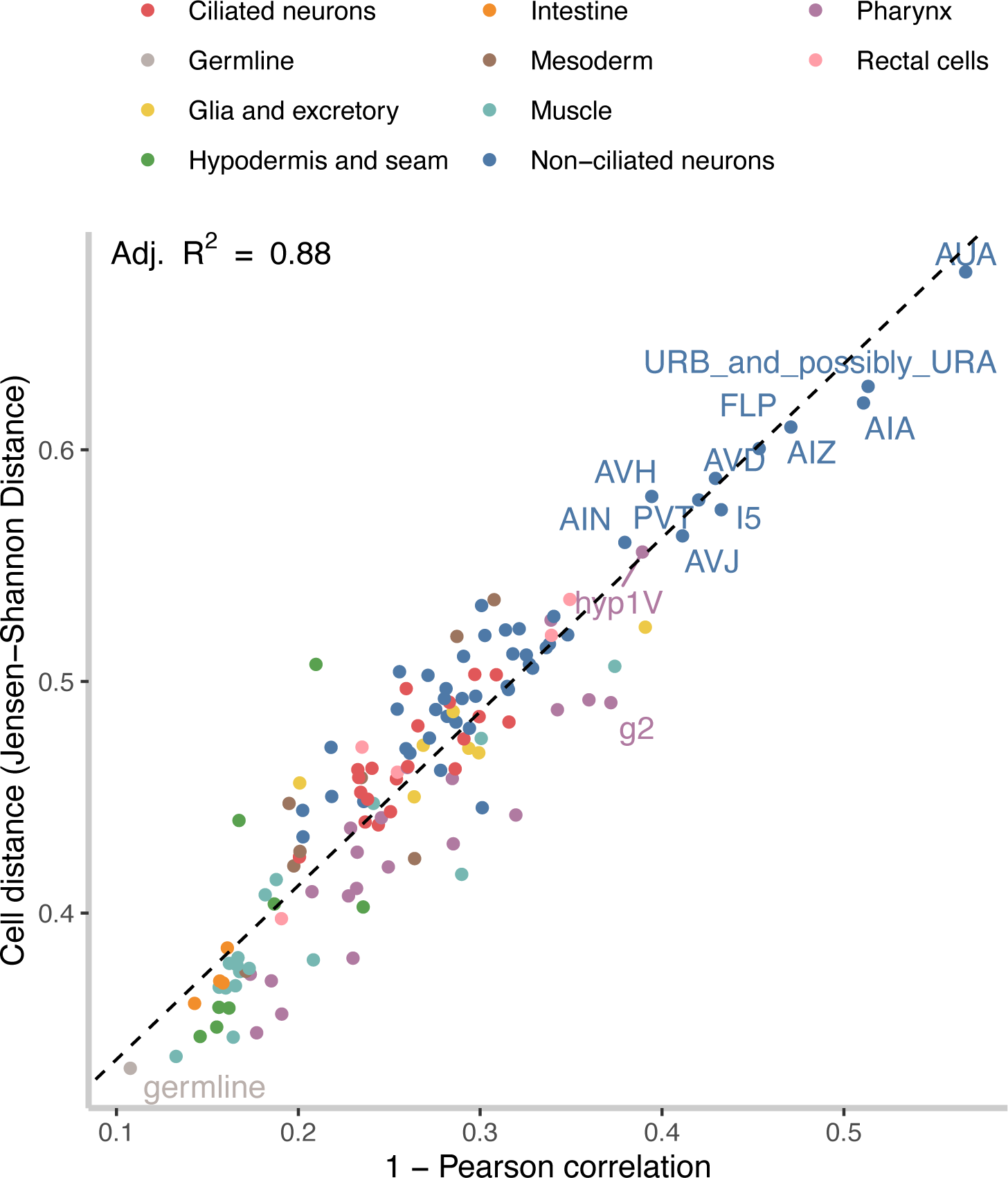
The Pearson correlation and JSD_cell_ show a strong linear relationship indicating the robustness of the cell distance measurement (Adj. R^2^ of 0.88).

**Supp. Figure 33.**
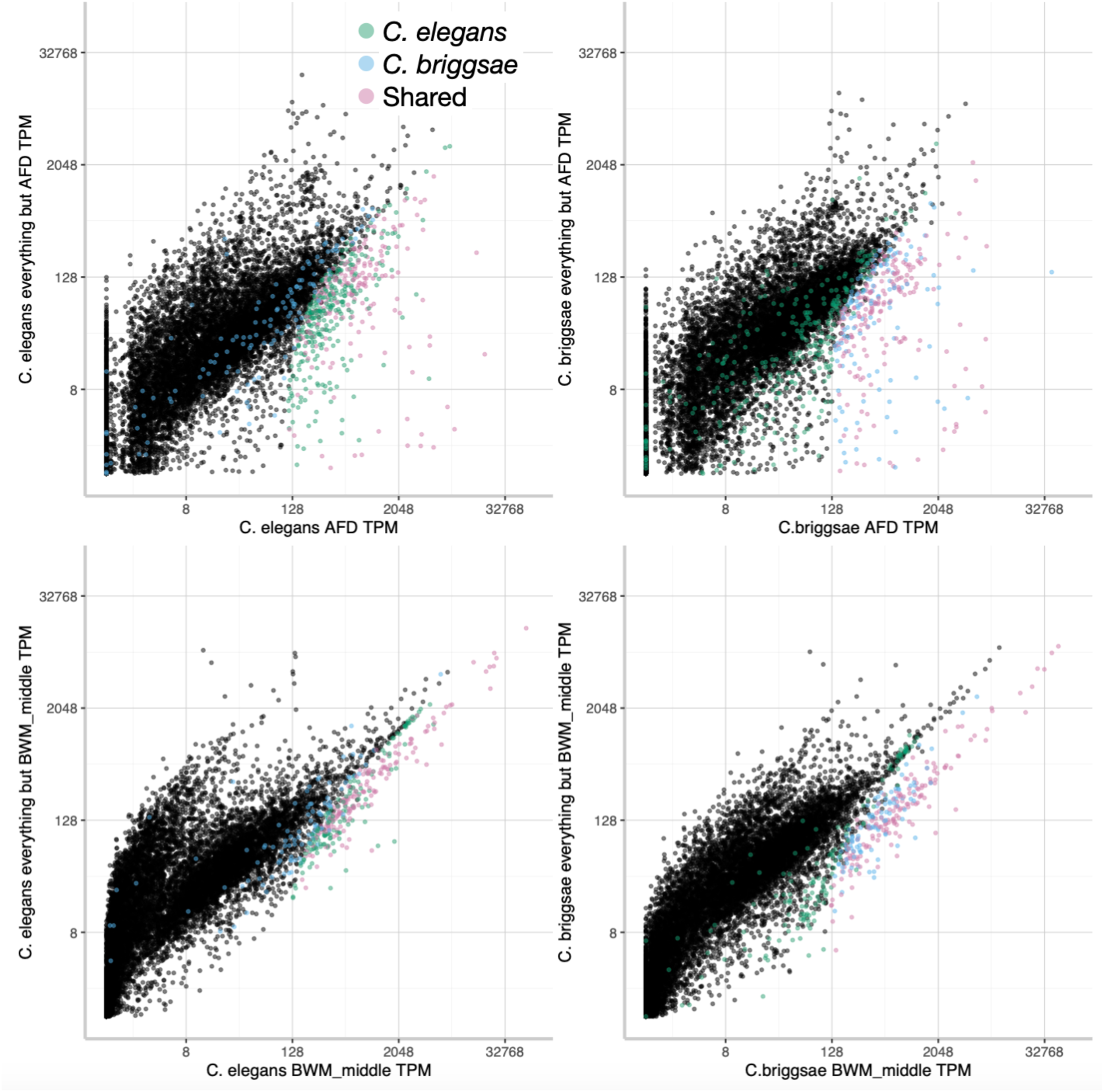
The expression of all 1:1 orthologs in TPM for AFD (top) and BWM_middle (bottom) versus the expression in everywhere but AFD or BWM_middle. The color of the dots indicates whether the gene is a marker of AFD or BWM_middle in *C. elegans*, *C. briggsae*, or both. Generally, genes that are markers in one species, but not the other are either poorly expressed in one species, or expressed more highly in other cell types, and thus losing specificity.

**Supp. Figure 34.**
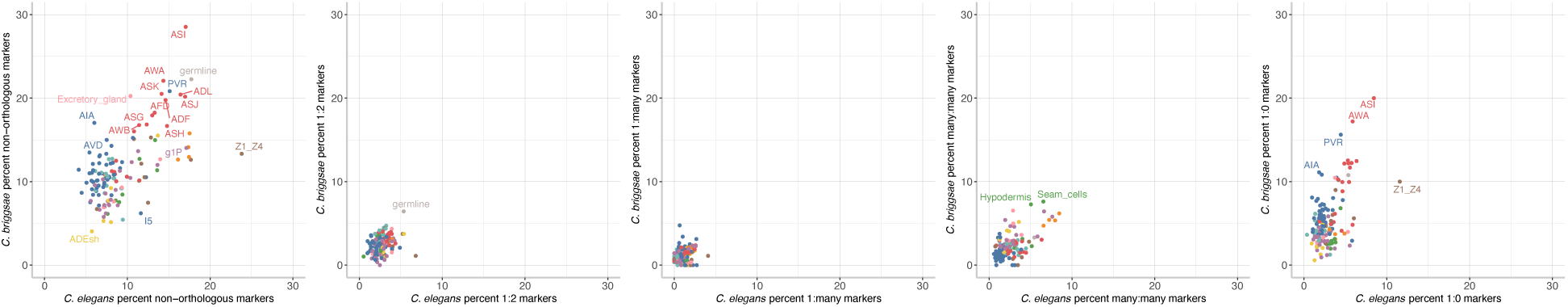
The percentage of total *C. elegans* markers that are in each orthology relationship (1:2, 1:many, many:many, and 1:0) are shown. The colors denote the cell class as outlined in Figure 3.

**Supp. Figure 35.**
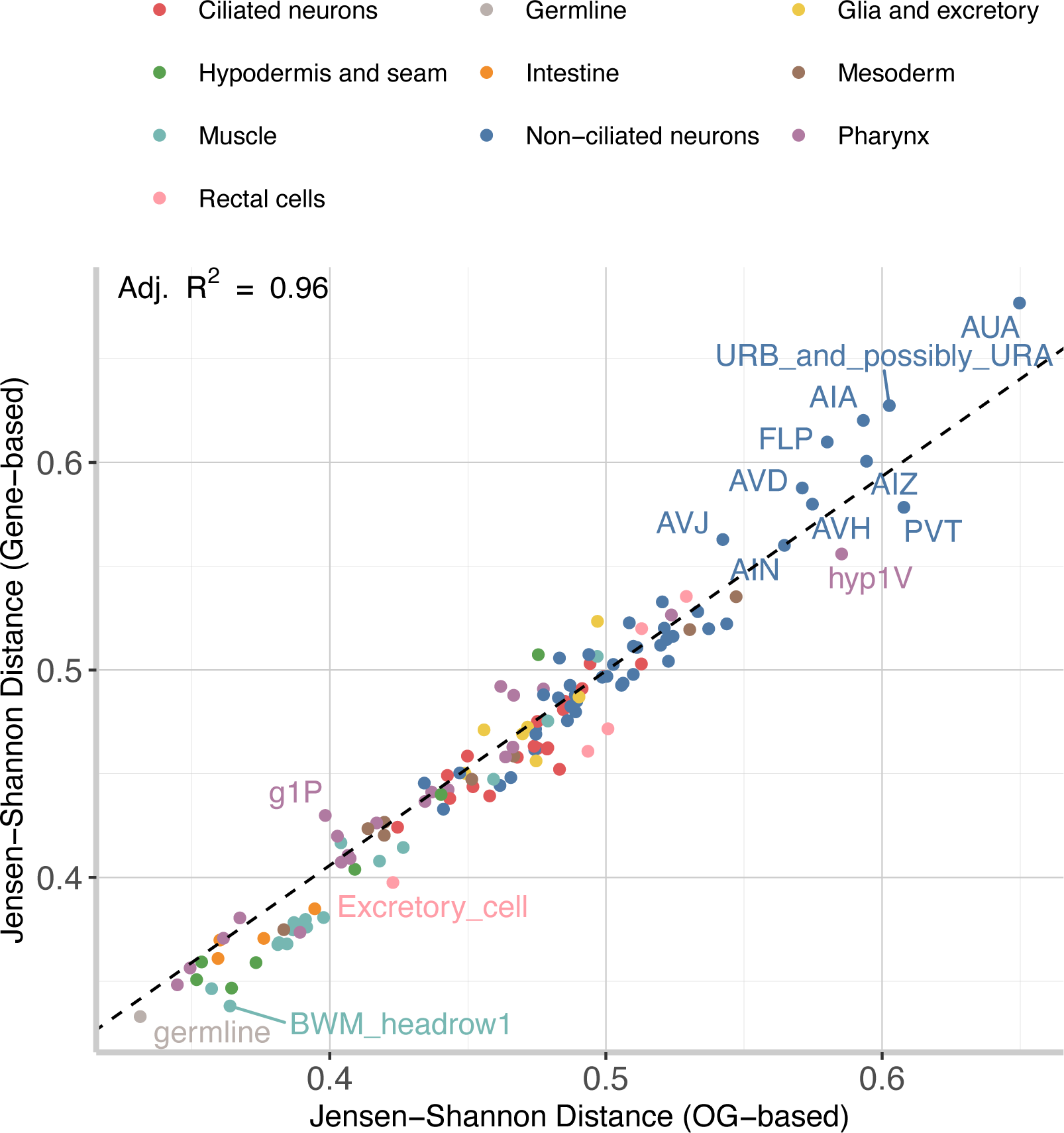
JSD_cell_ was calculated on either the TPM values from the 1:1 orthologs (gene-based) or on the summary TPM values of the orthogroups. There was a strong linear relationship between the two methods of calculating JSD_cell_, indicating that non 1:1 orthologs do not have a strong effect on the cell distance.

**Supp. Figure 36.**
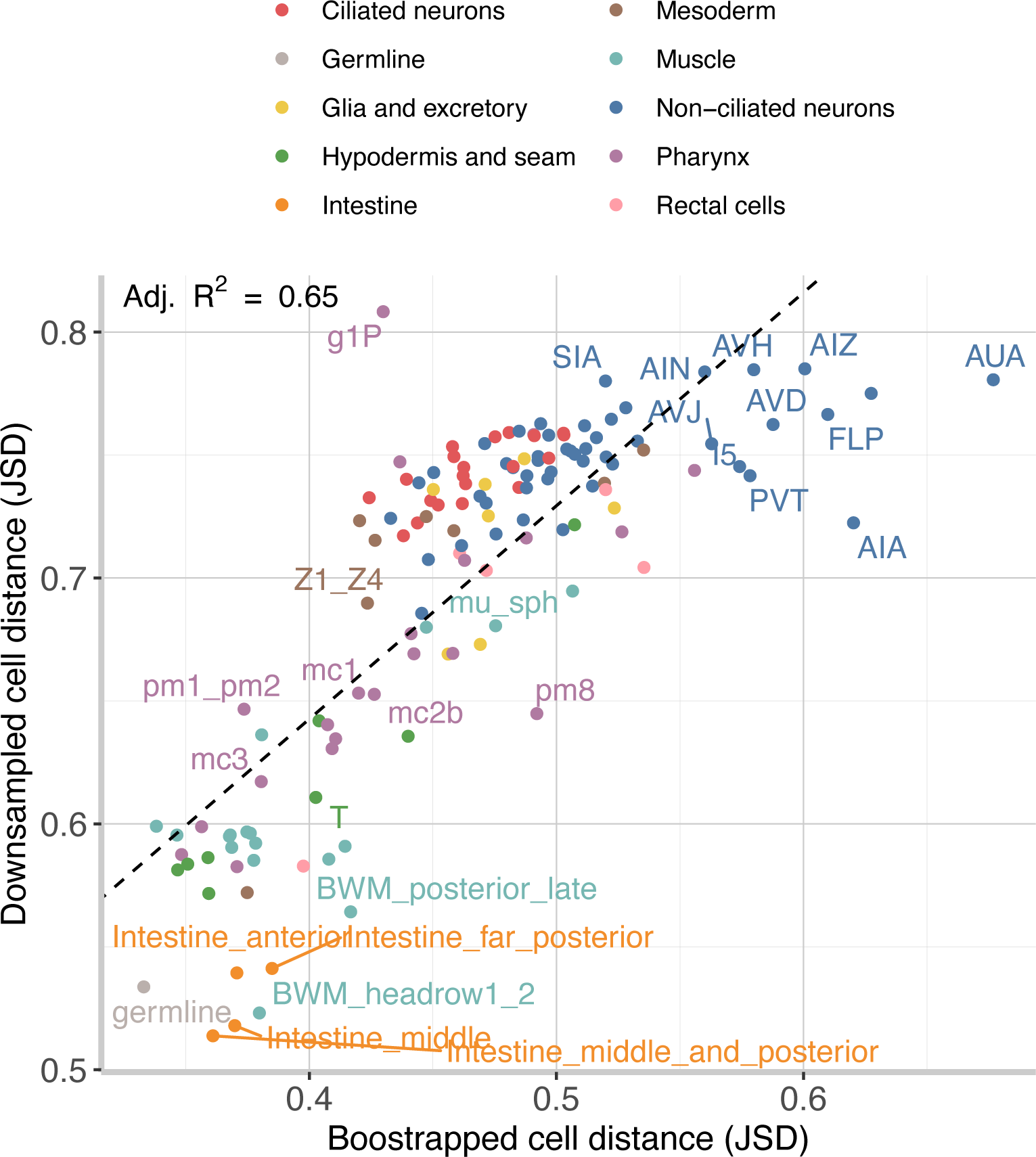
A comparison of the original, bootstrapped JSD_cell_ values for each cell type versus the median JSD_cell_ of 10,000 random downsamples of each cell type to 10 cells each to correct for sampling biases. Overall, there was a good correspondence between the values, indicating that the collected cell count does not have a large influence on the JSD_cell_ calculation. The largest deviations in rank order between the two calculated JSD_cell_ values were either from either a high Gini coefficient cell type (g1P), or from well sampled cell types such as the body wall muscle, intestine, or germline.

**Supp. Figure 37.**
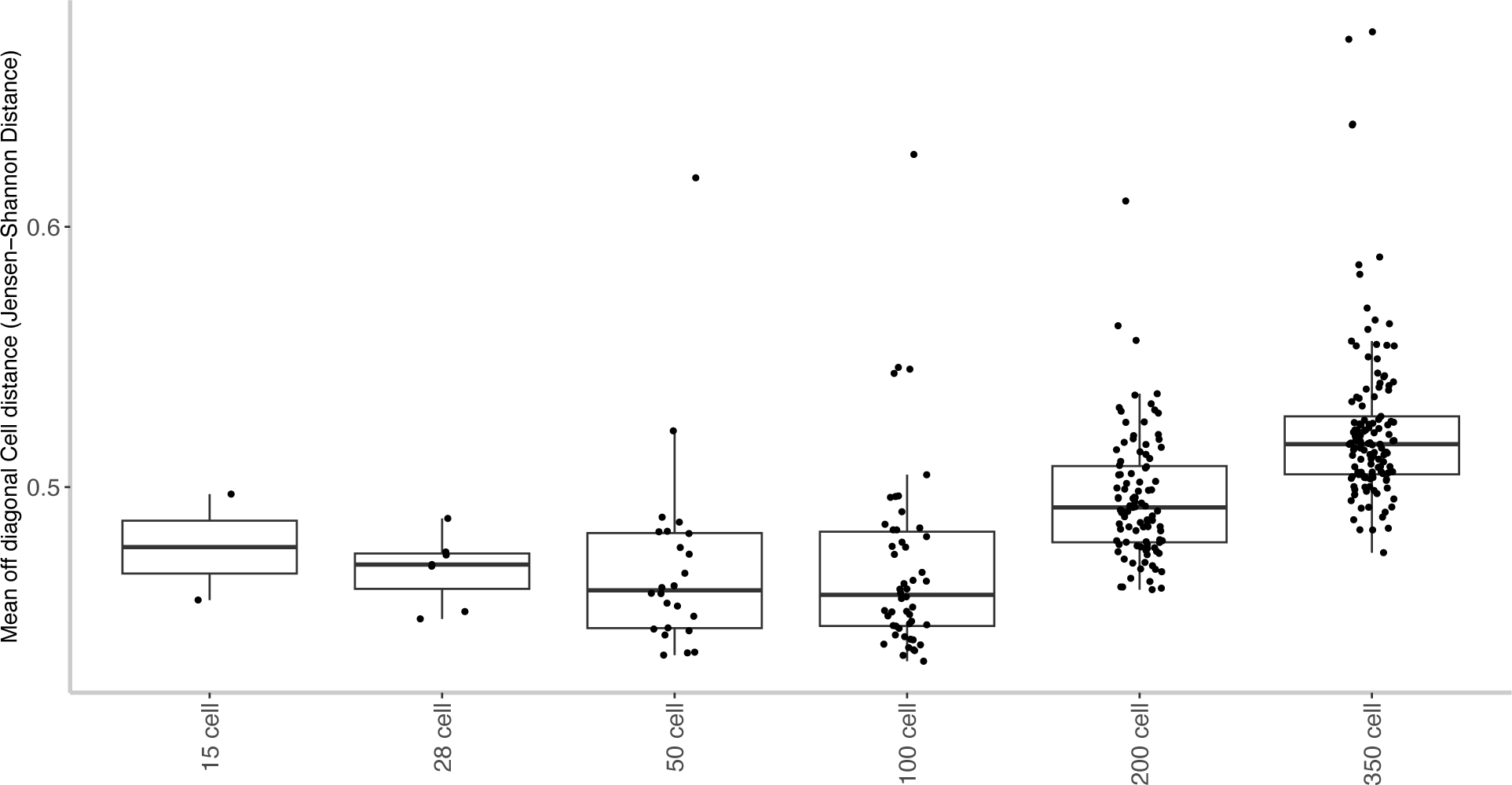
Mean JSD_cell_ value for comparisons between each *C. elegans* progenitor with the non-homologous progenitors from *C. briggsae* within each division stage. Early *C. elegans* progenitors (<= 100 cell) are more similar to early *C. briggsae* non-homologous cells than later cells.

**Supp. Figure 38.**
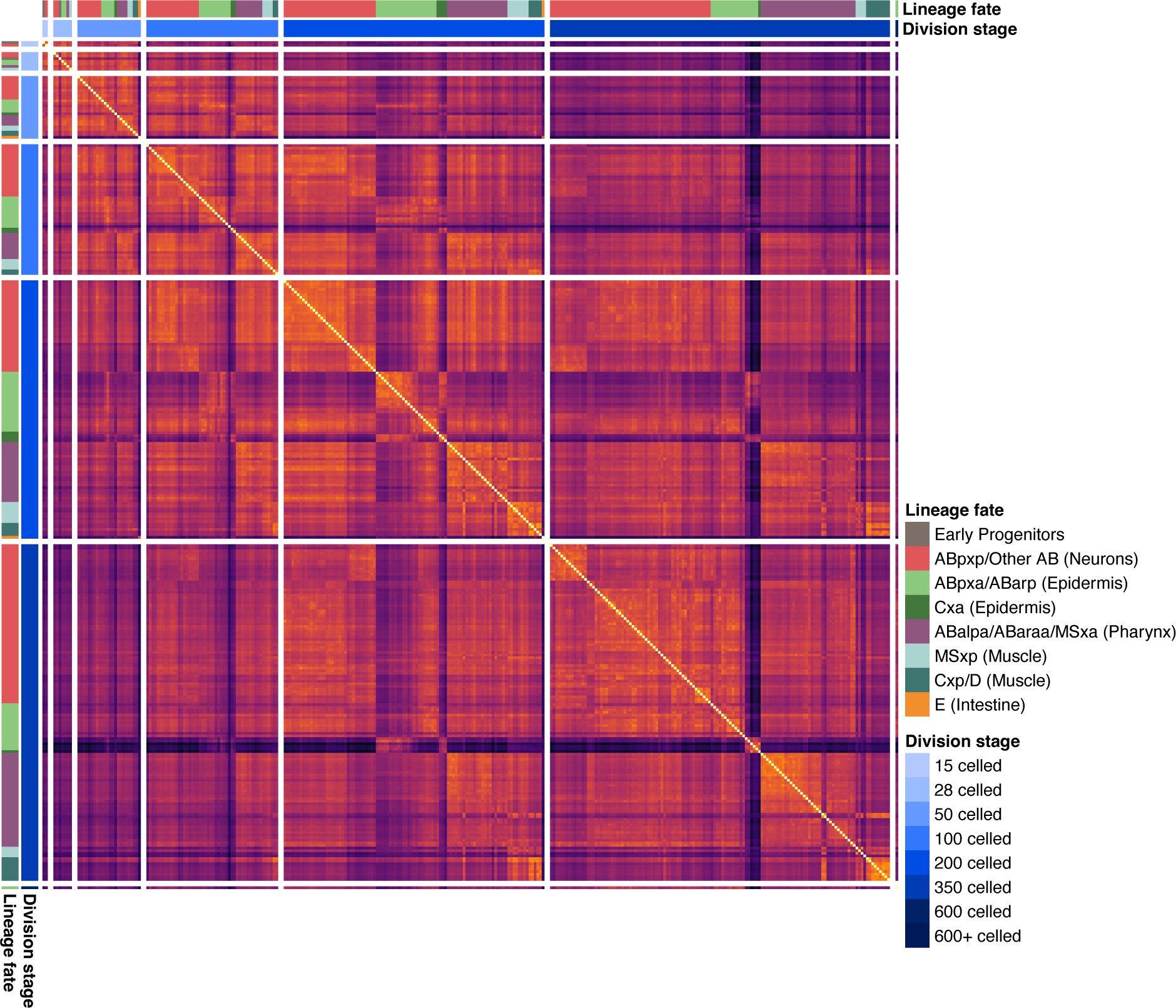
Heatmap illustrating the transcriptome divergences between all *C. elegans* progenitor cells calculated as the Jensen-Shannon Distance (JSD_cell_).

**Supp. Figure 39.**
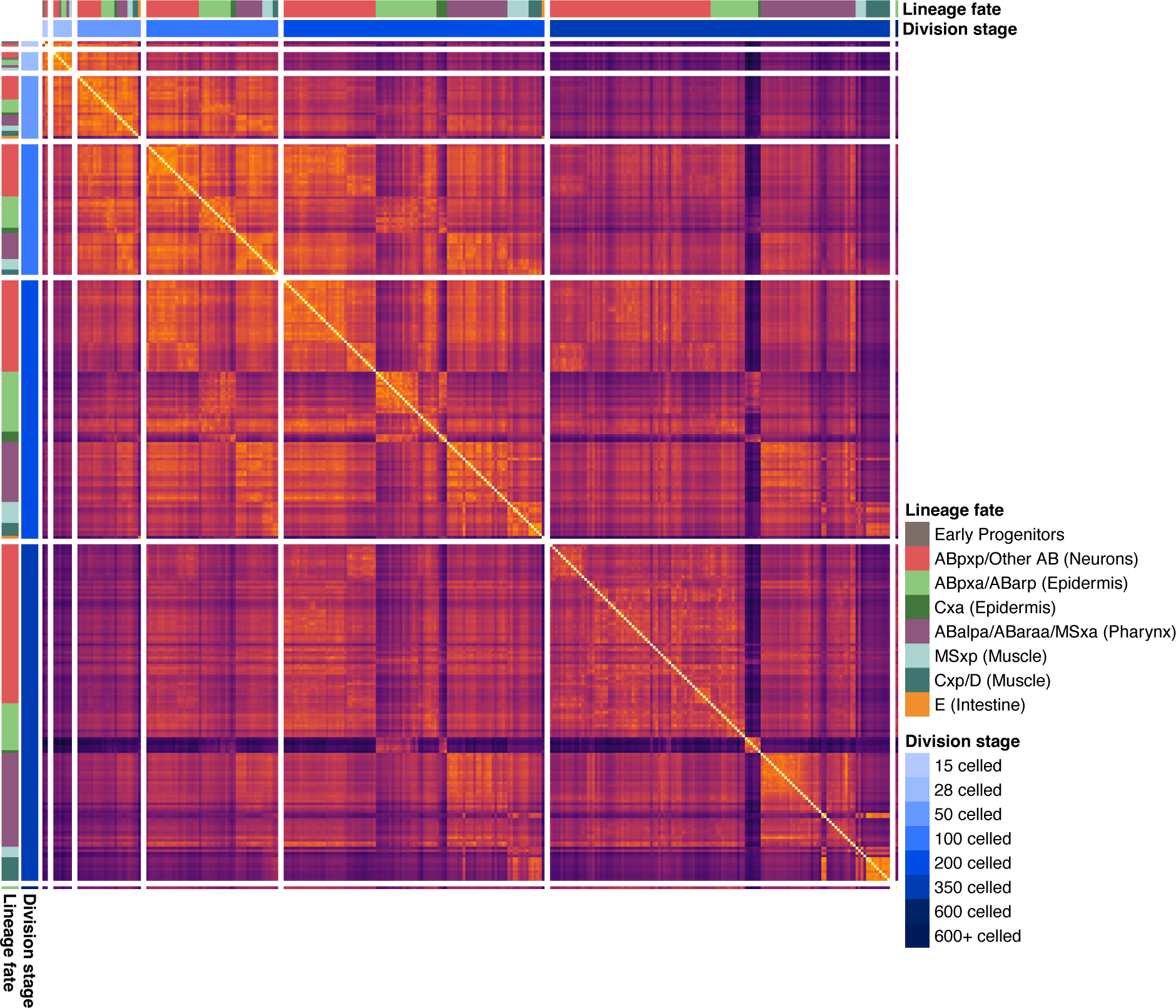
Heatmap illustrating the transcriptome divergences between all *C. briggsae* progenitor cells calculated as the Jensen-Shannon Distance (JSD_cell_).

**Supp. Figure 40.**
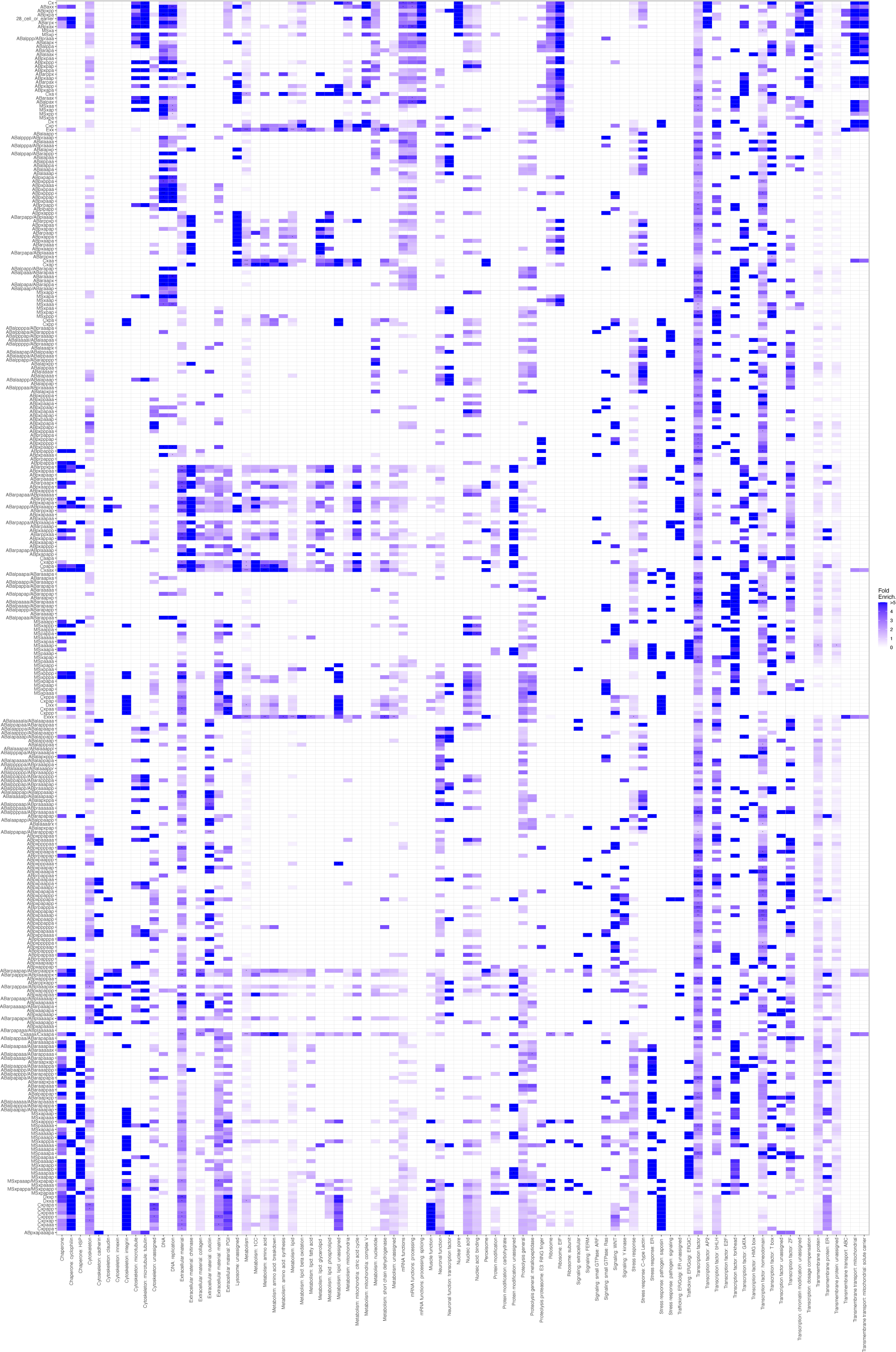
The enrichment of the WormCat category annotations for the shared markers of the progenitor cell types for *C. elegans*. The stars represent Bonferroni corrected p-values of a Fisher’s Exact Test where * is p <0.05, ** is p <0.005, and *** is p <0.0005.

**Supp. Figure 41.**
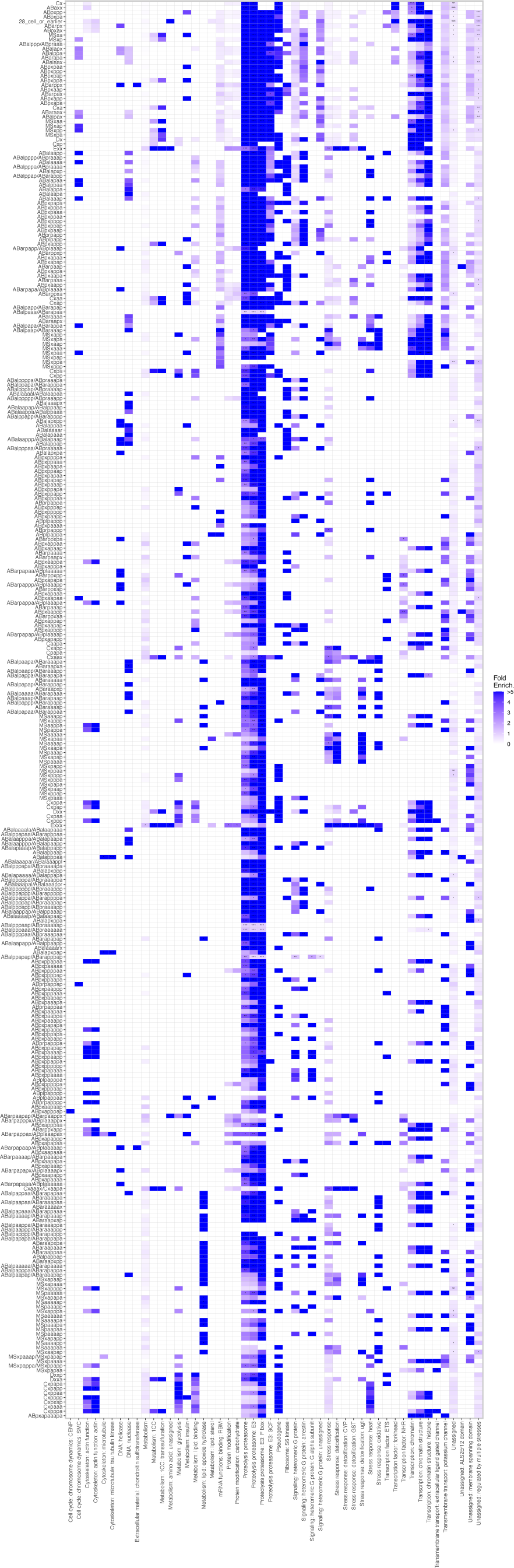
The enrichment of the WormCat category annotations for the non 1:1 markers of the progenitor cell types for *C. elegans*. The stars represent Bonferroni corrected p-values of a Fisher’s Exact Test where * is p <0.05, ** is p <0.005, and *** is p <0.0005.

**Supp. Figure 42.**
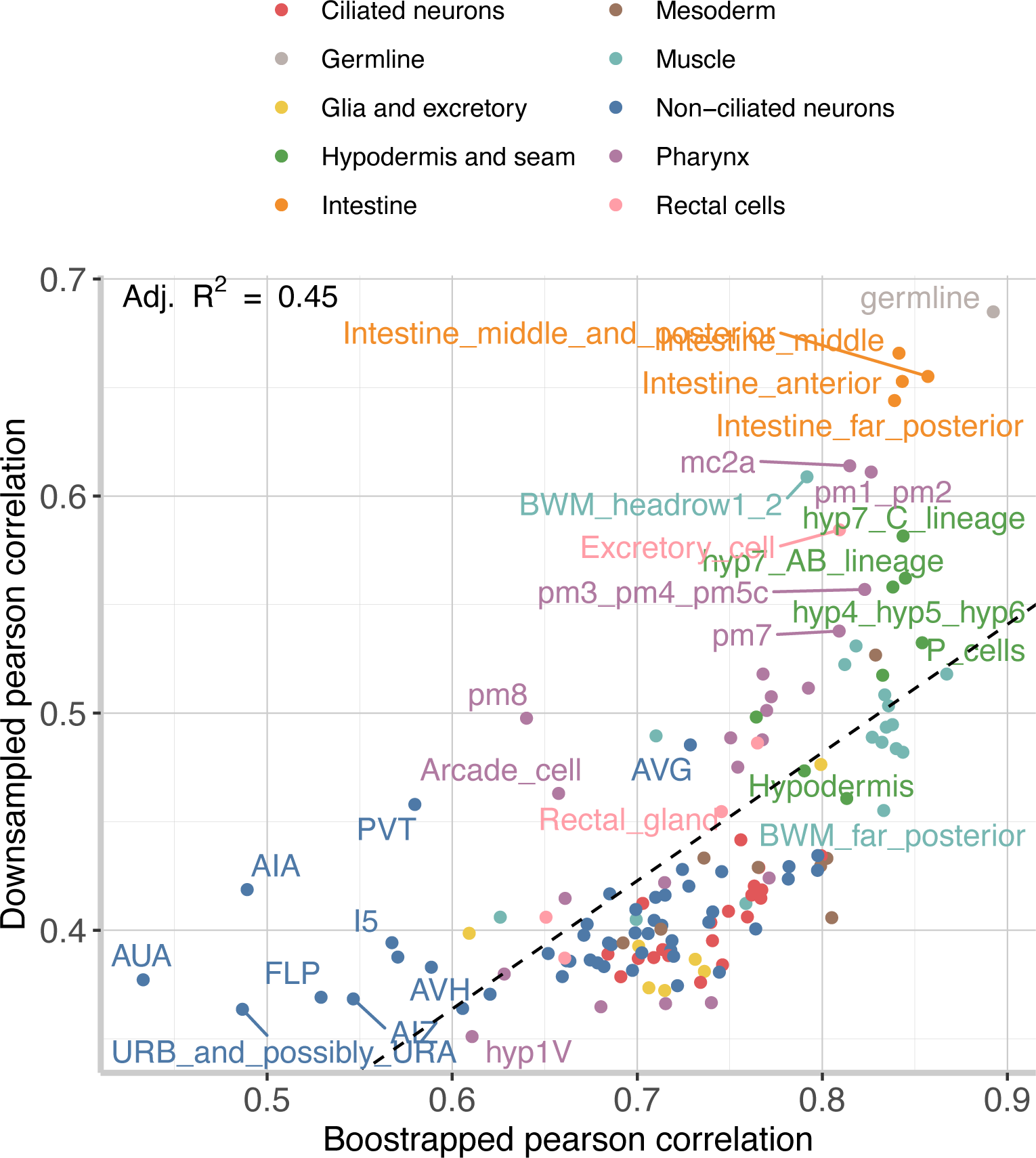
A comparison of the original, bootstrapped cell expression Pearson correlation values for each cell type versus the median Pearson correlation values of 10,000 random downsamples of each cell type to 10 cells each to correct for sampling biases. Overall, there was a worse correspondence between the values for the Pearson correlation versus the JSD_cell_, indicating that the JSD_cell_ is more robust to data quality.

